# A minimal sequence motif drives selective tRNA dihydrouridylation by hDUS2

**DOI:** 10.1101/2023.11.04.565616

**Authors:** Jingwei Ji, Nathan J. Yu, Ralph E. Kleiner

## Abstract

The post-transcriptional reduction of uridine to dihydrouridine (D) by dihydrouridine synthase (DUS) enzymes is among the most ubiquitous transformations in RNA biology. D is found at multiple sites in tRNAs and studies in yeast have proposed that each of the four eukaryotic DUS enzymes modifies a different site, however the molecular basis for this exquisite selectivity is unknown and human DUS enzymes have remained largely uncharacterized. Here we investigate the substrate specificity of human dihydrouridine synthase 2 (hDUS2) using mechanism-based crosslinking with 5-bromouridine (5-BrUrd)-modified oligonucleotide probes and *in vitro* dihydrouridylation assays. We find that hDUS2 modifies U20 in the D loop of diverse tRNA substrates and identify a minimal GU motif within the tRNA tertiary fold required for directing its activity. Further, we use our mechanism-based platform to screen small molecule inhibitors of hDUS2, a potential anti-cancer target. Our work elucidates the principles of substrate modification by a conserved DUS and provides a general platform to studying RNA modifying enzymes with sequence-defined activity-based probes.

## INTRODUCTION

Post-transcriptional modifications on RNA play an important role in biological processes^1^. To date, over 150 modifications have been found on RNA^2^. In particular, transfer RNA (tRNA) is the most extensively modified, containing on average 13 modifications per molecule. Modifications on tRNA affect gene expression at the translational level through diverse mechanisms and many are broadly conserved throughout evolution^3, 4^. Generally, modifications in the anticodon loop regulate codon-anticodon interactions, while modifications in the tRNA body are involved in proper folding and stabilization of tertiary structure. Emerging evidence indicates that tRNA modifications are dynamically regulated and mediate translational programs in response to cell state or external stimuli^5^. Therefore, investigating the molecular mechanisms underlying the activity of RNA modifying enzymes is critical to understanding how RNA modification levels are controlled and regulated in biological systems.

Dihydrouridine (D) is one of the most abundant and highly conserved tRNA modifications^6, 7^. In eukaryotes, D modifications are installed by four dihydrouridine synthase enzymes (DUS) and mainly found in the eponymous tRNA D loop at positions 16/17, 20, and 20a/20b, or at position 47 in the tRNA variable loop. D is non-planar and adopts the C2’ endo conformation^8^, which disfavors its incorporation in double-stranded RNA, and suggests a role in modulating tRNA structure. However, the biological function of D has remained mysterious. Interestingly, D levels are enriched in psychrophilic bacteria^9^ and DUS enzymes are implicated in human cancers^10^, but the underlying mechanisms are unknown.

Dihydrouridylation of tRNAs presents a challenging problem in molecular recognition. In yeast, Phizicky and co-workers evaluated the substrate specificities of the four DUS enzymes using microarray and primer extension analysis to show that these proteins have non-overlapping substrate sites^11^. Despite sharing a conserved active site, three of the four yeast DUS enzymes modify distinct uridine residues within close proximity to one another in the D loop (i.e. Dus1p modifies U16/17, Dus2p modified U20, and Dus4p modifies 20a/20b) (Fig. 1a). How do individual DUS enzymes select the correct substrate sites and reject adjacent uridine residues presented within similar sequence and structural context? Further, work from our lab^12^ and others^13, 14^ has shown that D also exists on messenger RNA (mRNA), indicating that DUS enzymes can recognize and modify diverse RNA substrates.

**Figure 1.**
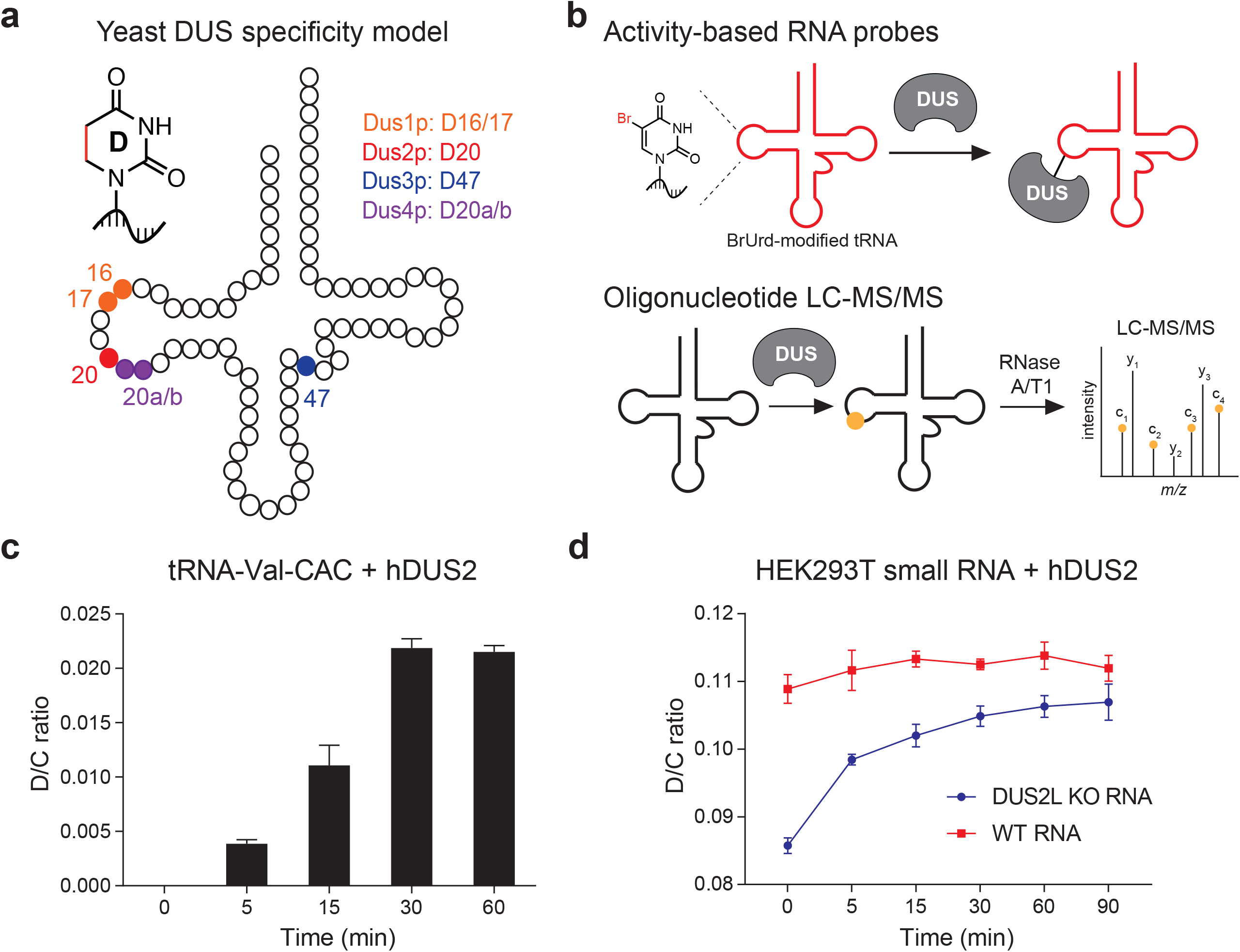
Biochemical investigation of human dihydrouridine synthase 2 (hDUS2). **(a)** Dihydrouridine sites on tRNA and the specificity of the corresponding yeast DUS enzymes. **(b)** Strategy for hDUS2 biochemical study. **(c)** Time course of D formation on IVT tRNA-Val-CAC after enzymatic reaction with recombinant hDUS2. D content was quantified by nucleoside LC-QQQ-MS. Three independent biological replicates were analyzed. Values represent mean ± s.d. (n=3). **(d)** Time course of D formation on bulk small RNA isolated from WT HEK293T or DUS2L KO cells after enzymatic reaction with recombinant hDUS2. D content was quantified by nucleoside LC-QQQ-MS. Three independent biological replicates were analyzed. Values represent mean ± s.d. (n=3).

Structural insights into bacterial DUS-tRNA interactions have revealed that residues U16 and U20, modified by DusC subfamily and DusA subfamily enzymes, respectively, and located on opposite sides of the D loop, are selectively recognized by a major reorientation in binding mode of the tRNA substrate^15, 16^. Whereas yeast and bacterial DUS enzymes have been studied in defined systems *in vitro*^17, 18^, the human DUS homologues remain largely uncharacterized. X-ray crystal structures of individual domains from human DUS2L (hDUS2) have been determined^19,20^ and Hamdane and co-workers characterized binding of the dsRBD domain to tRNA^20, 21^, however they did not report a co-structure of the full-length protein with tRNA or study dihydrouridylation across a range of potential substrates. Consequently, further investigation is needed to elucidate the principles underlying substrate selection by human DUS enzymes.

Previously, we used metabolic RNA labeling with 5-fluorouridine (5-FUrd) to induce mechanism-based crosslinking between human DUS3 (DUS3L) and its cellular RNA substrates, enabling activity-based profiling of DUS3L and transcriptome-wide mapping of its modification sites^12^. Despite the proposed mechanistic similarity among DUS enzymes, we found that 5-FUrd did not react appreciably with human DUS enzymes other than DUS3L. In addition, we did not characterize the nature of the 5-FUrd-DUS3L crosslink nor did we reconstitute DUS3L-RNA crosslinking *in vitro*. Herein, we investigate the *in vitro* substrate specificity of hDUS2, a human DUS enzyme implicated in lung cancer proliferation^10^, using two complementary approaches (Fig. 1b). First, we develop sequence-defined RNA activity-based probes for hDUS2 using RNA oligonucleotides modified with 5-bromouridine (5-BrU) and evaluate mechanism-based crosslinking with a panel of tRNA-like substrates. Next, we study dihydrouridylation of *in vitro* transcribed tRNAs by oligonucleotide LC-MS/MS. Using these two approaches, we establish rules governing the substrate specificity of hDUS2 and evaluate small molecule inhibitors. Taken together, our work provides a general framework for studying RNA modifying enzymes using oligonucleotide-based activity probes and reveals insight into the installation of an abundant tRNA modification implicated in human disease.

## RESULTS

### Recombinant hDUS2 is active in vitro

To investigate the substrate specificity of hDUS2 using mechanism-based crosslinking, we first purified recombinant enzyme from *E. coli* (Supplementary Fig. 1) and confirmed its activity by measuring D formation on an *in vitro* transcribed (IVT) tRNA substrate (Supplementary Table 1) by LC-QQQ-MS (Fig. 1c). We chose human tRNA-Val-CAC since this tRNA has multiple potential D modification sites, including U residues at positions 17, 20, and 20a in the D loop, and 47 in the variable loop (Fig. 1a). Previously, Hamdane and co-workers demonstrated that hDUS2 modifies U20 on bulk tRNA isolated from yeast but were not able to observe activity on an IVT tRNA species^21^. To our knowledge, *in vitro* studies of hDUS2 with human tRNAs have not been reported. Gratifyingly, we detected D formation on IVT tRNA-Val-CAC after only 5 min incubation with recombinant hDUS2 (Fig. 1c) and achieving maximum conversion at 30 min, validating the activity of our bacterially expressed enzyme and also demonstrating that hDUS2 can modify tRNAs lacking endogenous post-transcriptional modifications. The maximum D concentration measured on tRNA-Val-CAC is consistent with on average no more than one D modification per tRNA (i.e. D/C ratio = 0.022 and there are 20 C residues in tRNA-Val-CAC), suggesting a specific substrate site rather than promiscuous tRNA modification. We also found that enzyme activity decreased by ∼50% if we omitted tRNA refolding (Supplementary Fig. 2), indicating the importance of tRNA structure, and was negligible in a catalytically dead hDUS2 C116A mutant (Supplementary Fig. 3). As an additional control for *in vitro* enzyme specificity, we measured D formation on bulk endogenous small RNA (<200 nt) purified from WT HEK293T cells or a matched DUS2L KO cell line generated using CRISPR/Cas9 technology^22^. D levels on bulk small RNA are reduced by 22.7% in DUS2L KO cells and incubation with recombinant hDUS2 resulted in a 19.9% increase in D concentration, effectively restoring D to WT levels (Fig. 1d). In contrast, we observed no activity of hDUS2 on bulk small RNA isolated from WT HEK293T cells. Taken together, our data show that recombinant hDUS2 generated through heterologous expression in *E. coli* is catalytically competent, can install D on unmodified tRNA transcripts, and shows comparable specificity to that of the native protein.

### Mechanism-based crosslinking of hDUS2 with 5-bromouridine (BrU)-modified tRNA

After demonstrating that hDUS2 modifies IVT tRNA-Val-CAC, we chose this tRNA sequence as a starting point to investigate mechanism-based crosslinking with modified RNA. In previous work with human DUS3L (which we showed modifies U47 in the variable loop)^12^, we proposed a mechanism for DUS crosslinking with 5-FUrd-modified RNA that involves nucleophilic attack of the conserved catalytic Cys residue on the C5 position after reduction (Fig. 2a), however we did not directly characterize the adduct, nor demonstrate its formation outside of the cell. Further, whereas 5-FUrd metabolic labeling induces crosslinking between DUS3L and labeled cellular RNA, we did not detect efficient crosslinking with other human DUS enzymes. In contrast, C5-halogenated pyrimidine analogues containing chloro or bromo substitutions can generate crosslinked adducts with all four human DUS enzymes^22^. Therefore, we focused on 5-bromouridine (5-BrUrd) as a mechanism-based probe for hDUS2. To study crosslinking between hDUS2 and BrUrd-modified-RNA, we *in vitro* transcribed tRNA-Val-CAC using 5-BrUrd triphosphate (5-BrUTP) in place of UTP resulting in a tRNA containing BrUrd in place of every U residue (hereafter “BrU-tRNA-Val-CAC”) (Supplementary Fig. 4). Next, we incubated BrU-tRNA-Val-CAC with recombinant hDUS2 in buffer containing NADPH and analyzed protein-RNA crosslinking by anti-hDUS2 western blot. We observed the formation of a protein-RNA crosslink that was confirmed by RNase treatment and was not found with tRNA-Val-CAC containing canonical U residues (Fig. 2b, Supplementary Fig. 5). As expected, we did not observe efficient crosslinking between IVT tRNA-Val-CAC generated with 5-fluoroUTP and hDUS2 (data not shown). The presence of the terminal CCA residues in the acceptor stem, which are added post-transcriptionally, had no effect on crosslinking efficiency (Supplementary Fig. 6). We further measured the efficiency of hDUS2 crosslinking (EC_50_ = 45.66 +/- 8.5 nM) using a dose titration experiment with BrU-tRNA-Val-CAC (Supplementary Fig. 7). Crosslinking yields did not exceed ∼50% despite using up to 75-fold excess of BrU-tRNA-Val-CAC, which is likely due to instability of the enzyme during the reaction but could also reflect inefficiency in the crosslinking chemistry.

**Figure 2.**
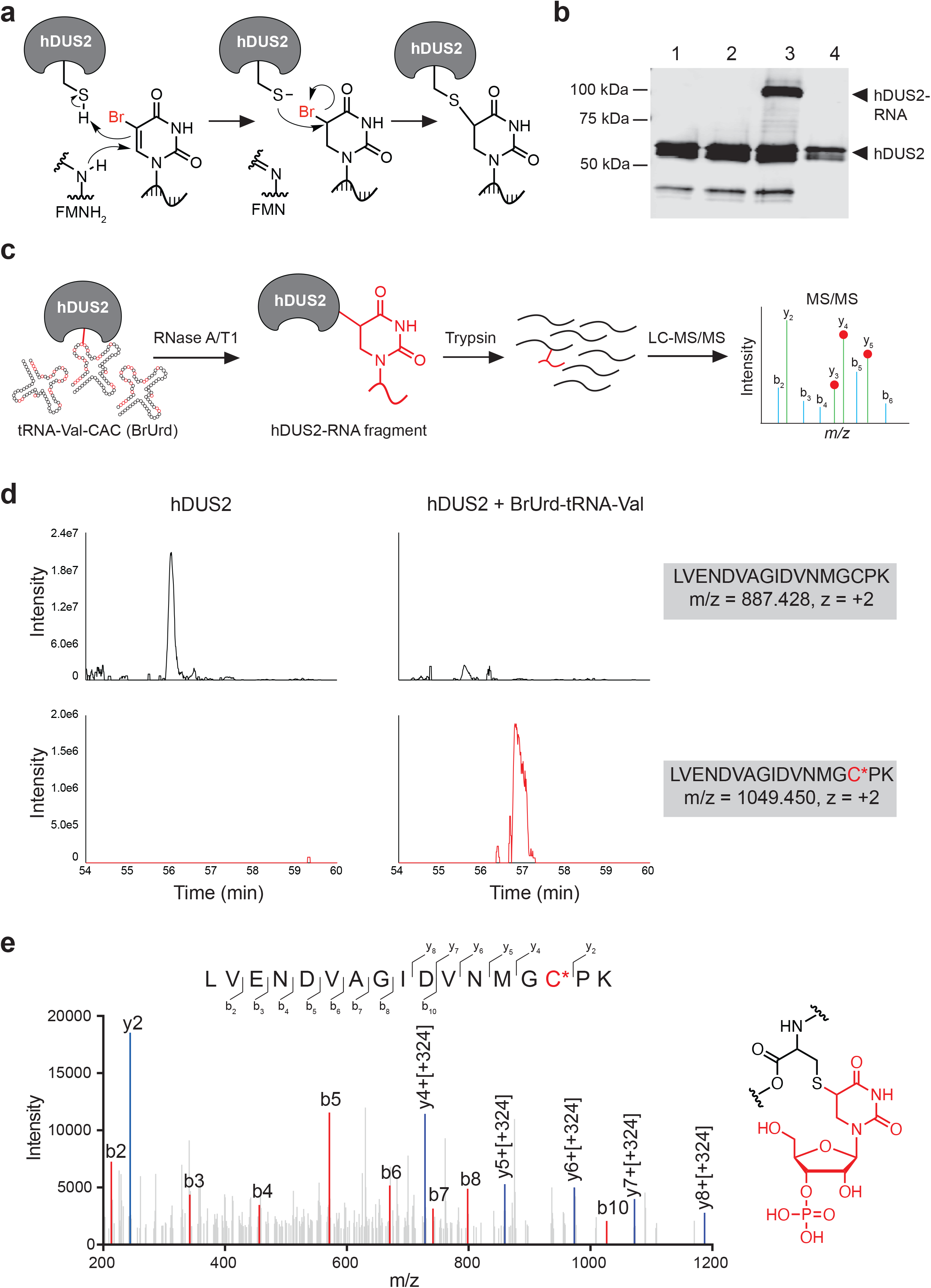
Mechanism-based crosslinking of hDUS2 with 5-BrUrd-modified tRNA. **(a)** Proposed mechanism of crosslinking between hDUS2 and 5-BrUrd-modified RNA. **(b)** Western blot analysis of crosslinking between 5-BrUrd-modified tRNA-Val-CAC and recombinant hDUS2. Lane 1: hDUS2 only; lane 2: hDUS2 with unmodified tRNA-Val-CAC; lane 3: hDUS2 with BrUrd-modified tRNA-Val-CAC; lane 4: RNase treatment of lane 3. Full blot can be found in Supplementary Fig. 5. **(c)** Schematic of bottom-up proteomic workflow to characterize the tRNA-hDUS2 covalent adduct. **(d)** Extracted ion chromatogram of crosslinked peptide-oligo species (bottom) and unmodified peptide fragment (top) in hDUS2 control sample and reaction with BrUrd-modified tRNA-Val-CAC. **(e)** MS/MS analysis of the modified peptide fragment (*m/z* = 1049.450) produced from the reaction of hDUS2 with BrU-tRNA-Val-CAC after digestion with RNase A/T1 and trypsin. The proposed structure of the peptide-nucleotide adduct is shown on the right.

We next characterized the putative covalent adduct formed between hDUS2 and BrU-tRNA-Val-CAC using mass spectrometry. We performed crosslinking (Supplementary Fig. 8), followed by digestion with RNase A, RNase T1, and trypsin, to generate small oligonucleotide-peptide adducts amenable to LC-MS characterization using bottom-up proteomics (Fig. 2c). Due to the higher concentrations of protein and RNA in this scaled-up reaction, we observed ∼90% crosslinking efficiency (Supplementary Fig. 8). According to our proposed crosslinking mechanism, the covalent bond should be formed between a nucleophilic residue in the protein (most likely the conserved catalytic Cys residue) and the C5 position of BrUrd with loss of Br (Fig. 2a). We therefore performed an unbiased search for peptide-oligo adducts consistent with this mechanism allowing for oligonucleotide length up to four residues (due to the challenge of ionizing large oligo-peptide species) and no more than two missed RNaseA/T1 cleavages (Supplementary Table 2), and found a molecular ion with m/z ([M+2H]^2+^) = 1049.450, corresponding to the mass of peptide LVENDVAGIDVNMGCPK (encompassing the catalytic C116 residue) modified with dihydrouridine monophosphate (Fig. 2d). In contrast, we did not observe the 1049.450 ion in tryptic digests of hDUS2 alone, instead finding the unmodified LVENDVAGIDVNMGCPK peptide ([M+2H]^2+^ = 887.428). Similarly, the unmodified LVENDVAGIDVNMGCPK peptide was absent in the crosslinked sample. We confirmed the peptide-nucleotide adduct identity using MS/MS analysis of b and y ions, which unambiguously localized the nucleotide modification to C116 (Fig. 2e). Despite the presence of BrUrd at multiple positions in tRNA-Val-CAC, we were not able to detect other possible peptide-oligonucleotide adducts (Supplementary Table 2), suggesting that crosslinking occurs in a specific manner, or generates multiple species that yield nucleotide adducts with identical mass after digestion. Taken together, our findings demonstrate that hDUS2 efficiently forms a mechanism-based covalent adduct with BrUrd-modified tRNA-Val-CAC involving catalytic Cys116 and provide a sequence-defined RNA probe for studying the activity and substrate specificity of hDUS2 *in vitro*.

### Profiling the substrate specificity of hDUS2 with BrUrd-modified tRNA

With our activity-based *in vitro* crosslinking assay in hand, we next investigated the tRNA substrate specificity of hDUS2. We generated a panel of 11 BrUrd-modified human tRNAs by IVT and evaluated crosslinking to hDUS2 at two different concentrations (1 µM and 10 µM) (Fig. 3a-3d, Supplementary Fig. 9). Among the 11 different tRNAs tested, we identified crosslinked adducts to 9 species. The two tRNAs that did not crosslink were tRNA-Arg-ACG and tRNA-Phe-GAA, which both lack U at position 20 (Supplementary Table 3). Among the 9 tRNAs demonstrating measurable crosslinking behavior, 8 out of 9 contain U20. Crosslinking proceeded with dramatically different efficiency on tRNAs, indicating that specific sequence and/or structural determinants are responsible for modulating recognition and enzymatic modification. Crosslinking to BrUrd-modified tRNA-Glu-TTC and tRNA-Val-CAC proceeded with the highest yield (∼50% crosslinking at 1 µM tRNA), and these tRNAs have similar D loop sequences with U at positions 20 and 20a. Crosslinking to tRNA-Gly-GCC and tRNA-Cys-GCA, which contain U at position 20 but not 20a proceeded less efficiently (25-30% at 1 µM tRNA), and we detected low level crosslinking (4-17% at 1 µM tRNA) to tRNA-Lys-CTT, tRNA-Tyr-GTA, tRNA-Asp-GTC, tRNA-Met-CAT and tRNA-Pro-CGG. Crosslinking efficiency for all tRNAs but tRNA-Val-CAC and tRNA-Glu-TTC increased when tRNA concentration in the reaction was raised from 1 µM to 10 µM.

**Figure 3.**
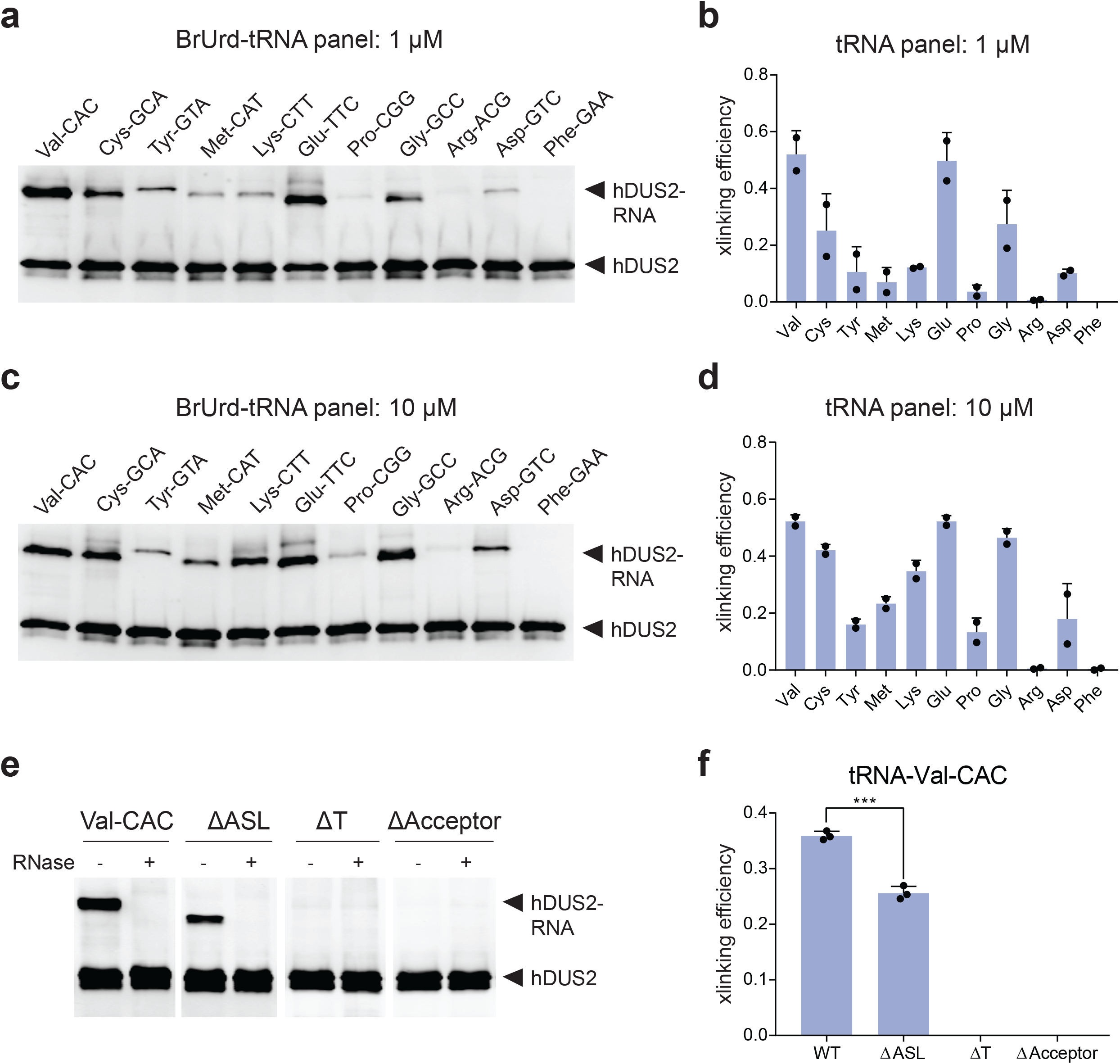
Investigation of hDUS2 specificity using mechanism-based crosslinking with a panel of 5-BrUrd-modified tRNAs. **(a)** Reaction of hDUS2 with BrUrd-modified tRNAs at 1 µM concentration. RNA-protein adduct formation was characterized by anti-hDUS2 western blot. Full blot can be found in Supplementary Fig. 9. Two independent replicates were performed. **(b)** Quantitation of crosslinking from **(a).** Values represent mean ± s.d. **(c)** Reaction of hDUS2 with BrUrd-modified tRNAs at 10 µM. Full blot can be found in Supplementary Fig. 9. Two independent replicates were performed. **(d)** Quantitation of crosslinking from **(c).** Values represent mean ± s.d. **(e)** Crosslinking reaction between hDUS2 and truncated BrUrd-modified tRNA-Val-CAC constructs. Adduct formation was characterized by anti-hDUS2 western blot. Full blot can be found in Supplementary Fig. 14. **(f)** Quantitation of crosslinking from **(e).** Values represent mean ± s.d. Three independent replicates were performed.

Our data and previous observations with yeast Dus2p^11^ support dihydrouridylation and mechanism-based crosslinking at position 20 in the D loop. To validate this finding, we generated tRNA-Val-CAC containing a U20A mutation by IVT and studied crosslinking (Supplementary Fig. 10). No crosslinked band was formed when hDUS2 was incubated with the U20A mutant, supporting this as the site of crosslinking or as a critical residue for enzyme recognition. Interestingly, in our tRNA panel (Fig. 3a-3d) we detected low-level crosslinking to elongator tRNA-Met-CAT, which lacks U at position 20 or 20a but contains U16 in the D loop. To locate the crosslinking site in tRNA-Met, we evaluated two different tRNA-Met isoacceptors (tRNA-Met-CAT-4-1 and tRNA-iMet) that completely lack U residues in the D loop using our crosslinking assay (Supplementary Fig. 11). We observed a similar crosslinking adduct with tRNA-Met-CAT-4-1 but not for tRNA-iMet, suggesting that crosslinking can occur outside of the D loop. We further demonstrated that hDUS2 can install D on tRNA-Met-CAT-2-1 and 4-1 isoacceptors by LC-QQQ-MS analysis, however modification levels were 50-fold lower than observed with tRNA-Val-CAC (Supplementary Fig. 12), and we therefore did not pursue these findings further.

### Molecular determinants of hDUS2-RNA crosslinking

We investigated the structural determinants for hDUS2 crosslinking using tRNA-Val-CAC as our model system. Synthetic 22-mer oligonucleotides mimicking the D-arm of tRNA-Val-CAC containing BrUrd at position 20, 20a, or both (Supplementary Table 4), failed to exhibit crosslinking, even at high concentrations (Supplementary Fig. 13). We next generated truncated BrUrd-modified tRNA-Val-CAC derivatives lacking the T arm, anticodon stem-loop (ASL), or acceptor stem (Supplementary Fig. 14a). Deletion of either the T arm, or acceptor stem of the tRNA completely abolished crosslinking to hDUS2 (Fig. 3e, 3f, Supplementary Fig. 14b). In contrast, we still detected crosslinking to a tRNA lacking the ASL, although crosslinking efficiency was lower than with full-length tRNA. Our data support a model in which hDUS2-tRNA recognition requires the presence of multiple sequence elements including the D arm, T arm, and acceptor stem. While these regions are not proximal to one another in linear sequence, they form close contacts in the L-shaped three-dimensional tRNA structure^23^. Notably, the ASL is not strictly required for hDUS2 modification, which is consistent with the ability of hDUS2 to modify tRNAs with diverse anticodon sequences and also suggests that tRNA lacking the ASL can still fold into a native-like core structure. Hamdane and co-workers proposed a similar model for hDUS2-tRNA recognition based upon NMR and SAXS analysis of the hDUS2 dsRBD domain in complex with tRNA^20^.

DUS proteins are generally composed of only two conserved domains, an N-terminal catalytic domain adopting a TIM barrel fold (TBD) and a unique C-terminal helical domain (HD), whereas human hDUS2 also contains a dsRBD^19,24,25^, which has been proposed to be essential for tRNA modification primarily through interactions with the acceptor stem and the TψC arm^25,20^. Therefore, we used our crosslinking assay to study the importance of the dsRNA binding domain (dsRBD) in hDUS2. We generated a truncated hDUS2 lacking the dsRBD and evaluated crosslinking with the same panel of BrUrd-modified tRNAs assayed above. Whereas we could still detect crosslinking to the most efficient hDUS2 substrates, tRNA-Val-CAC and tRNA-Glu-TTC, albeit at lower efficiency than with full-length hDUS2, we could not observe crosslinks between the DUS domain alone and other BrUrd-modified tRNAs (Supplementary Fig. 15). We therefore conclude that while the dsRBD is not strictly required for tRNA modification, it does play an important role in substrate binding.

### Oligonucleotide LC-MS analysis of hDUS2 substrate specificity

Although our activity-based crosslinking assay provides a facile method to study hDUS2 activity and substrate specificity, we questioned whether BrUrd modification could artificially affect enzyme recognition or catalysis. Further, for tRNAs containing multiple adjacent U residues, such as tRNA-Val-CAC and tRNA-Glu-TTC that contain U at 20 and 20a, identifying the precise site(s) of modification by crosslinking analysis can be challenging. Therefore, to understand whether crosslinking efficiency is reflective of bona fide dihydrouridylation, and to more confidently establish hDUS2 substrate sites, we set up an oligonucleotide LC-MS platform to characterize the site of hDUS2-mediated D formation on unmodified IVT tRNA substrates. In brief, unmodified IVT tRNAs were incubated with hDUS2, digested into small oligo fragments using sequence-specific nucleases (i.e. RNase A or RNase T1), and analyzed by negative mode LC-MS (Fig. 4a). D formation can be detected by a 2 Da increase in the modified oligonucleotide mass and its position within the oligo can be determined by MS/MS fragmentation of the oligo backbone.

**Figure 4.**
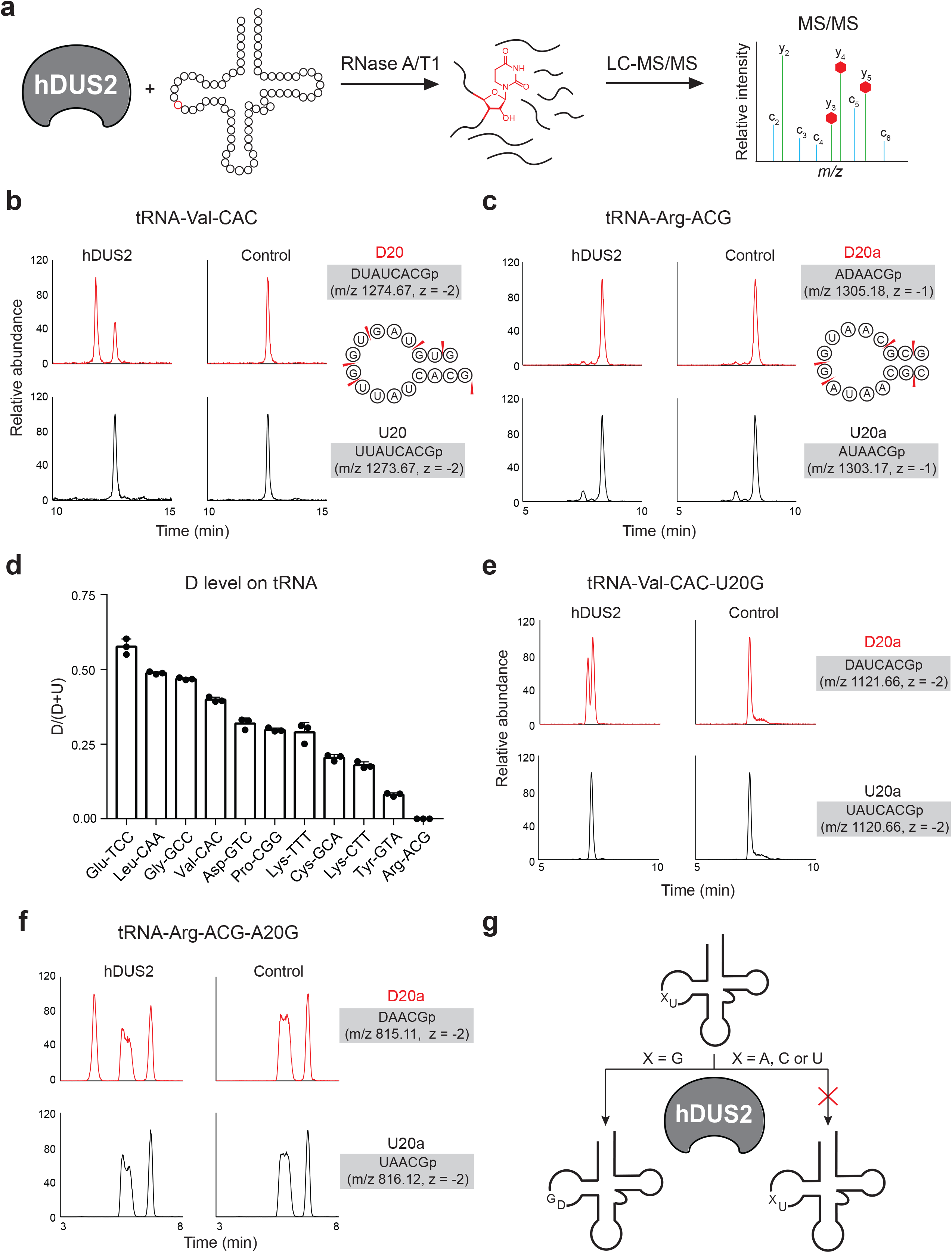
Oligonucleotide LC-MS/MS analysis of tRNA dihydrouridylation by hDUS2. **(a)** Scheme for oligonucleotide LC-MS/MS analysis of hDUS2-catalyzed dihydrouridine formation. **(b)** Extracted ion chromatograms of D-modified and corresponding unmodified oligo fragments produced from the reaction of hDUS2 with tRNA-Val-CAC. The sequence of each oligo with *m/z* value and charge state is indicated on the right. **(c)** Extracted ion chromatograms of D-modified and corresponding unmodified oligo fragments produced from the reaction of hDUS2 with tRNA-Arg-ACG. The sequence of each oligo with *m/z* value and charge state is indicated on the right. **(d)** Measured D formation on IVT tRNAs after hDUS2 reaction. Data was collected from extracted ion chromatograms of corresponding D-modified and unmodified oligos. The D/(D+U) ratio was calculated by comparing the ion intensity of the D-containing oligo against that of the U-containing oligo. Values represent mean ± s.d. (n=3). **(e)** Extracted ion chromatograms of D-modified and corresponding unmodified oligo fragments produced from the reaction of hDUS2 with tRNA-Val-CAC-U20G mutant. The sequence of each oligo with *m/z* value and charge state is indicated on the right. **(f)** Extracted ion chromatograms of D-modified and corresponding unmodified oligo fragments produced from the reaction of hDUS2 with tRNA-Arg-ACG-A20G mutant. The sequence of each oligo with *m/z* value and charge state is indicated on the right. **(g)** Proposed specificity model for hDUS2.

We picked a similar same set of tRNAs used for the crosslinking assay, substituting tRNA-Phe-GAA, which lacks U residues at 20/20a/20b positions, with tRNA-Leu-CAA, a reported substrate for yeast Dus2p^11^. Next, we performed oligonucleotide MS analysis after treatment with hDUS2 and detected D modification on 10 of the 11 selected tRNA substrates (Fig. 4b, 4c, Supplementary Fig. 16-36) – only tRNA-Arg-ACG was not a substrate for hDUS2. In all cases, we only detected one D modification site per tRNA, which mapped to position 20 in the D loop (Supplementary Fig. 16-36). This was the case even for tRNA-Glu-TTC and tRNA-Val-CAC, which contain adjacent U residues at positions 20 and 20a (Supplementary Fig. 17, 23). We quantified D stoichiometry by comparing the abundance of the corresponding D-modified and unmodified oligo using MS1 ion intensity (Fig. 4d, Supplementary Fig. 16-36, Supplementary Table 5) and found that dihydrouridylation efficiency correlated closely with crosslinking efficiency (Fig. 3d) – indeed tRNA-Glu-TTC, tRNA-Val-CAC, and tRNA-Gly-GCC were the top substrates in both assays. In addition, tRNA-Leu-CAA was modified efficiently by hDUS2 but was not studied using the BrUrd-based crosslinking assay. Similar to our crosslinking-based study, we analyzed whether the terminal 3’ CCA affected the reaction and did not observe significant differences (Supplementary Fig. 37-41), indicating that hDUS2 does not recognize these nucleotides.

How does hDUS2 select a single U residue among the multiple possible U substrates within the D loop? In particular, how does it differentiate between adjacent U residues on the same side of the D loop (i.e. 20/20a/20b) as we observed for tRNA-Val-CAC, tRNA-Glu-TTC, and tRNA-Leu-CAA. The positions preceding U20 in tRNAs are almost invariably G18 and G19, therefore we investigated whether these preceding residues were important in mediating hDUS2 modification at U20. To test this hypothesis explicitly, we generated mutant tRNA-Val-CAC constructs containing U20C or U20G substitutions and evaluated D formation by hDUS2 using our oligo LC-MS assay. D formation was abolished on the U20C substrate (Supplementary Fig. 42), indicating that 20a cannot be modified even in the absence of a competing U substrate at position 20. Surprisingly, we detected D formation at 20a in the U20G construct, suggesting that the G residue introduced through mutation was directing D formation at a new site (Fig. 4e, Supplementary Fig. 43, 44). Similarly, we generated a tRNA-Arg-ACG construct containing an A20G substitution. While native tRNA-Arg is not modified by hDUS2 (Fig. 4c, Supplementary Fig. 36), installation of a G residue in place of A directly upstream of U at 20a serves to direct D formation at this position in the mutant tRNA (Fig. 4f, Supplementary Fig. 45-46). Finally, since many of the hDUS2 tRNA substrates that we identified contain two G residues upstream of the modification site, we tested whether both were required on tRNA-Val-CAC. Evaluation of tRNA-Val-CAC-G18A mutant clearly demonstrated that only a single G residue is required immediately upstream of the modification site (Supplementary Fig. 47, 48). Taken together, we show that hDUS2 selectively installs D at position 20 and our study identifies the necessary and sufficient D loop sequence element that directs modification – namely, a single G residue 5’ to the modification site (Fig. 4g).

### Screening hDUS2 inhibitors using oligonucleotide-based activity probes

Human DUS2 has been implicated as a cancer target based upon its overexpression in non-small cell lung carcinomas (NSCLC), therefore identifying small molecule inhibitors for hDUS2 can have therapeutic value^10^. We envisioned that our mechanism-based crosslinking assay with BrUrd-modified tRNA could provide a useful method to screen direct inhibitors of hDUS2 activity (Fig. 5a), particularly since no high-throughput DUS activity assays have been described. Previously, hDUS2 was identified as an off-target of the acrylamide-containing EGFR inhibitor PF-6274484^26^. To test if this compound can inhibit hDUS2 activity *in vitro*, we pretreated the protein with PF-6274484 (Fig. 5b) and performed activity-based crosslinking with BrU-tRNA-Val-CAC. Our data shows inhibition of hDUS2-tRNA crosslinking at 100uM but not lower concentrations (Fig. 5c, Supplementary Fig. 49), confirming the feasibility of identify DUS inhibitors using this assay. Next, we picked a series of commercially available EGFR covalent inhibitors (Fig. 5b) that all share the 4-amino-quinazoline scaffold and acrylamide electrophile designed to target cysteine^27^. We found that in addition to PD168393, AST1306 and Canertinib also show inhibition of hDUS2 activity (Fig. 5c, Supplementary Fig. 50) at 100 μM and AST1306 is the most potent in this series. Surprisingly, Afatinib was found to modestly activate hDUS2. From this small screen, we conclude that compounds based on the 4-amino-quinazoline scaffold containing unsubstituted acrylamides are promising lead candidates to develop inhibitors of hDUS2. Finally, to investigate the generality of our crosslinking-based assay, we overexpressed human DUS1L and DUS3L proteins in HEK293T cells and incubated lysate with BrU-tRNA-Val-CAC. Our results show efficient crosslinking to DUS1L and DUS3L (Supplementary Fig. 51) and demonstrate how BrUrd-modified tRNAs can be used as general activity-based probes for the human DUS family.

**Figure 5.**
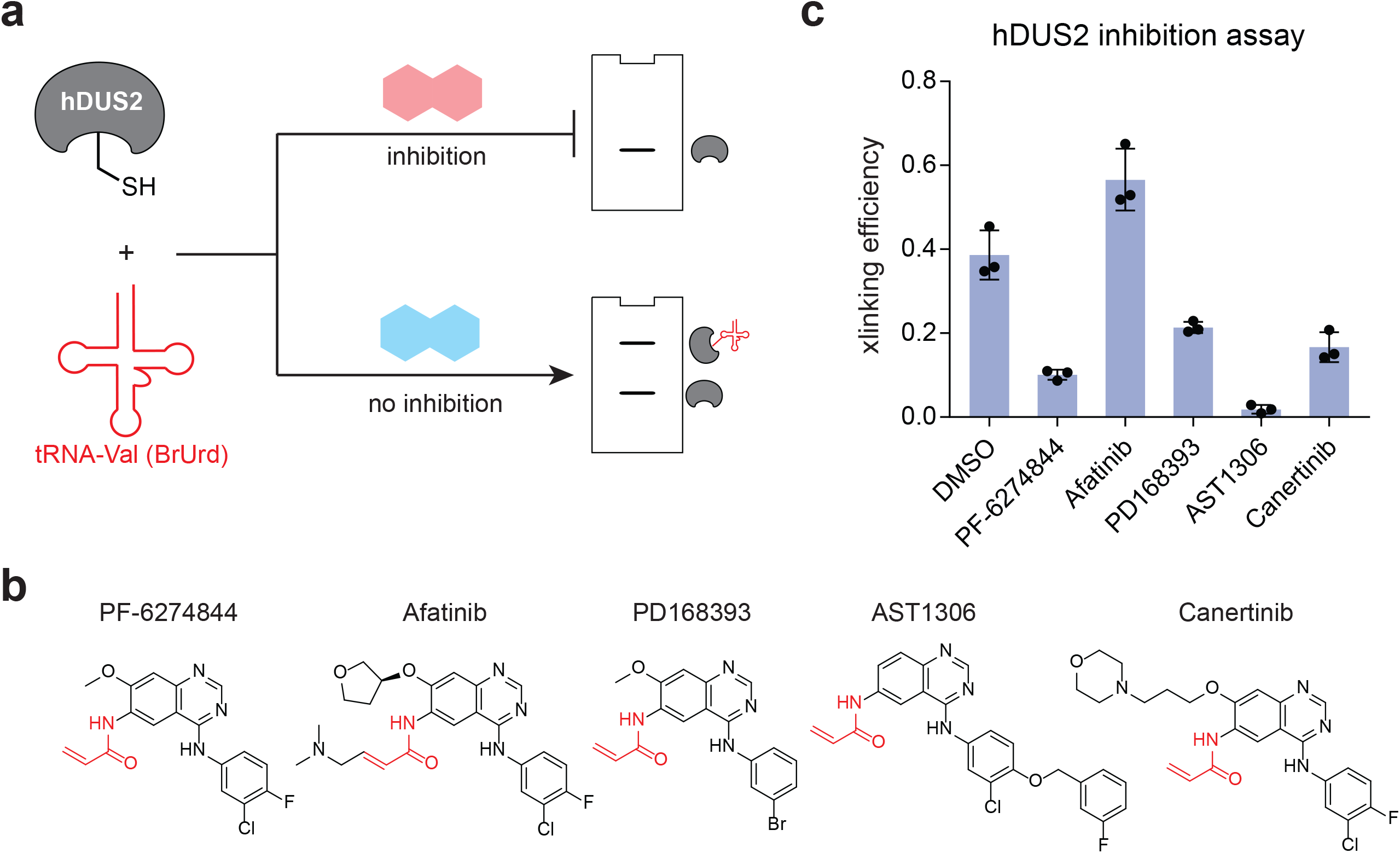
Screening small molecule hDUS2 inhibitors using activity-based oligo probes. **(a)** Scheme for small molecule inhibition assay. **(b)** Chemical structures of inhibitor compounds investigated herein. **(c)** Crosslinking efficiency between hDUS2 and BrU-tRNA-Val-CAC after pre-treatment of hDUS2 with indicated small molecule inhibitor. Crosslinking blots can be found in Supplementary Fig. 50. Values represent mean ± s.d. (n=3).

## DISCUSSION

In this manuscript, we investigate the substrate specificity of hDUS2 using mechanism-based crosslinking and oligonucleotide LC-MS/MS. DUS enzymes and dihydrouridine are conserved throughout evolution, but we lack an understanding of the molecular determinants for substrate recognition and modification by these enzymes. Here we develop 5-BrUrd-modified tRNA activity probes and characterize their mechanism-based crosslinking with recombinant hDUS2. Combined with *in vitro* dihydrouridylation assays and oligonucleotide LC-MS/MS analysis, we survey a panel of putative tRNA substrates and establish that a minimal GU motif in the tRNA D loop directs selective enzyme modification. Further, we apply our activity probing strategy to screen a small panel of acrylamide-based small molecule inhibitors of hDUS2. Taken together, our work reveals molecular insights into hDUS2-mediated tRNA dihydrouridylation and provides a general approach for studying RNA modifying enzymes with defined oligonucleotide-based activity probes.

Using a panel of IVT human tRNA substrates, we show that hDUS2 specifically installs D at position 20 in the D loop. Indeed, among the 12 tRNAs that we investigated, all 10 that contain U at this position were modified by hDUS2, albeit at different efficiencies. The ability of this enzyme to modify diverse tRNAs at a specific site suggests a universal principle for tRNA substrate selection – our work demonstrates that a preceding 5’ G residue is responsible for directing D formation in the D loop, and G19 is invariant among eukaryotic tRNAs explaining the abundance of the D20 modification. We could not recapitulate D formation in minimal stem-loop oligonucleotides nor in truncated tRNAs (with the exception of a tRNA lacking in the ASL), indicating the importance of tRNA tertiary structure in hDUS2 recognition, as demonstrated by Hamdane and co-workers in their studies of the isolated dsRBD^20^. In contrast to their findings, however, we show that unmodified IVT tRNAs are substrates for hDUS2, suggesting that the installation of D20 can occur early in the tRNA modification circuit. What is the role of the preceding G residue in directing modification at U20? G19 is known to Watson-Crick base pair with C56 in the T loop^23^, forming a conserved tertiary interaction at the tip of the tRNA “elbow”. It is likely that direct recognition of the G19:C56 pair in the context of this structure is responsible for setting the register of modification. Interestingly, mutational insertion of G at position 20 in either tRNA-Val-CAC or tRNA-Arg-ACG enables modification by hDUS2 at position 20a. One possible explanation for this behavior is that G20 pairs with C56 in place of G19 leaving the overall elbow structure unperturbed. Further exploration of hDUS2-tRNA recognition, including understanding the role of G19 and catalytic preference among different tRNA substrates, will undoubtedly benefit from structural analysis of hDUS2-tRNA catalytic complexes, and our mechanism-based oligonucleotides probes should enable covalent trapping of these intermediates. Similarly, our strategy should enable biochemical characterization of other DUS enzymes, as the catalytic mechanisms are thought to be broadly conserved.

This work is inspired by our RNABPP^12,28^ strategy that relies upon metabolic labeling of RNA with modified nucleosides. Whereas metabolic labeling can facilitate the study of native RNA-enzyme interactions in cells, there are several advantages to the application of sequence-defined oligonucleotide probes. First, the selection of modified nucleotide is not constrained by the biosynthetic machinery. Second, sequence-defined probes can be applied in the test tube to study individual proteins, screen small molecule inhibitors, and trap enzyme-substrate complexes for structural analysis. Finally, these constructs could be applied to target and inhibit specific RNA modifying enzymes in contrast to metabolically incorporated probes that act transcriptome-wide. DUS enzymes, including hDUS2, have been implicated in cancer^10^ and the design of oligonucleotide-based inhibitors and activity-based screening platforms for these proteins has applications in therapeutic development.

## ACKNOWLEDGEMENTS

We thank John Eng for assistance with oligonucleotide LC-MS/MS analysis. R.E.K. acknowledges support from the NIH (R01 GM132189), NSF (MCB-1942565), and the Alfred P. Sloan Foundation. J.J. and N.J.Y. were supported by a generous gift from the Edward C. Taylor 3rd Year Graduate Fellowship in Chemistry. All authors acknowledge financial support from Princeton University.

