## Supplementary Information for "A minimal sequence motif drives selective tRNA dihydrouridylation by hDUS2"

Table of contents

i. Materials and Methods 2
ii. Supplementary Tables 10
iii. Supplementary Figures 15

### Materials and Methods

*Plasmid construction, protein expression and purification*

The cDNA for hDUS2 was purchased from Dharmacon. The full coding sequence and truncated version (DUS domain 1-333) were PCR amplified, digested, and ligated into the NdeI/NotI sites of a modified pET28a vector, with a $His_6$ tag at the N terminus. Mutagenesis of Cys116 to Ala was made by overlap extension PCR. $His_6$-hDUS2 (full-length, DUS domain, and C116A mutant) was expressed in *Escherichia coli* strain BL21(DE3) for 20 hours at 17 °C with 1 mM IPTG[1]. Cells were lysed by sonication and purified using Cobalt-NTA affinity resin (Thermo Fisher) according to the manufacturer's recommendations. The most concentrated fractions were loaded onto a HiTrap Q column (GE Healthcare) and eluted with NaCl gradient (20 mM Tris pH 8.0, 50 mM - 1M NaCl, 5 mM DTT over 30 column volumes). $His_6$-hDUS2 (full-length and C116A mutant) was further purified by gel filtration using a Superdex 200 Increase 10/300 GL (GE Healthcare) column with the following buffer: 20 mM Tris pH 8.0, 100 mM NaCl, 5 mM DTT. The resulting protein was concentrated to 1 mg/ml with an Amicon centrifugal filter (Millipore).

*In vitro transcription of RNA substrates*

The sequences of human cytosolic tRNAs were obtained from GtRNAdb (http://gtrnadb.ucsc.edu/index.html). DNA templates for WT and mutant tRNAs were generated by primer extension with Klenow fragment (3'->5' exo-) (NEB) using two overlapping synthetic DNA oligonucleotides (sequences are listed in Supplementary Table 1). The first annealing step was performed in 1x rCutSmart buffer (NEB) by incubating at 94 °C for 5 mins and slowly cooling down to 65 °C for 5 min. Annealed primers were then extended by Klenow fragment at 37 °C for 1 hour to make the dsDNA template. The unmodified RNA substrates were transcribed using T7 RNA polymerase (generated according to literature precedent[2]) at 37 °C for 2 hours in a 200 µl of reaction containing 10 µg of dsDNA template, 50 mM Tris-HCl, pH 7.5, 15 mM

$MgCl_2$, 5 mM dithiothreitol, 2 mM spermidine, and 2 mM NTPs. BrU-modified tRNA was generated using the same conditions, except that 5-bromouridine triphosphate (5-BrUTP, Jena Bioscience) was substituted for UTP in the reaction. Reactions were purified by 8% denaturing polyacrylamide gel and recovered via passive elution and ethanol precipitation. The RNA concentration was determined by UV absorbance at 260 nm.

*Endogenous small RNA isolation*

Cells were cultured at 37 °C in a humidified atmosphere with 5% $CO_2$ in DMEM (Life Technologies) supplemented with 10% fetal bovine serum (Atlanta), 1x penicillin-streptomycin (Life Technologies) and 2 mM L-glutamine (Life Technologies). Total RNA was extracted using TRIzol reagent (Thermo Fisher, 15596018) from WT HEK 293T and matched DUS2L knockout cells generated by CRISPR/Cas9 methods[3]. Small RNA was isolated using Zymo RNA Clean & Concentrator-5 (Zymo Research, R1016) following the manufacturer's protocol.

*In vitro dihydrouridylation assays with recombinant hDUS2*

Transfer RNAs were refolded by heating at 65 °C for 5 min and slow cooling to 25 °C at a rate of 0.1 °C/sec in RNase free water. 2 μM tRNA was incubated with 200 nM purified $hDUS2^{WT}$ or $hDUS2^{C116A}$ in 20 mM Tris-HCl pH 8.0, 100 mM NaCl, 5 mM $MgCl_2$, 2 mM DTT, 10 mM NADPH for 1 hour at 37 °C. RNA substrates were extracted into the aqueous phase using TRIzol LS reagent (Thermo Fisher, 10296010), and further purified by Zymo RNA Clean & Concentrator-5 (Zymo Research, R1016) following the manufacturer's protocol.

*Nucleoside liquid chromatography-tandem mass spectrometry*

LC-QQQ-MS sample preparation was performed as previously described[4]. RNA (300-600 ng) was digested with nuclease P1 (Wako, 145-08221) in 20 μl of buffer (7 mM NaOAc, 0.4 mM $ZnCl_2$, pH 5.2) at 37 °C for 2 hours. Then sample was dephosphorylated with Antarctic phosphatase (NEB, M0289) before LC-QQQ-MS analysis. Chromatography was performed on a Thermo Scientific Hypersil GOLD aQ column (3 μm, 150 x 2.1 mm) at 36 °C using a 7 min isocratic method with water (containing 0.1 % formic acid) at a flow rate of 0.4 mL/min. Dynamic multiple reaction monitoring (DMRM) was used for analysis of modified and canonical nucleosides on an Agilent 1260 Infinity II HPLC coupled to an Agilent 6470 triple quadrupole mass spectrometer in positive ion mode. The following source parameters were used for the mass spectrometry: gas temperature 175 °C, sheath gas flow 12 L/min, capillary voltage 2,500V. MS1 (parent ion) to MS2 (deglycosylated base ion) transition for each nucleoside was set as follows: m/z 247 → 115 for D, m/z 268 → 136 for A, m/z 244 → 112 for C, m/z 284 → 152 for G, and m/z 245 → 113 for U. Commercial ribonucleosides were used to generate standard curves. D levels were normalized to the concentration of C in the sample.

*Oligonucleotide liquid chromatography-tandem mass spectrometry*

Reactions containing IVT tRNA and hDUS2 were extracted by acid-phenol:chloroform, pH 4.5 (ThermoFisher, AM9720), followed by purification on Zymo RNA Clean and Concentrator columns. Purified tRNA was then site-specific digested to oligonucleotides following literature precedent.[5] For RNase T1 digestion, purified RNA (10-20 μg) was incubated with RNase T1 (ThermoFisher, 50 units T1/ μg of RNA) in 100 μl of buffer 220 mM $NH_4OAc$, pH 7.0 for 2 hours at 37 °C. For RNase A digestion, purified RNA (10-20 μg) was incubated with RNase A (Sigma, 70856, 0.1 U/ μg of RNA) and dephosphorylated with QuickCIP (NEB, M0525S, 5 U/ μg of RNA) by incubation in 100 μl of buffer 220 mM $NH_4OAc$, pH 7.0 at 37 °C for 2 hours. The digestion mixture was then lyophilized and reconstituted in mobile phase A (10 mM HFIP and 8.6 mM TEA). Digested oligonucleotides were separated on a Poroshell 120 EC-C18 (2.1 x 50 mm, 1.9 μm, Agilent) at a flow rate of 250 μl/min and eluted using a gradient from 0% B (ACN) to 5% B over 18 min. Tandem MS analysis was performed on an Agilent 6545 XT LC-Q-TOF system in negative mode. The ionization source working parameters were as follows: capillary voltage 3.5 kV; gas temperature 275°C, drying gas flow rate 12 l/min, the sheath gas temperature 350°C and sheath gas flow 12 l/min. Each analysis segment contains a full scan from m/z 120-3200 at a fixed acquisition rate of one spectra per second. Detected RNA fragments were compared against the theoretical oligo fragment ions produced by RNase digestion. Detected RNA fragments from the hDUS2-treated samples were further compared against those from tRNA samples without treatment with hDUS2 enzyme. Unique fragments were selected for CID under 3 collision energy channels (35, 45, 55 V) for fragmentation. Manual annotation of MS/MS spectra was performed by calculating expected m/z values of RNase digested fragments based on the parent tRNA sequences using the MongoOligo online calculator.

*RNA oligonucleotides synthesis*

Site specific BrU-modified 22 mer oligos were chemically synthesized on an ABI 394 oligonucleotide synthesizer (Applied Biosystems) using UltraMild synthesis methods (Glen Research). After resin cleavage and full deprotection, the crude synthesis mixture was purified by reverse-phase HPLC with a Zorbax Eclipse XDB-C18 semiprep column on an Agilent 1260 Infinity instrument, and purity was tested with an Infinity Poroshell 120 EC-C18 analytical column. A gradient of 5 to 20% acetonitrile in 0.1 M triethylammonium acetate over 25 minutes was used. The purified fractions were combined, lyophilized and characterized by high-resolution mass spectrometry (HRMS) on an Agilent 6220 Accurate-Mass Time-of-Flight LC/MS (ESI-TOF) in negative mode.

*Gel-based crosslinking assay*

1 μM or 10 μM BrU-modified tRNA substrates were refolded by heating at 65 °C for 5 min and slow cooling to 25 °C at a rate of 0.1 °C/sec in water. 1 μM IVT tRNA was incubated with 200 nM recombinant hDUS2 in 20 mM Tris-HCl pH 8.0, 100 mM NaCl, 5 mM $MgCl_2$, 2 mM DTT, 10

mM NADPH in a total volume of 20 µl for 1 hour at 37 °C and quenched with the addition of 10 µL of 3x SDS sample buffer. The crosslinked adduct and free protein were separated on a 10% SDS-PAGE gel followed by western blot analysis with recombinant anti-DUS2L antibody (Abcam, ab181262). The intensity of the crosslinked RNA-protein species and non-crosslinked protein was quantified by densitometry analysis using ImageJ. Cross-linking efficiency was calculated with the following formula:

$$crosslinking\ efficiency = \frac{crosslinking\ adduct}{(crosslinking\ adduct + non - crosslinked\ protein)}$$

For dose titration experiments, the binding curve was fitted based on a 4-parameter dose-response equation using GraphPad Prism:

$$Y = Bottom + \left(X^{Hillslope}\right) * (Top - Bottom)/(X^{Hillslope} + EC30^{Hillslope})$$

Three technical replicates were analyzed.

For crosslinking in cellular lysate, 3xFlag-tagged hDUS1 and hDUS3 proteins were overexpressed in HEK293T cells by transient transfection. Cells were lysed in 10 mM HEPES pH 7.5, 200 mM NaCl, 1% Triton, 10 mM $MgCl_2$, 1 mM DTT, and protease inhibitor (cOmplete™, Mini, EDTA-free Protease Inhibitor Cocktail, 11836170001) added freshly, and adjusted to 1 mg/ml. Next, 10 µM tRNA was incubated with 45 µg of cell lysate and 1 ul of RNase Inhibitor, Murine (NEB, M0314S) in 50 µl at 37 °C for 1 hour. Control samples were then treated with 2 µl of RNase A/T1 Mix (ThermoFisher) at 37 °C for 30 min. The reaction was quenched with 3X sample buffer and heated to 95 °C for 5min. Samples were resolved by SDS-PAGE and analyzed by western blot with anti-Flag M2 antibody (Sigma, F1804).

*Mass spectrometry-based proteomics*

2 µg of recombinant hDUS2 was incubated with 7.5 µg of BrU-modified tRNA-Val-CAC in 20 mM Tris-HCl pH 8.0, 100 mM NaCl, 5 mM $MgCl_2$, 2 mM DTT, 10 mM NADPH in a total volume

of 50 µl at 37 °C for 1 hour. Next, 5 µl of the reaction was used to confirm crosslinking by gel-based analysis and the rest of the sample was treated with 2 µl of RNase A/T1 Mix (ThermoFisher) at 37 °C for 30 min. The samples were adjusted to 200 µl with 50 mM Ammonium Bicarbonate pH 8.0, and reduced with 5 mM TCEP at 60 °C for 10 min, followed by alkylation with 15 mM chloroacetamide in the dark at room temperature for 30 min. 50 ng of Trypsin Gold (Promega) was added to each sample and incubated end-over-end at 37 °C for 16 hours. The digested sample was acidified by adding TFA to 0.2% final concentration, and were desalted using SDB stage-tips.[6] Digested samples were dried completely in a SpeedVac and resuspended with 21 µl of 0.1% formic acid pH 3. Samples were loaded onto 45 C18-AQ (45 cm x 75 µm x 1.9 µm, Dr. Maisch, Germany), and separated by an Easy-nLC 1200 UPLC system, followed by analysis with an Orbitrap Fusion Lumos (Thermo Scientific, USA). Germany) mated to metal emitter in-line with an Orbitrap Fusion Lumos (Thermo Scientific, USA). The column temperature was set at 50 °C and a one-hour gradient method with a flow rate of 300 nl/min was used. The mass spectrometer was operated in data-dependent acquisition mode with 120,000 resolution MS1 scan (positive mode, profile data type, AGC 4e5, Max IT 54ms, 375-1500 m/z) in the Orbitrap followed by HCD fragmentation in the ion trap with 35% collision energy. A dynamic exclusion list was invoked to exclude previously fragmented peptides for 60 s and a maximum cycle time of 3 s was used. Peptides were isolated for fragmentation using the quadrupole (1.2 Da window). The ion-trap was operated in rapid mode with AGC 1e4, maximum IT of 54 ms and minimum of 5000 ions.

*Data analysis of Mass spectrometry-based proteomics*

Raw files were searched using Sequest HT and MS Amanda algorithms within the Proteome Discoverer 2.5 suite (Thermo Scientific, USA).[7] 10 ppm MS1 and 0.4 Da MS2 mass tolerances were specified. Carbamidomethylation of cysteine was used as a fixed modification, and oxidation of methionine, acetylation of protein N-termini and oligo adducts in Supplementary

Table 2 were set as dynamic modifications. Trypsin digestion with maximum of 2 missed cleavages were allowed. Data was searched against human database downloaded from UniProt.org. Scaffold (version Scaffold_5.1, Proteome Software Inc., Portland, OR) was used to validate MS/MS based peptide and protein identifications. Peptide identifications were accepted if they could be established at greater than 90.0% probability by the Scaffold Local FDR algorithm. Protein identifications were accepted if they could be established at greater than 99% probability and contained at least 2 identified peptides.[8]

*Small-molecule screening assay*

222 nM recombinant hDUS2 was pre-treated with 111 µM small molecule inhibitor in a total volume of 18 µl of reaction buffer (20 mM µl Tris-HCl pH 8.0, 100 mM NaCl, 5 mM $MgCl_2$, 2 mM DTT, 10 mM NADPH) on ice for 30 min. Then 2 µl of refolded BrU-modified tRNA-Val-CAC was added to 1 µM final concentration. The mixture was incubated at 37 °C for 1 hour and quenched by boiling with 3X sample buffer at 95 °C for 5 min. Anti-hDUS2 western blot was performed for crosslinking efficiency evaluation. Crosslinking adduct formation was measured by densitometry by drawing an identically sized box around each band and measuring the raw integrated density minus background. Crosslinking efficiency was calculated as described above.

| tRNA | | Sequence |
|---|---|---|
| Val-CAC | Fwd | AAGCTTAATACGACTCACTATAGTTTCCGTAGTGTAGTGGTTATCACGTTCGCCTCACA |
| | Rev | TGTTTCCGCCCGGTTTCGAACCGGGGACCTTTCGCGTGTGAGGCGAACGTGATAACCACTA |
| Val-G17A | Fwd | AAGCTTAATACGACTCACTATAGTTTCCGTAGTGTAGTAGTTATCACGTTCGCCTCACA |
| | Rev | TGTTTCCGCCCGGTTTCGAACCGGGGACCTTTCGCGTGTGAGGCGAACGTGATAACTACT |
| Val-U19C | Fwd | AAGCTTAATACGACTCACTATAGTTTCCGTAGTGTAGTGGCTATCACGTTCGCCTCACA |
| | Rev | TGTTTCCGCCCGGTTTCGAACCGGGGACCTTTCGCGTGTGAGGCGAACGTGATAGCCACTA |
| Val-U19A | Fwd | AAGCTTAATACGACTCACTATAGTTTCCGTAGTGTAGTGGATATCACGTTCGCCTCACA |
| | Rev | TGTTTCCGCCCGGTTTCGAACCGGGGACCTTTCGCGTGTGAGGCGAACGTGATATCCACT |
| Val-U19G | Fwd | AAGCTTAATACGACTCACTATAGTTTCCGTAGTGTAGTGGGTATCACGTTCGCCTCACA |
| | Rev | TGTTTCCGCCCGGTTTCGAACCGGGGACCTTTCGCGTGTGAGGCGAACGTGATACCCACT |
| Val-ΔASL | Fwd | AAGCTTAATACGACTCACTATAGTTTCCGTAGTGTAGTGGTtATCACGTAAGGtCCCC |
| | Rev | TGTTTCCGCCCGGTTTCGAACCGGGGaCCTTACGTGATaACCACTACACTACGGAAACC |
| Val-ΔT | Fwd | AAGCTTAATACGACTCACTATAGTTTCCGTAGTGTAGTGGTTATCACGTTCGCCTCACA |
| | Rev | TGTTTCCGCGGaCCTTTCGCGTGTGAGGCGAACGTGATaACCACTACACTACGGAAACC |
| Val-ΔAcceptor | Fwd | AAGCTTAATACGACTCACTATAGCGAAAGGTCCCCGGTTCGAAACCG |
| | Rev | GCGAACGTGATAACCACTACACTACGCCCGGTTTCGAACCGGGGACCTTTCGC |
| Val-+CCA | Fwd | AAGCTTAATACGACTCACTATAGTTTCCGTAGTGTAGTGGTtATCACGTTCGCCTCACACG |
| | Rev | TGGTGTTTCCGCCCGGTTTCGAACCGGGGaCCTTTCGCGTGTGAGGCGAACGTGATAACCAC |
| Glu-TCC | Fwd | AAGCTTAATACGACTCACTATAGTCCCACATGGTCTAGCGGTtAGGATTCCTGGTTTTCA |
| | Rev | TTCCCACACCGGGAGTCGAACCCGGGCCGCCTGGGTGAAAACCAGGAATCCTaACCGCTA |
| Leu-CAA | Fwd | AAGCTTAATACGACTCACTATAGTCAGGATGGCCGAGTGGTCTAAGGCGCCAGACTCAAG |
| | Rev | TGTCAGAAGTGGGATTCGAACCCACGCCTCCATTGGAGACCAGAACTTGAGTCTGGCGCC |
| Gly-GCC | Fwd | AAGCTTAATACGACTCACTATAGCATGGGTGGTTCAGTGGTAGAATTCTCGCCTGCCAC |
| | Rev | TGCATGGGCCGGGAATCGAACCCGGGCCTCCCGCGTGGCAGGCGAGAATTCTACCACTGA |
| Asp-GTC | Fwd | AAGCTTAATACGACTCACTATAGCCTCGTTAGTATAGTGGTGAGTATCCCCGCCTGTCAC |
| | Rev | CTCCCCGTCGGGGAATCGAACCCCGGTCTCCCGCGTGACAGGCGGGGATACTCACCACTA |
| Pro-CGG | Fwd | AAGCTTAATACGACTCACTATAGGCTCGTTGGTCTAGGGGTATGATTCTCGCTTCGGGT |
| | Rev | GGGCTCGTCCGGGATTTGAACCCGGGACCTCTCACACCCGAAGCGAGAATCATACCCCTAG |
| Pro-+CCA | Fwd | AAGCTTAATACGACTCACTATAGGCTCGTTGGTCTAGGGGTATGATTCTCGCTTCGGGTGT |
| | Rev | TGGGGGCTCGTCCGGGATTTGAACCCGGGACCTCTCACACCCGAAGCGAGAATCATACCCC |
| Lys-TTT | Fwd | AAGCTTAATACGACTCACTATAGCCCGGATAGCTCAGTCGGTAGAGCATCAGACTTTTA |
| | Rev | CGCCCGAACAGGGACTTGAACCCTGGaCCCTCAGATTAAAAGTCTGATGCTCTACCGACT |
| Cys-GCA | Fwd | AAGCTTAATACGACTCACTATAGGGGGTATAGCTCAGTGGTAGAGCATTTGACTGCAGAT |
| | Rev | AGGGGGCACCCGGATTTGAACCGGGGACCTCTTGATCTGCAGTCAAATGCTCTACCACTG |
| Lys-CTT | Fwd | AAGCTTAATACGACTCACTATAGCCCGGCTAGCTCAGTCGGTAGAGCATGAGACTCTTAA |
| | Rev | CGCCCAACGTGGGGCTCGAACCCACGACCCTGAGATTAAGAGTCTCATGCTCTACCGACT |
| Tyr-GTA | Fwd | AAGCTTAATACGACTCACTATAGCCTTCGATAGCTCAGTTGGTAGAGCGGAGGACTGTAG |
| | Rev | TCCTTCGAGCCGGAATCGAACCAGCGACCTAAGGATCTACAGTCCTCCGCTCTACCAACT |
| Arg-ACG | Fwd | AAGCTTAATACGACTCACTATAGGGCCAGTGGCGCAATGGATAACGCGTCTGACTACGGAT |
| | Rev | CGAGCCAGCCAGGAGTCGAACCTGGAATCTTCTGATCCGTAGTCAGACGCGTTATCCAT |

**Supplementary Table 1.** DNA templates for IVT. tRNA sequences were obtained from gtRNA database. T7 RNA polymerase promoter is underlined.

| Sequence | Adduct mass |
|---|---|
| U | 324.04 |
| AU | 653.09 |
| GU | 669.08 |
| UU | 707.97 |
| UC | 629.08 |
| GUU/UGU | 1053.02 |
| GGU/GUG | 1014.13 |
| UUC | 1011.00 |
| UUU | 1089.89 |
| UCC/CCU/CUC | 932.10 |
| UCG/GUC | 972.11 |
| UAG | 996.12 |
| AUC | 958.13 |
| UAU | 1037.03 |
| GUAG | 1343.18 |
| UUAU | 1420.96 |
| ACGU | 1301.16 |
| UAUC | 1342.07 |

**Supplementary Table 2.** Neutral mass of potential RNA oligo fragments crosslinked to hDUS2 peptides after RNase A and T1 treatment and trypsin digestion.

| tRNA | D loop | | | | | | | | | | | | | Activity at 1 µM | | | Activity at 10 µM | | |
|---|---|---|---|---|---|---|---|---|---|---|---|---|---|---|---|---|---|---|---|
| | | | | | | | | | | | | | | Rep1 | Rep2 | Mean | Rep1 | Rep2 | Mean |
| Val-CAC | U | A | G | U | – | **G** | **G** | **U** | U | A | U | C | A | 0.46 | 0.58 | 0.52 | 0.51 | 0.54 | 0.53 |
| Glu-TTC | U | A | G | C | – | **G** | **G** | **U** | U | A | G | G | A | 0.43 | 0.57 | 0.50 | 0.54 | 0.51 | 0.53 |
| Gly-GCC | C | A | G | U | – | **G** | **G** | **U** | – | A | G | A | A | 0.19 | 0.41 | 0.30 | 0.49 | 0.44 | 0.47 |
| Cys-GCA | C | A | G | U | – | **G** | **G** | **U** | – | A | G | A | G | 0.06 | 0.44 | 0.25 | 0.43 | 0.41 | 0.42 |
| Lys-CTT | C | A | G | U | C | **G** | **G** | **U** | – | A | G | – | – | 0.13 | 0.22 | 0.17 | 0.37 | 0.32 | 0.35 |
| Met-CAT | C | A | G | U | – | **G** | **G** | G | C | A | G | C | G | 0.03 | 0.17 | 0.10 | 0.25 | 0.22 | 0.24 |
| Asp-GTC | U | A | G | U | – | **G** | **G** | **U** | G | A | G | U | A | 0.00 | 0.11 | 0.06 | 0.09 | 0.27 | 0.18 |
| Tyr-GTA | C | A | G | U | U | **G** | **G** | **U** | – | A | G | A | G | 0.02 | 0.25 | 0.14 | 0.17 | 0.15 | 0.16 |
| Pro-CGG | U | A | G | G | – | **G** | **G** | **U** | – | A | U | G | A | 0.02 | 0.05 | 0.04 | 0.10 | 0.17 | 0.14 |
| Arg-ACG | C | A | A | U | – | **G** | **G** | A | U | A | A | C | G | 0.01 | 0.02 | 0.015 | 0.004 | 0.008 | 0.006 |
| Phe-GAA | C | A | G | U | U | **G** | **G** | G | – | A | G | A | G | 0.00 | 0.00 | 0.00 | 0.007 | 0.00 | 0.004 |

**Supplementary Table 3.** D loop sequence and crosslinking efficiency for BrUrd-modified tRNA assayed in Figure 3c. tRNA sequence is from GtRNAdb (http://gtrnadb.ucsc.edu/). G18 and G19 are in bold and U20 is labeled in red. The crosslinking efficiency is from the crosslinking assay with either 1 µM or 10 µM BrUrd-modified tRNA and quantified based on the densitometry of anti-hDUS2 WB.

| Oligo | Sequence | Full mass | Charge (z) | Calculated m/z | Observed m/z |
|---|---|---|---|---|---|
| 1 | GGGGUGUAGUGGUUAUCACCCC | 7046.94 | 3 | 2347.98 | 2347.97 |
| 2 | GGGGUGUAGUGG-BrU-UAUCACCCC | 7124.85 | 3 | 2373.95 | 2373.96 |
| 3 | GGGGUGUAGUGGU-BrU-AUCACCCC | 7124.85 | 3 | 2373.95 | 2373.92 |
| 4 | GGGGUGUAGUGG-BrU-BrU-AUCACCCC | 7202.76 | 3 | 2399.92 | 2399.91 |

**Supplementary Table 4.** Synthetic oligonucleotides used in this work.

| tRNA | D loop | | | | | | | | | | | | [D]/[D+U] |
|---|---|---|---|---|---|---|---|---|---|---|---|---|---|
| Glu-TTC | U | A | G | C | — | **G** | **G** | **U** | U | A | G | G | A | 0.58 ± 0.021 |
| Leu-CAA | G | A | G | U | — | **G** | **G** | **U** | C | U | A | A | G | 0.49 ± 0.003 |
| Gly-GCC | C | A | G | U | — | **G** | **G** | **U** | — | A | G | A | A | 0.47 ± 0.003 |
| Val-CAC | U | A | G | U | — | **G** | **G** | **U** | U | A | U | C | A | 0.40 ± 0.007 |
| Val-G18A | U | A | G | U | — | A | **G** | **U** | U | A | U | C | A | 0.35 ± 0.006 |
| Asp-GTC | U | A | G | U | — | **G** | **G** | **U** | G | A | G | U | A | 0.32 ± 0.015 |
| Pro-CGG | U | A | G | G | — | **G** | **G** | **U** | — | A | U | G | A | 0.30 ± 0.005 |
| Lys-TTT | C | A | G | U | C | **G** | **G** | **U** | — | A | G | — | — | 0.29 ± 0.027 |
| Val-U20G | U | A | G | U | — | **G** | **G** | G | **U** | A | U | C | A | 0.24 ± 0.005 |
| Cys-GCA | C | A | G | U | — | **G** | **G** | **U** | — | A | G | A | G | 0.20 ± 0.009 |
| Lys-CTT | C | A | G | U | C | **G** | **G** | **U** | — | A | G | — | — | 0.18 ± 0.009 |
| Arg-A20G | C | A | A | U | — | **G** | **G** | G | **U** | A | A | C | G | 0.15 ± 0.006 |
| Tyr-GTA | C | A | G | U | U | **G** | **G** | **U** | — | A | G | A | G | 0.08 ± 0.005 |
| Arg-ACG | C | A | A | U | — | **G** | **G** | A | U | A | A | C | G | 0.00 ± 0.000 |
| Val-U20C | U | A | G | U | — | **G** | **G** | C | U | A | U | C | A | 0.00 ± 0.000 |

**Supplementary Table 5.** D loop sequence and dihydrouridine modification stoichiometry for tRNAs and their mutants assayed in Figure 4c. G18 and G19 are labeled in bold and D site is labeled in bold and red. Modification stoichiometry was calculated by oligonucleotide LC-MS quantification comparing the D-containing fragment against the corresponding unmodified U-containing fragment. Values represent mean (n=3). Modification sites are labeled in red.

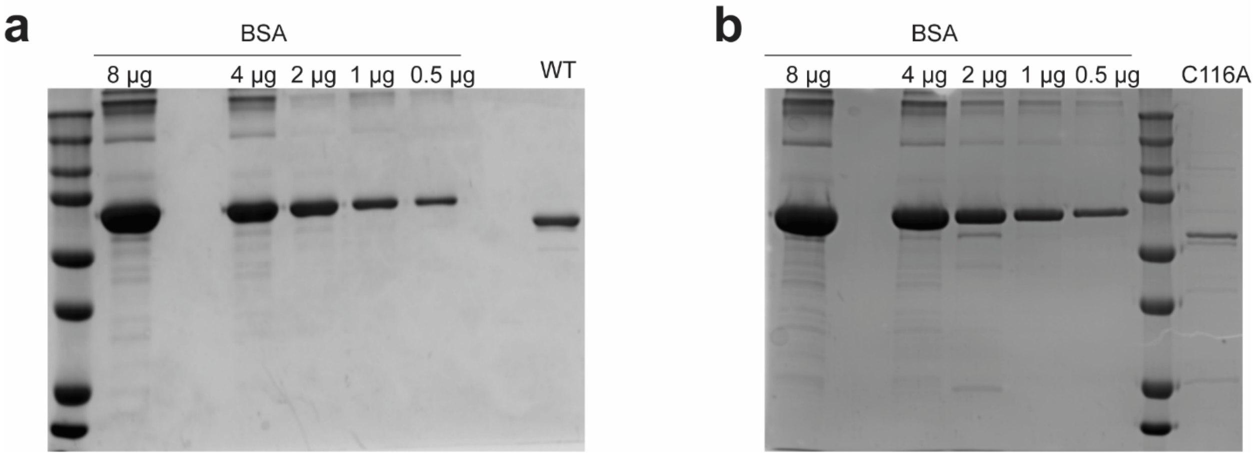


**Supplementary Figure 1.** SDS-PAGE analysis of recombinant purified $His_6$-hDUS2 **(a)** and $His_6$-hDUS2 (C116A) **(b)** with BSA as the standard for concentration quantification.

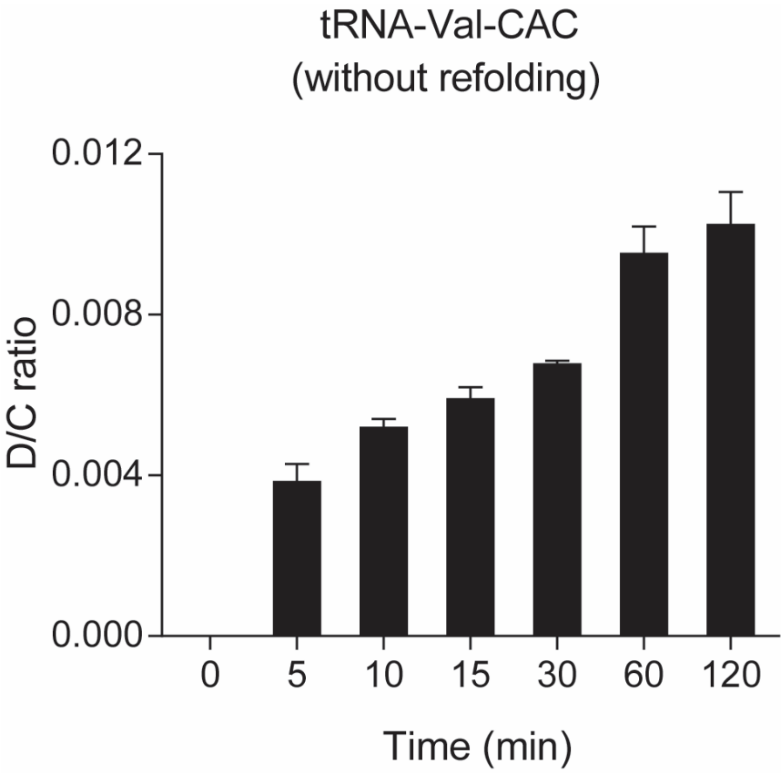


**Supplementary Figure 2.** LC-QQQ-MS quantitation of D formation on IVT tRNA-Val-CAC (prepared without refolding) by WT hDUS2. Recombinant hDUS2 (WT) was incubated with tRNA-Val-CAC in reaction buffer. D formation was quantified by nucleoside LC-QQQ-MS. Three independent biological replicates were analyzed. Values represent mean ± s.d. (n=3).

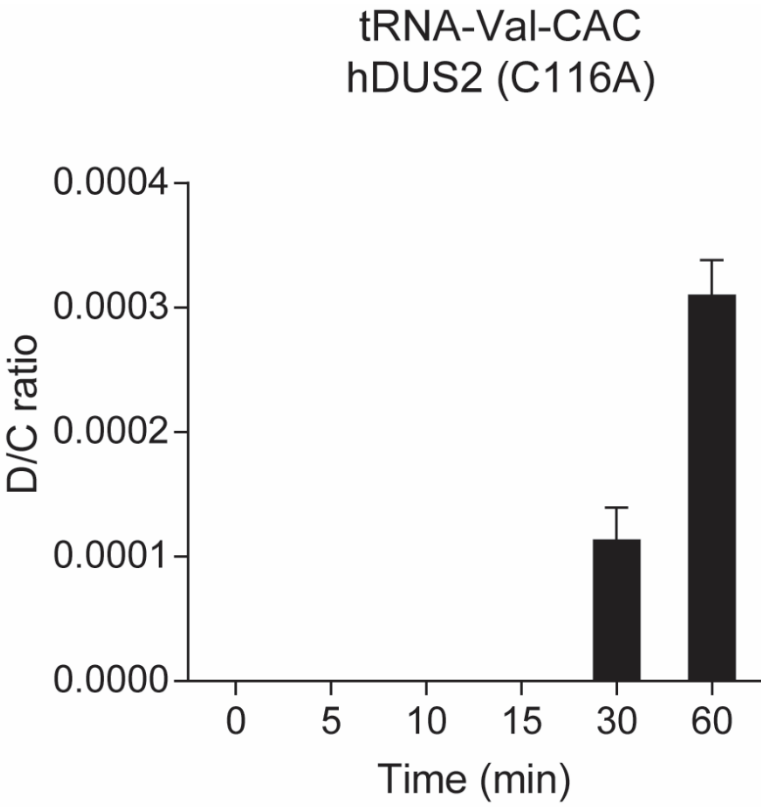


**Supplementary Figure 3.** LC-QQQ-MS quantitation of D formation on tRNA-Val-CAC. D formation on IVT tRNA-Val-CAC by hDUS2-C116A. The experiment was performed as in Supplementary Figure 2. Three independent biological replicates were analyzed. Values represent mean ± s.d. (n=3).

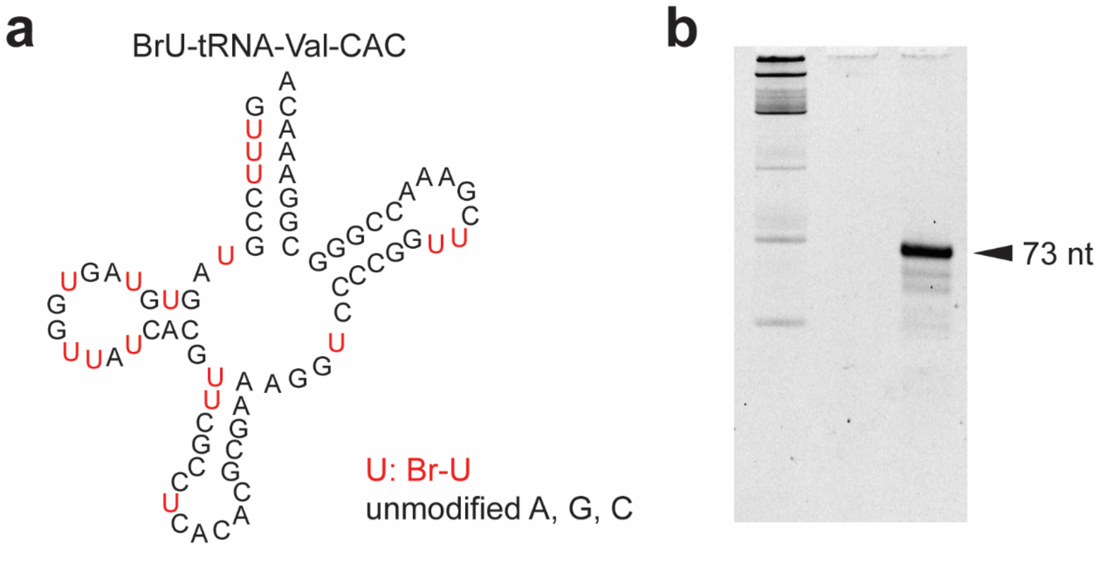


**Supplementary Figure 4.** BrUrd-modified tRNA-Val-CAC. **(a)** Structure of *in vitro* transcribed BrUrd-modified tRNA-Val-CAC. Residues in red represent BrUrd modification. **(b)** Purified IVT BrUrd-modified tRNA-Val-CAC.

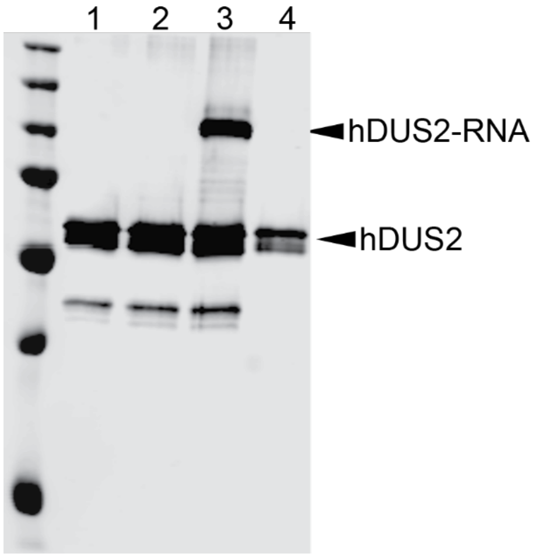


**Supplementary Figure 5.** Crosslinking between BrUrd-modified tRNA-Val-CAC and $His_6$-hDUS2. Full Western blot data for Figure 2b in the main text. Crosslinking was performed by incubating 200 nM of recombinant hDUS2 with 1 µM BrUrd-modified tRNA-Val-CAC in 20 mM Tris-HCl pH 8.0, 100 mM NaCl, 5 mM $MgCl_2$, 2 mM DTT, 10 mM NADPH in a total volume of 20 µl at 37 °C for 1 hour. RNase control samples were treated with 2 µl of RNase A/T1 mix at 37 °C for 30 min. Reactions were analyzed by anti-hDUS2 Western blot. 1, hDUS2 only; 2, hDUS2 with unmodified tRNA-Val-CAC; 3, hDUS2 with BrUrd-modified tRNA-Val-CAC; 4, RNase treatment of 3.

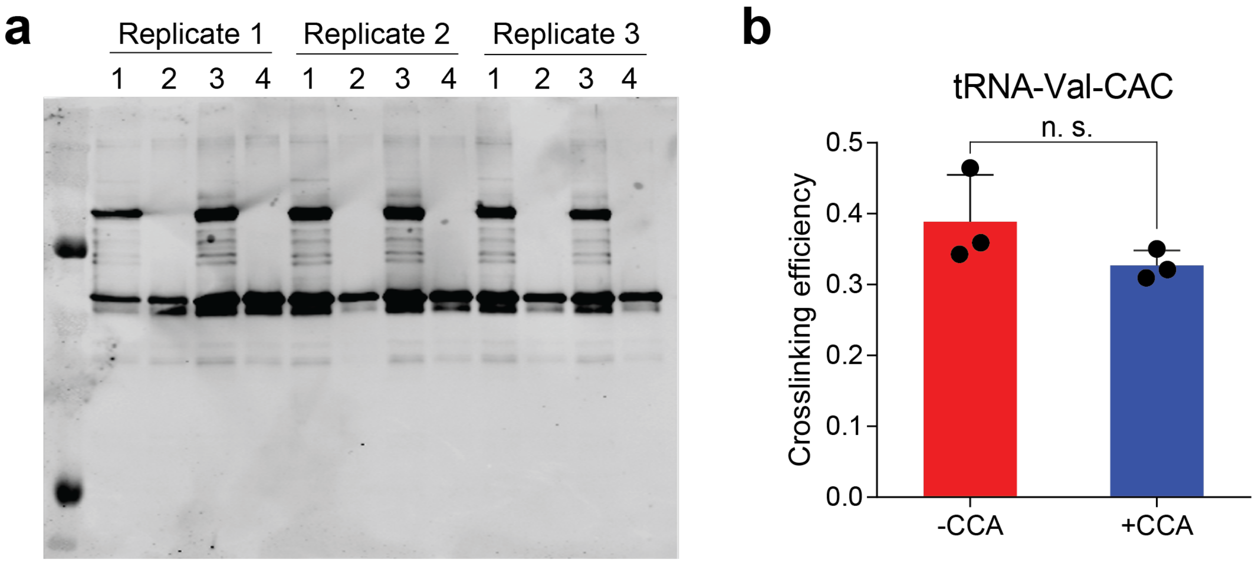


**Supplementary Figure 6.** Crosslinking of hDUS2 with BrUrd-modified tRNA-Val-CAC (lacking final CCA) or BrU-modified end-matured tRNA-Val-CAC (+CCA). **(a)** Crosslinking was performed by incubating 200 nM recombinant hDUS2 with 10 µM BrUrd-modified tRNA in 20 mM Tris-HCl pH 8.0, 100 mM NaCl, 5 mM $MgCl_2$, 2 mM DTT, 10 mM NADPH at 37 °C for 1 hour. RNase control samples were treated with 2 µl of RNase A/T1 mix at 37 °C for 30 min. Reactions were analyzed by anti-hDUS2 Western blot and three independent replicates were performed. 1, WT hDUS2 with BrU-modified tRNA-Val-CAC (-CCA); 2, RNase digestion of 1; 3, WT hDUS2 with full-length BrU-modified tRNA-Val-CAC (+CCA); 4, RNase digestion of 3. **(b)** Quantitation of crosslinking efficiency from **(a)**. Values represent mean ± s.d. A standard t-test was used to determine statistical significance; n.s. = not significant.

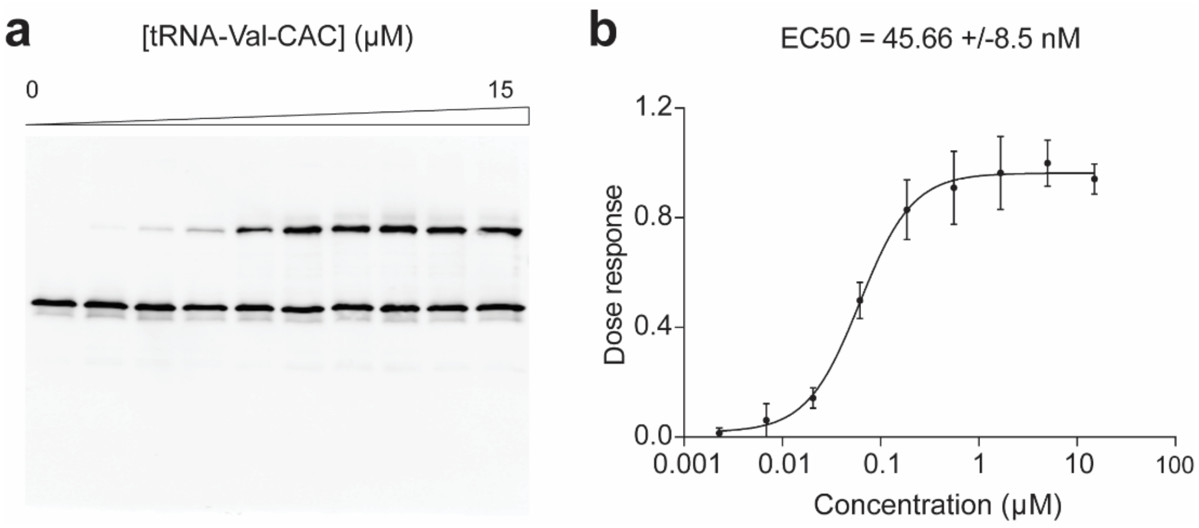


**Supplementary Figure 7.** Dose-dependent crosslinking between hDUS2 and BrUrd-modified tRNA-Val-CAC. **(a)** 200 nM hDUS2 was mixed with various concentrations of BrUrd-modified tRNA-Val-CAC (0-15 μM) in 20 mM Tris-HCl pH 8.0, 100 mM NaCl, 5 mM $MgCl_2$, 2 mM DTT, 10 mM NADPH in a total volume of 20μl at 37 °C for 1 hour. Anti-hDUS2 WB was used to detect product formation. Three independent replicates were performed. **(b)** Quantification of crosslinking in (a). $EC_{50}$ value was determined by fitting data to a sigmoidal dose-response curve. Values represent mean ± s.d (n=3).

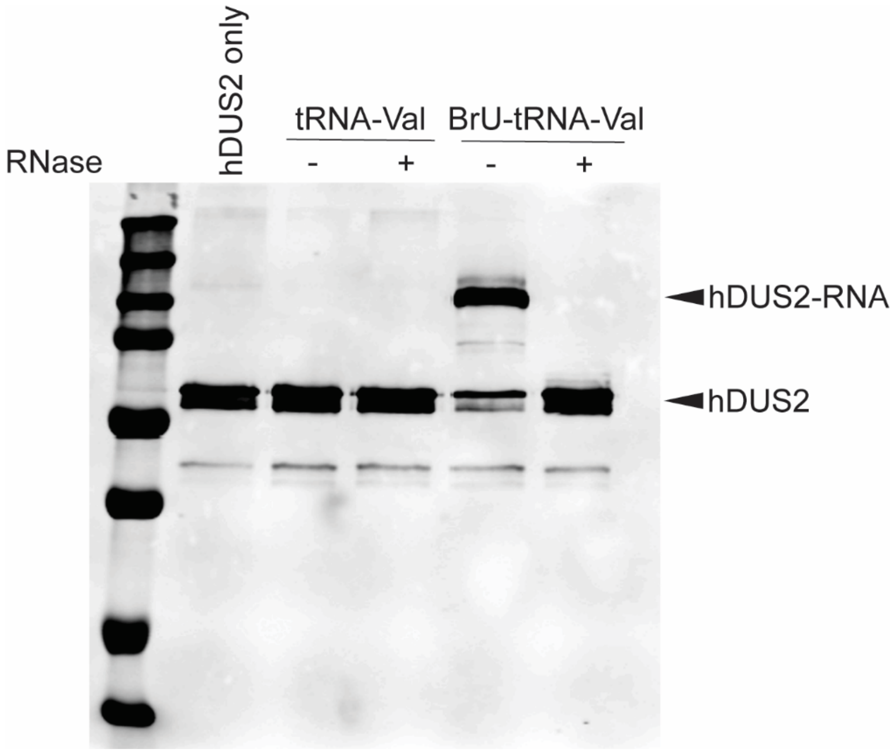


**Supplementary Figure 8.** Crosslinking between BrUrd-modified tRNA-Val-CAC and $His_6$-hDUS2 for proteomics analysis. Crosslinking was performed by incubating 2 µg of recombinant hDUS2 with 7.5 µg of BrUrd-modified tRNA-Val-CAC in 20 mM Tris-HCl pH 8.0, 100 mM NaCl, 5 mM $MgCl_2$, 2 mM DTT, 10 mM NADPH in a total volume of 50 µl at 37 °C for 1 hour. RNase control samples were treated with 2 µl of RNase A/T1 mix at 37 °C for 30 min. Reactions were analyzed by anti-hDUS2 Western blot.

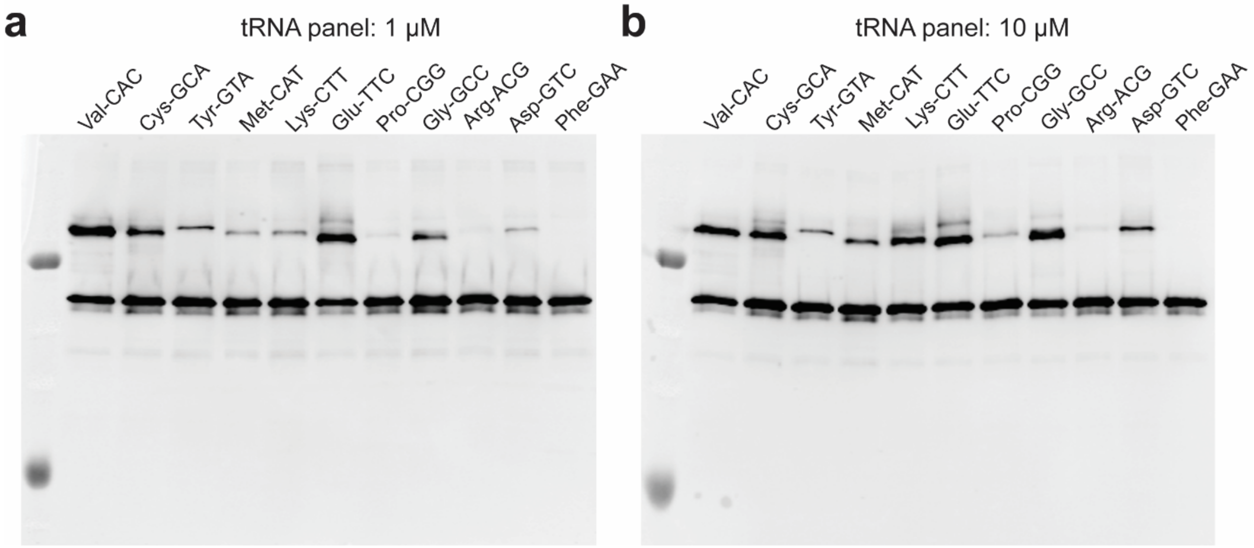


**Supplementary Figure 9.** Crosslinking assay between hDUS2 and BrU-modified tRNA panel. Full western blot data for Figure 3a and Figure 3c. 200 nM of protein was incubated with 1 µM **(a)** or 10 µM **(b)** of BrUrd-modified tRNA in 20 mM Tris-HCl pH 8.0, 100 mM NaCl, 5 mM $MgCl_2$, 2 mM DTT, 10 mM NADPH in a total volume of 20 µl for 1 hour at 37 °C. Reactions were analyzed by anti-hDUS2 WB. Two independent replicates were performed.

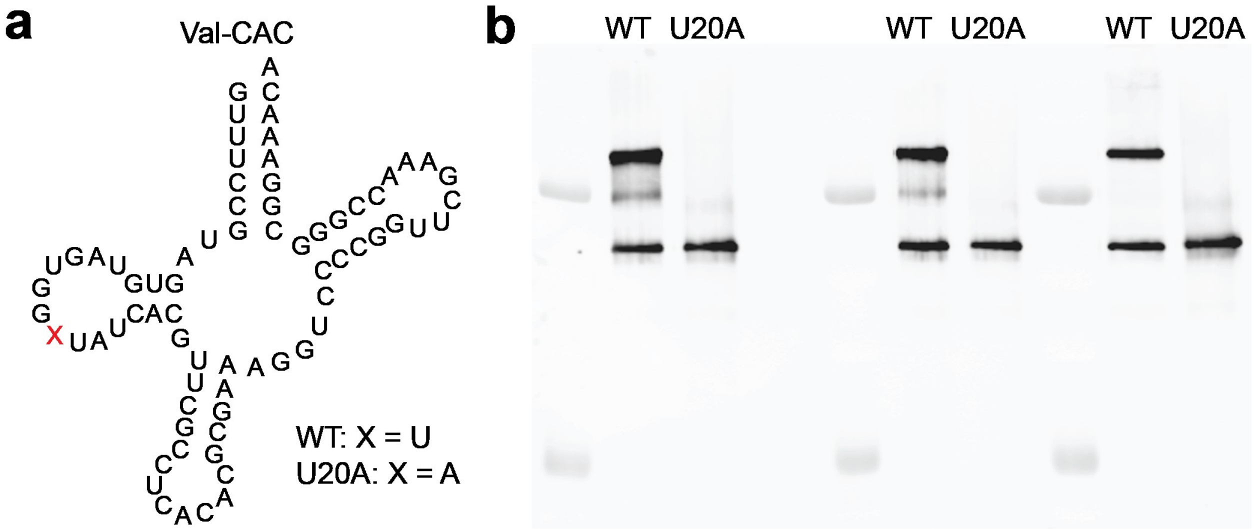


**Supplementary Figure 10.** Crosslinking assay between hDUS2 and BrUrd-modified tRNA-Val-CAC and U20A mutant. **(a)** Structure of tRNA-Val-CAC constructs**.** **(b)** Crosslinking result. Experiments were conducted as in Supplementary Fig. 6. 1 μM BrUrd-modified tRNA was used.

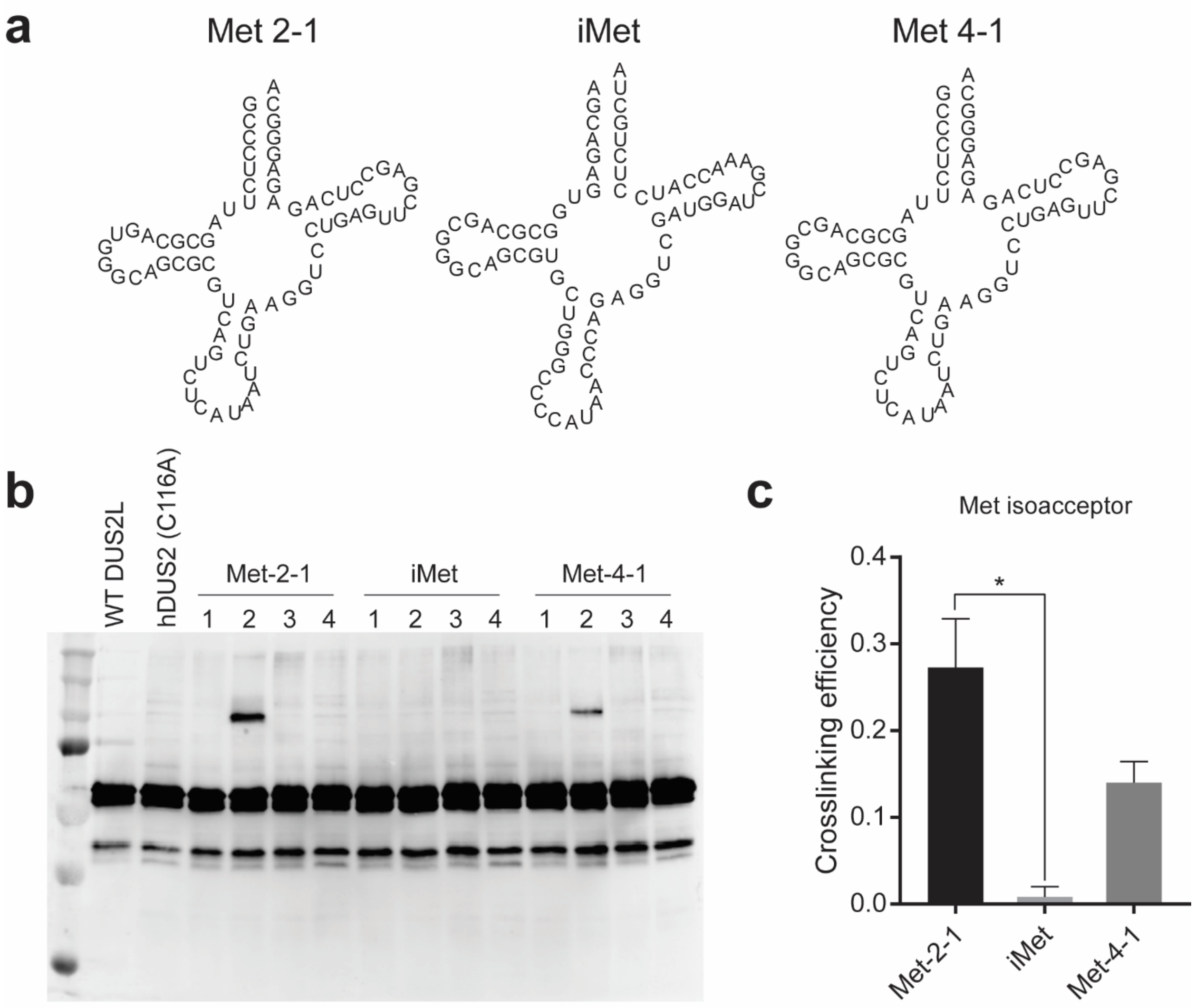


**Supplementary Figure 11.** Crosslinking assay between hDUS2 and BrUrd-modified tRNA-Met isoacceptors. **(a)** Architecture of tRNA-Met isoacceptors. **(b)** Crosslinking experiments were conducted as in Supplementary Fig. 6. 10 μM BrUrd-modified tRNA-Met was used. 1, WT hDUS2 with unmodified tRNA; 2, WT hDUS2 with BrUrd-modified tRNA; 3, RNase digestion of 2; 4, hDUS2-C116A with BrUrd-modified tRNA. Three independent replicates were performed. **(c)** Densitometry analysis of Western blot. Values represent mean ± s.d. (n=3). Crosslinking efficiency equals the intensity of crosslinked adduct over total protein intensity. A student's t-test was used to determine statistical significance between each tRNA isoacceptors. $p < 0.05$ is represented by *.

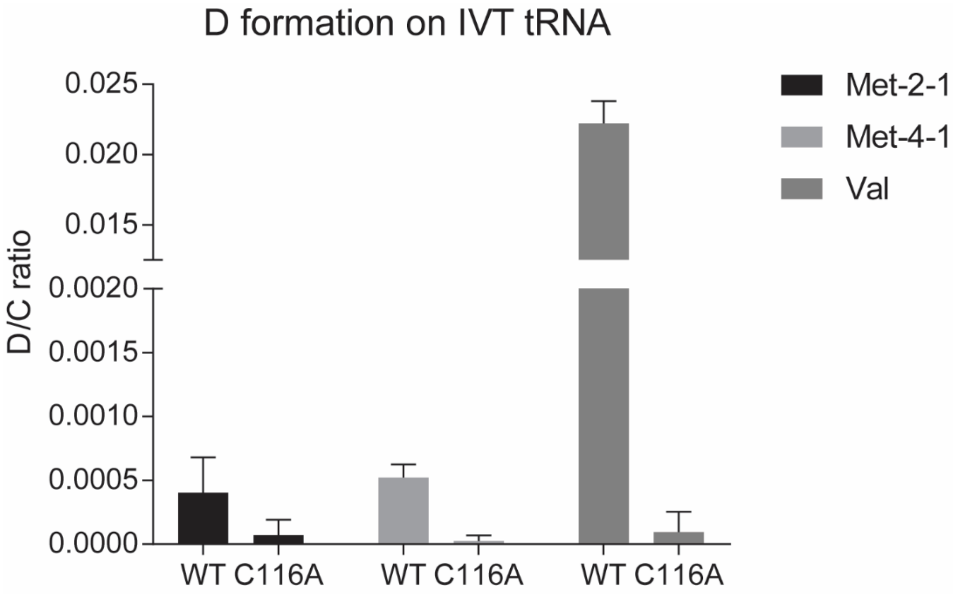


**Supplementary Figure 12.** LC-QQQ-MS quantitation of D formation on tRNA-Met isoacceptors. The experiment was conducted as in Supplementary Fig. 2. Values represent mean ± s.d. (n=3).

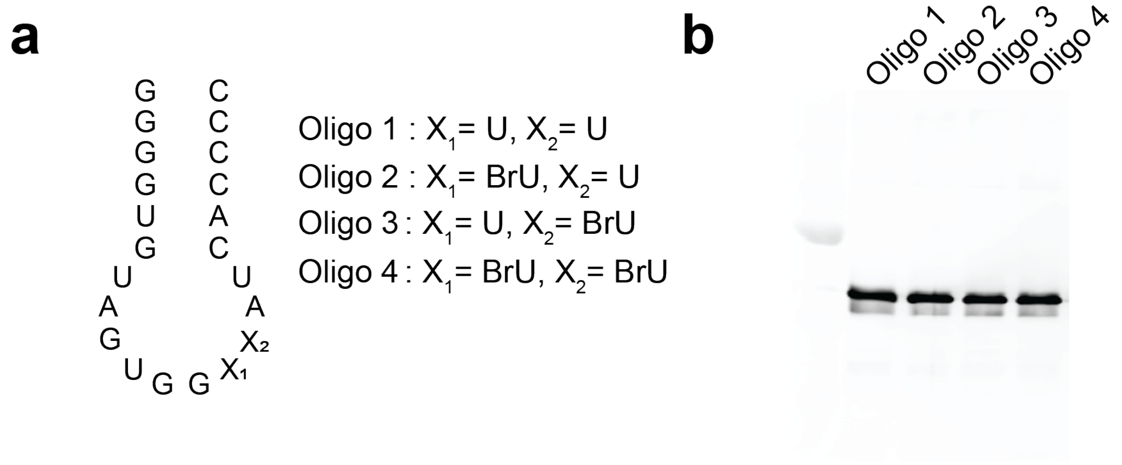


**Supplementary Figure 13.** Crosslinking assay between hDUS2 and synthetic BrU-containing oligos and unmodified oligo. **(a)** Sequence and structure of synthetic oligos with site-specific BrU modification. **(b)** Experiments were conducted as in Supplementary Fig. 6 using 100 μM oligo.

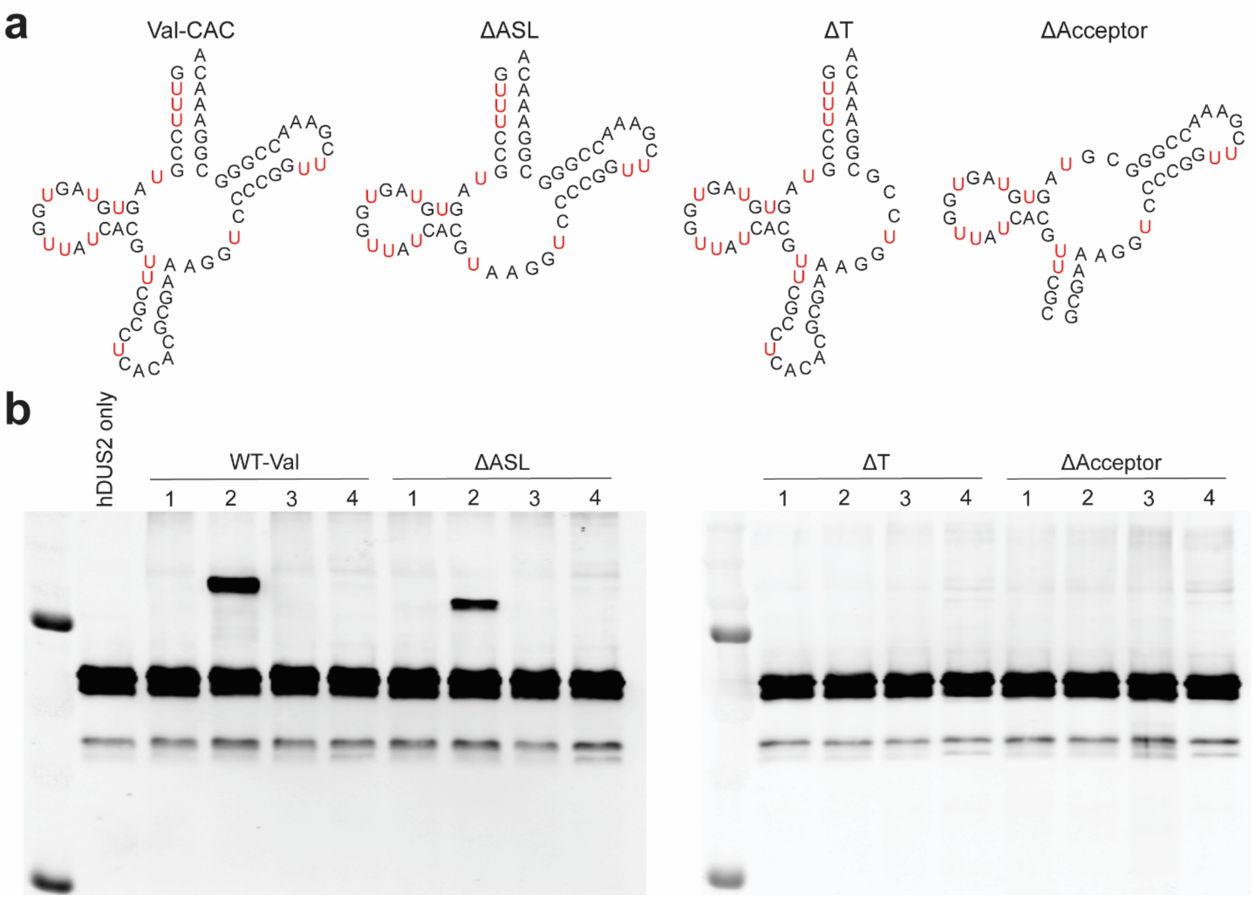


**Supplementary Figure 14.** Crosslinking assay between hDUS2 and truncated BrUrd-modified tRNA-Val-CAC. **(a)** Architecture of truncated tRNA-Val constructs. Residues in red represent BrU modification. **(b)** Full Western blot data for Figure 3e in the main text. Experiment was conducted as in Supplementary Fig. 5. 1, WT hDUS2 with unmodified tRNA; 2, WT hDUS2 with BrU-modified tRNA; 3, RNase digestion of 2; 4, hDUS2-C116A with BrU-modified tRNA.

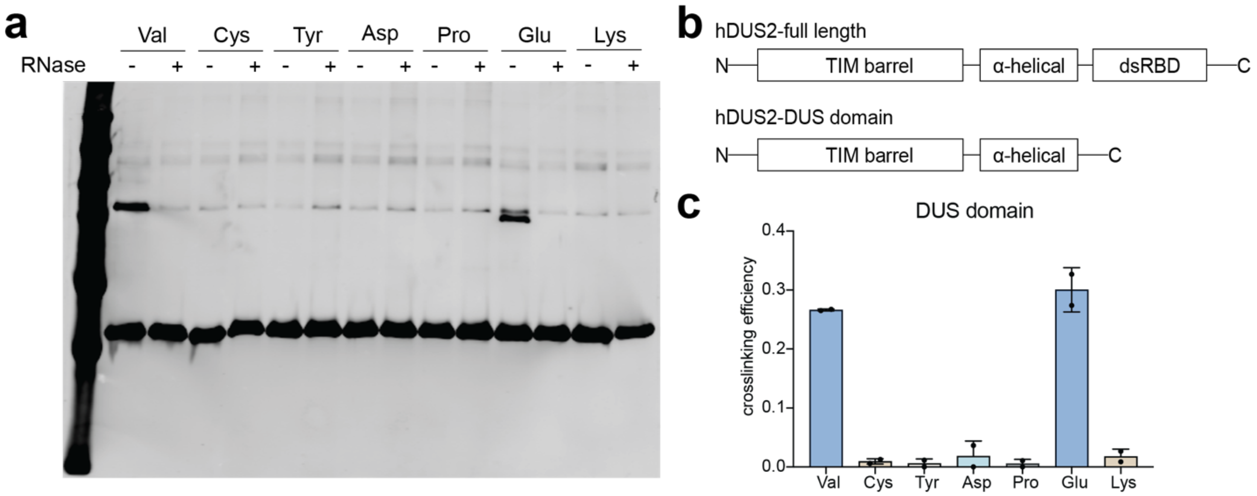


**Supplementary Figure 15.** Crosslinking assay between hDUS2-DUS domain and BrU-modified tRNA panel. **(a)** Experiment was conducted as in Supplementary Fig. 5 and reactions were analyzed by anti-His western blot. **(b)** Domain architecture of hDUS2 full-length and DUS domain. **(c)** Densitometry analysis of Western blot **(a)**. Values represent mean ± s.d. (n=2). Crosslinking efficiency equals intensity of crosslinked adduct over total protein intensity.

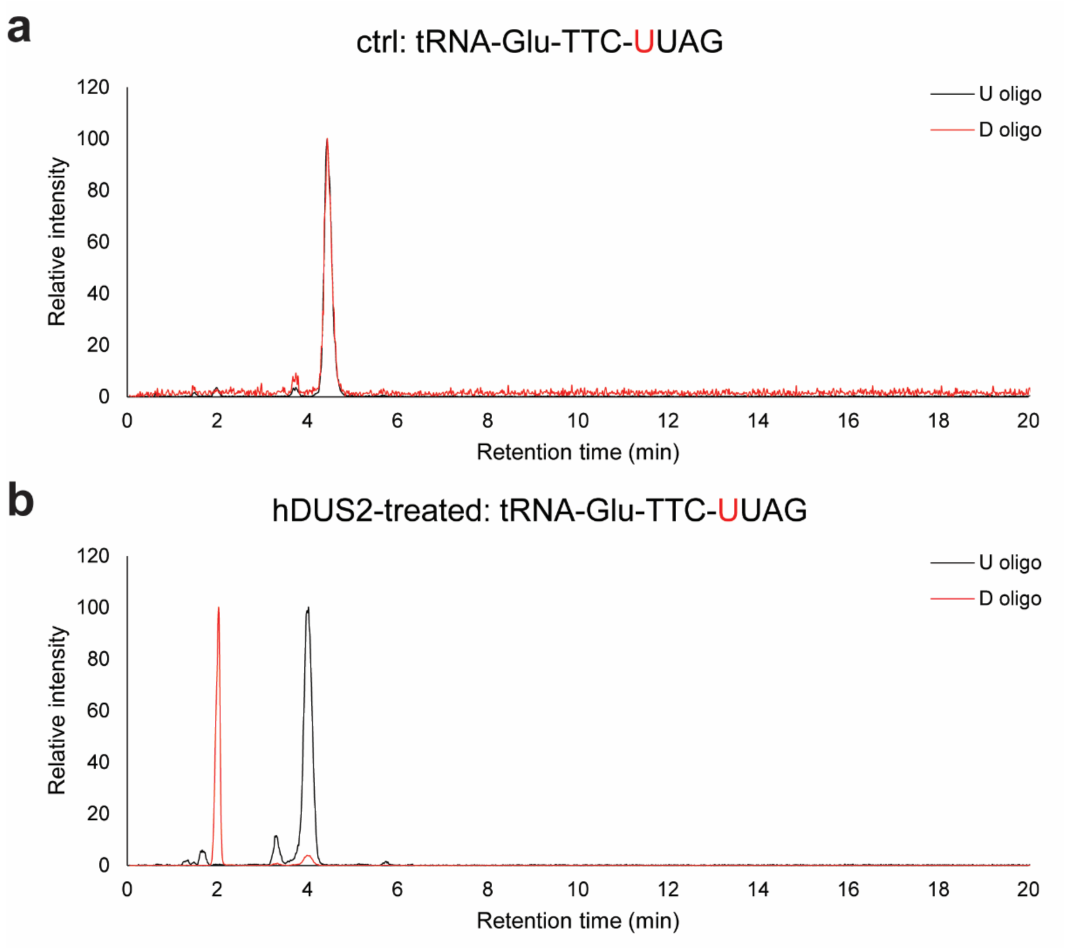


**Supplementary Figure 16.** Oligonucleotide LC-MS analysis of hDUS2-catalyzed dihydrouridine formation on tRNA-Glu-TTC. Extracted ion chromatogram of characteristic oligonucleotide fragment from RNase T1 digestion in control tRNA sample **(a)** and hDUS2-treated tRNA **(b)**. U-oligo: $[M\text{-}H]^-$ = 1303.174; D-oligo: $[M\text{-}H]^-$ = 1305.189.

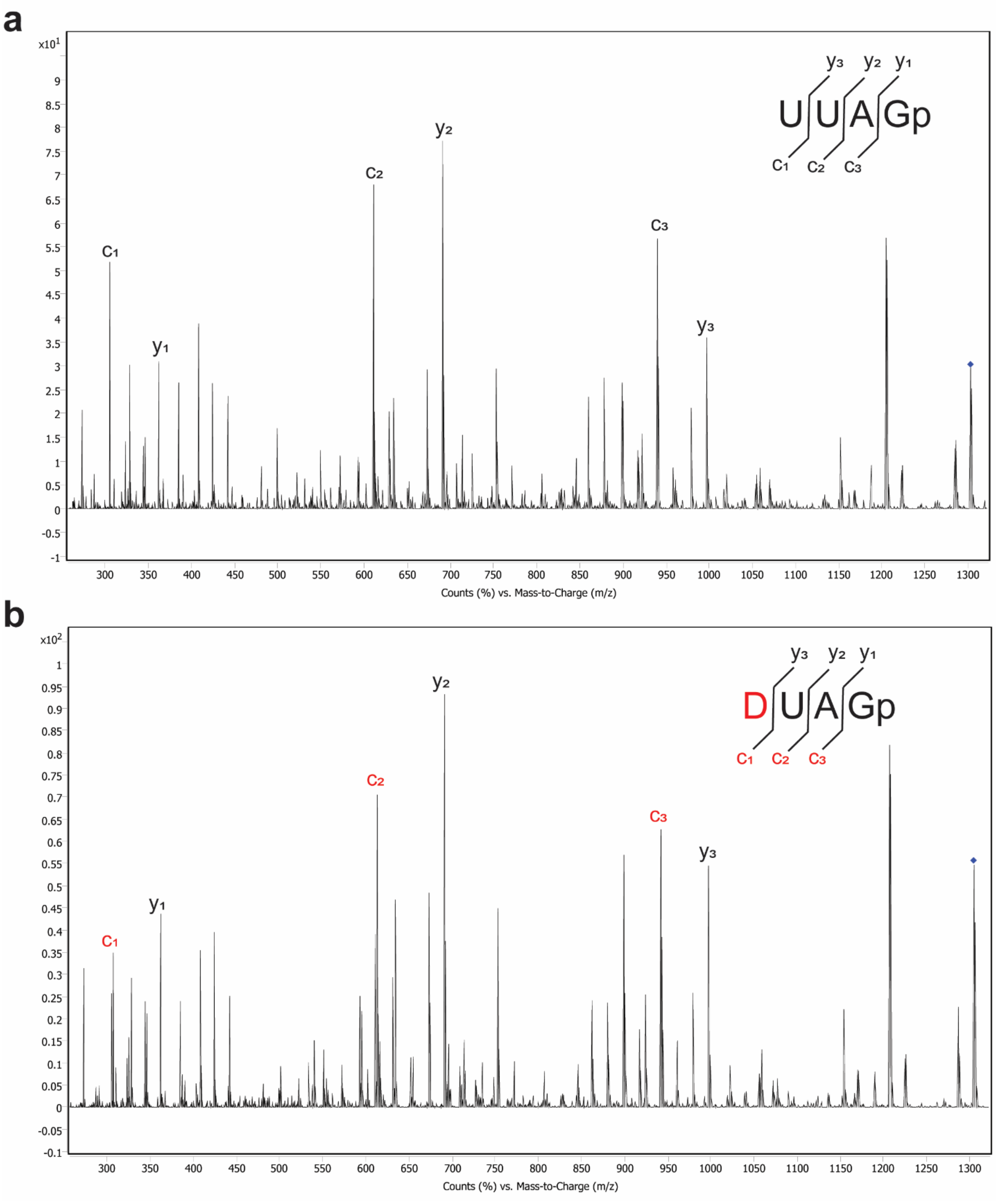


**Supplementary Figure 17.** MS/MS spectrum of UUAGp and DUAGp oligo fragments produced from RNase T1 digestion of control tRNA-Glu-TTC sample **(a)** and hDUS2-treated tRNA-Glu-TTC **(b)**. MS/MS fragmentation was performed by collision-induced dissociation. The U-containing precursor ion is $[M\text{-}H]^-$ = 1303.174. The D-containing precursor ion is $[M\text{-}H]^-$ = 1305.189.

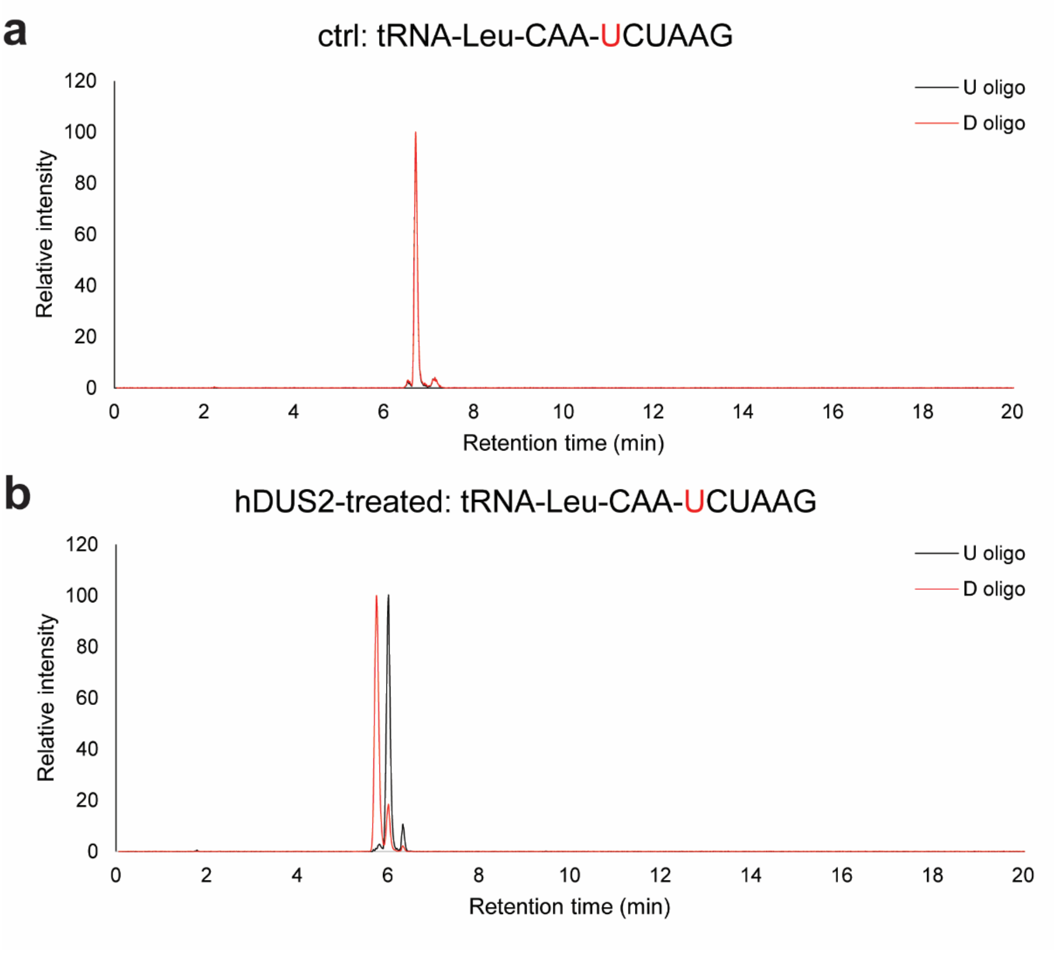


**Supplementary Figure 18.** Oligonucleotide LC-MS analysis of hDUS2-catalyzed dihydrouridine formation on tRNA-Leu-CAA. Extracted ion chromatography of characteristic oligonucleotide fragment from RNase T1 digestion in control tRNA sample **(a)** and hDUS2-treated tRNA **(b)**. U-oligo: $[M-2H]^{2-}$ = 968.1434; D-oligo: $[M-2H]^{2-}$ = 969.1509.

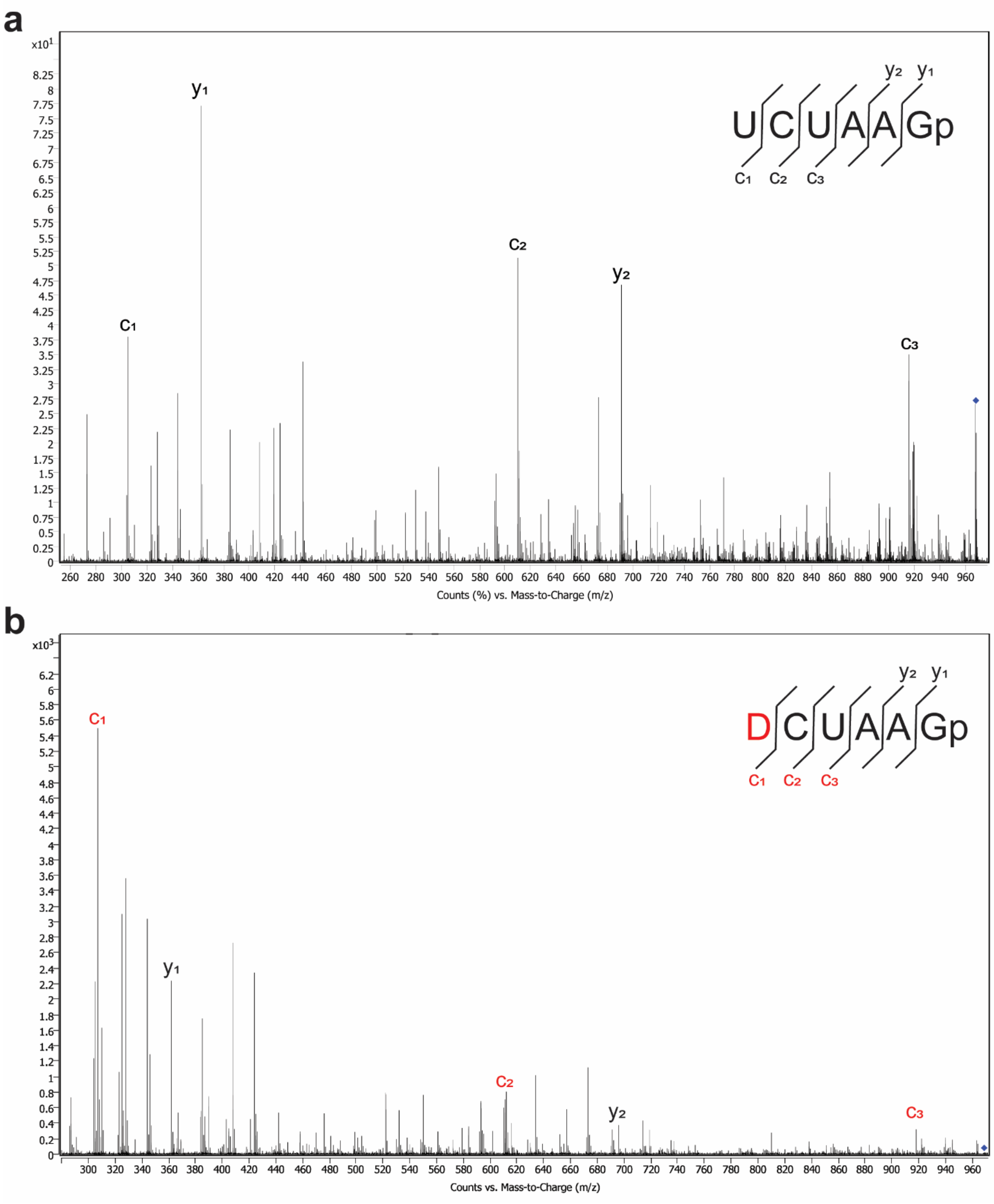


**Supplementary Figure 19.** MS/MS spectrum of UCUAAGp and DCUAAGp oligo fragments produced from RNase T1 digestion of control tRNA-Leu-CAA sample **(a)** and hDUS2-treated tRNA-Leu-CAA **(b)**. MS/MS fragmentation was performed by collision-induced dissociation. The U-containing precursor ion is $[M\text{-}2H]^{2-}$ = 968.1434. The D-containing precursor ion is $[M\text{-}2H]^{2-}$ = 969.1509.

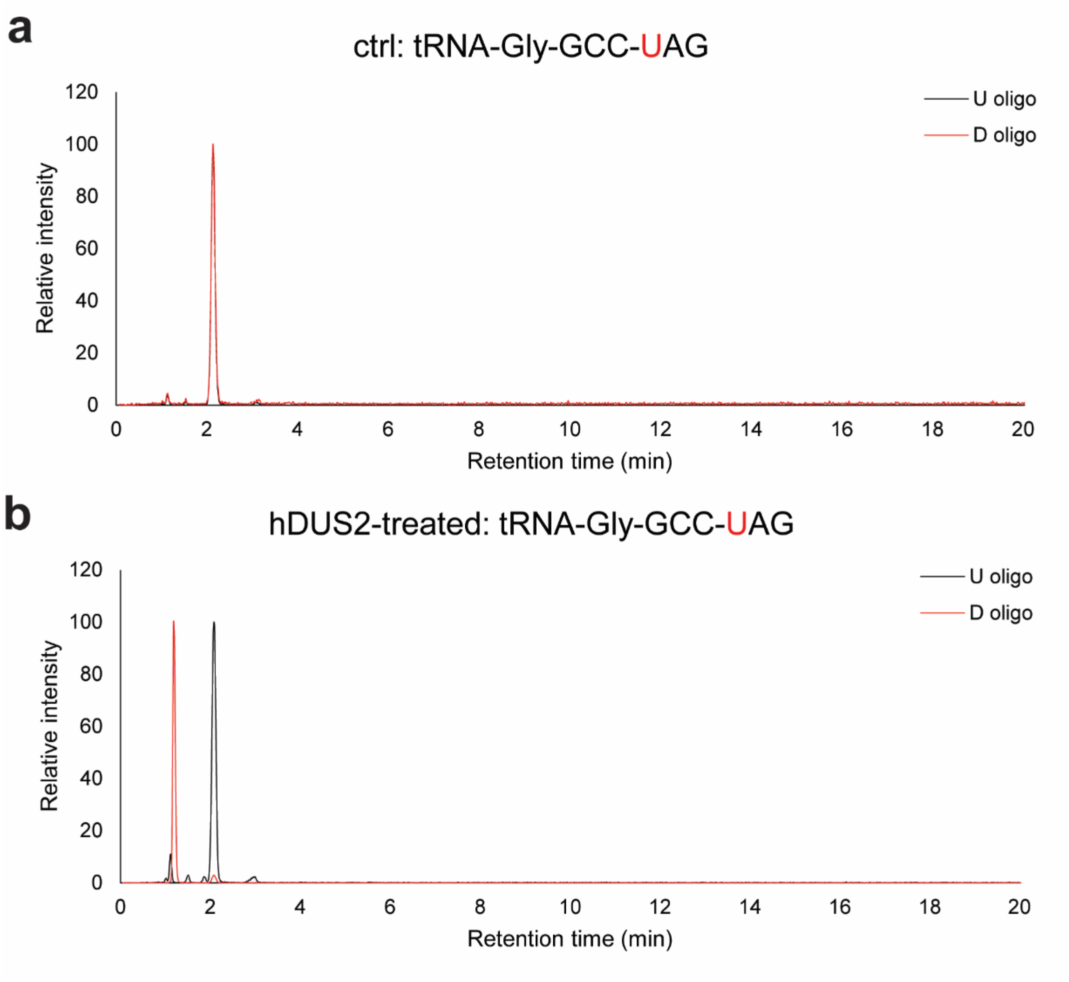


**Supplementary Figure 20.** Oligonucleotide LC-MS analysis of hDUS2-catalyzed dihydrouridine formation on tRNA-Gly-GCC. Extracted ion chromatography of characteristic oligonucleotide fragment from RNase T1 digestion in control tRNA sample **(a)** and hDUS2-treated tRNA **(b)**. U-oligo: $[M-H]^-$ = 997.1517; D-oligo: $[M-H]^-$ = 999.1597.

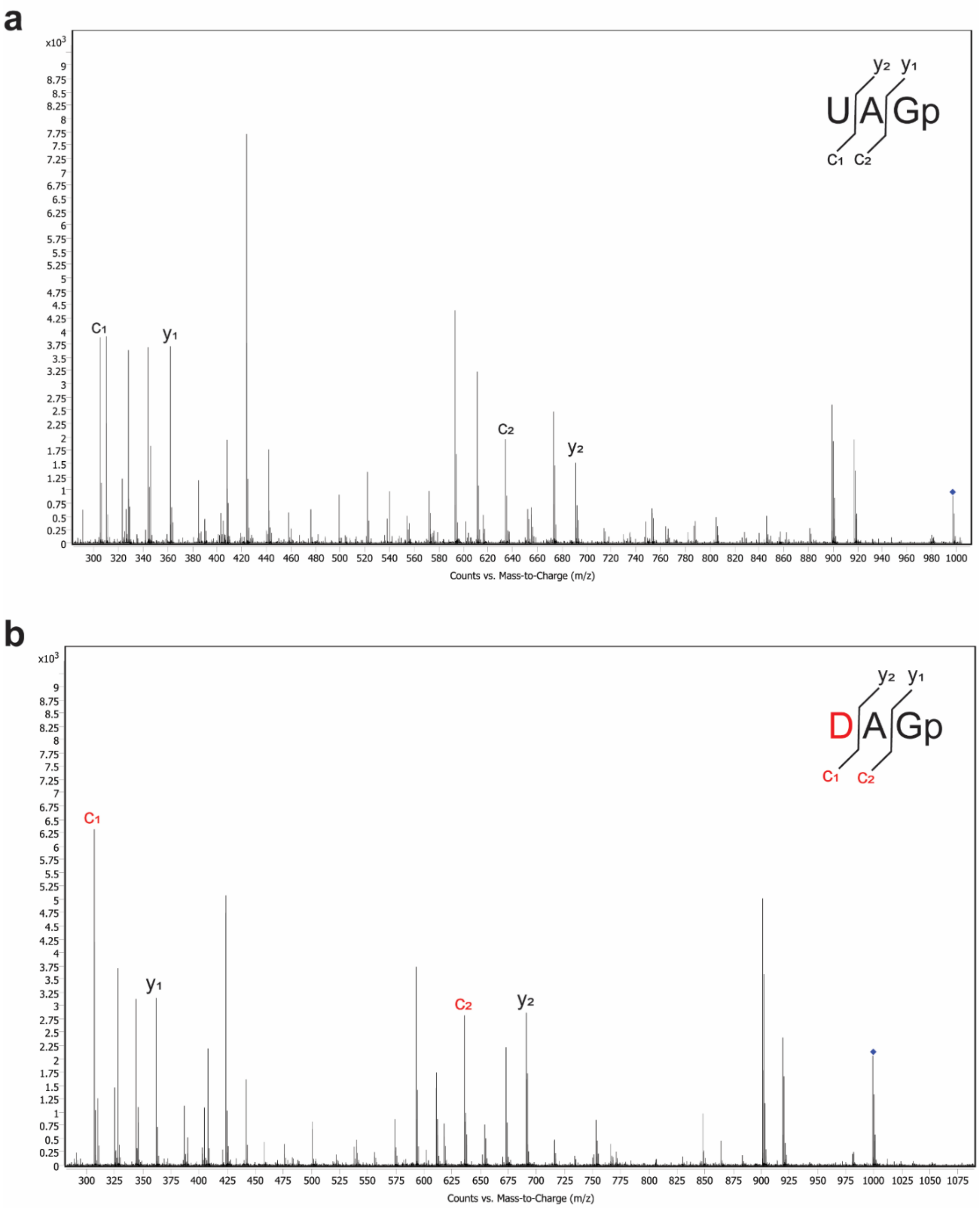


**Supplementary Figure 21.** MS/MS spectrum of UAGp and DAGp oligo fragments produced from RNase T1 digestion of control tRNA-Gly-GCC sample **(a)** and hDUS2-treated tRNA-Gly-GCC **(b)**. MS/MS fragmentation was performed by collision-induced dissociation. The U-containing precursor ion is [M-H]$^-$ = 997.1517. The D-containing precursor ion is [M-H]$^-$ = 999.1597.

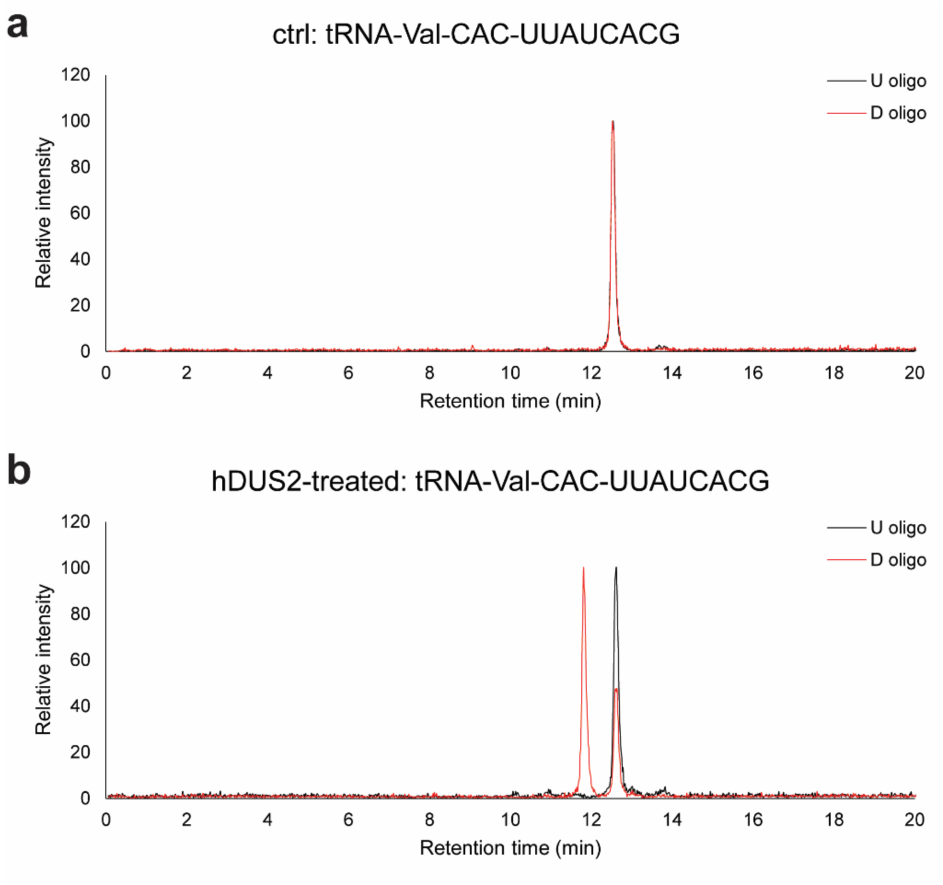


**Supplementary Figure 22.** Oligonucleotide LC-MS analysis of hDUS2-catalyzed dihydrouridine formation on tRNA-Val-CAC. Extracted ion chromatography of characteristic oligonucleotide fragment from RNase T1 digestion in control tRNA sample **(a)** and hDUS2-treated tRNA **(b)**. U-oligo: $[M-2H]^{2-}$ = 1273.669; D-oligo: $[M-2H]^{2-}$ = 1274.677.

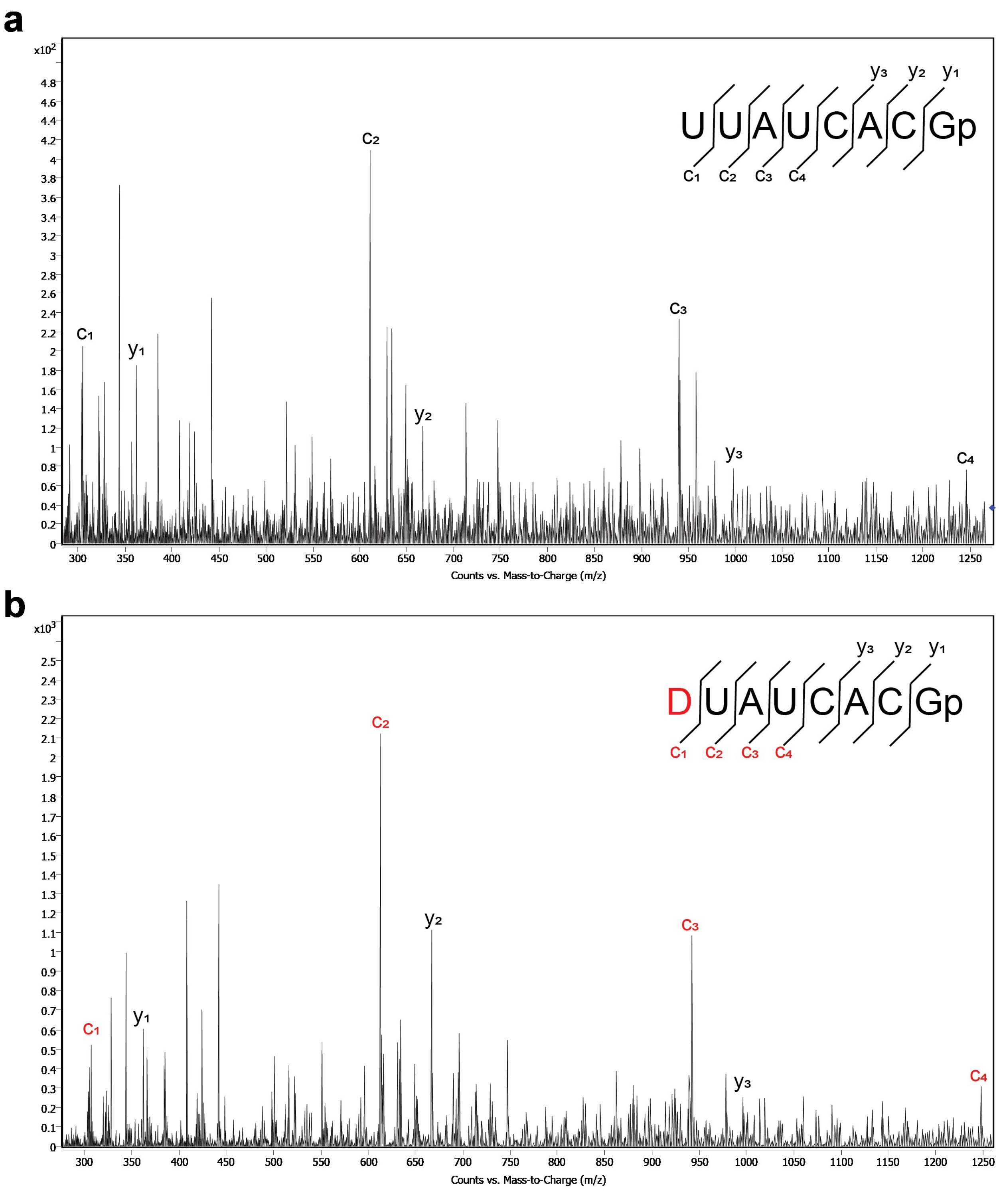


**Supplementary Figure 23.** MS/MS spectrum of UUAUCACGp and DUAUCACGp oligo fragments produced from RNase T1 digestion of control tRNA-Val-CAC sample **(a)** and hDUS2-treated tRNA-Val-CAC **(b)**. MS/MS fragmentation was performed by collision-induced dissociation. The U-containing precursor ion is $[M-2H]^{2-}$ = 1273.669. The D-containing precursor ion is $[M-2H]^{2-}$ = 1274.677.

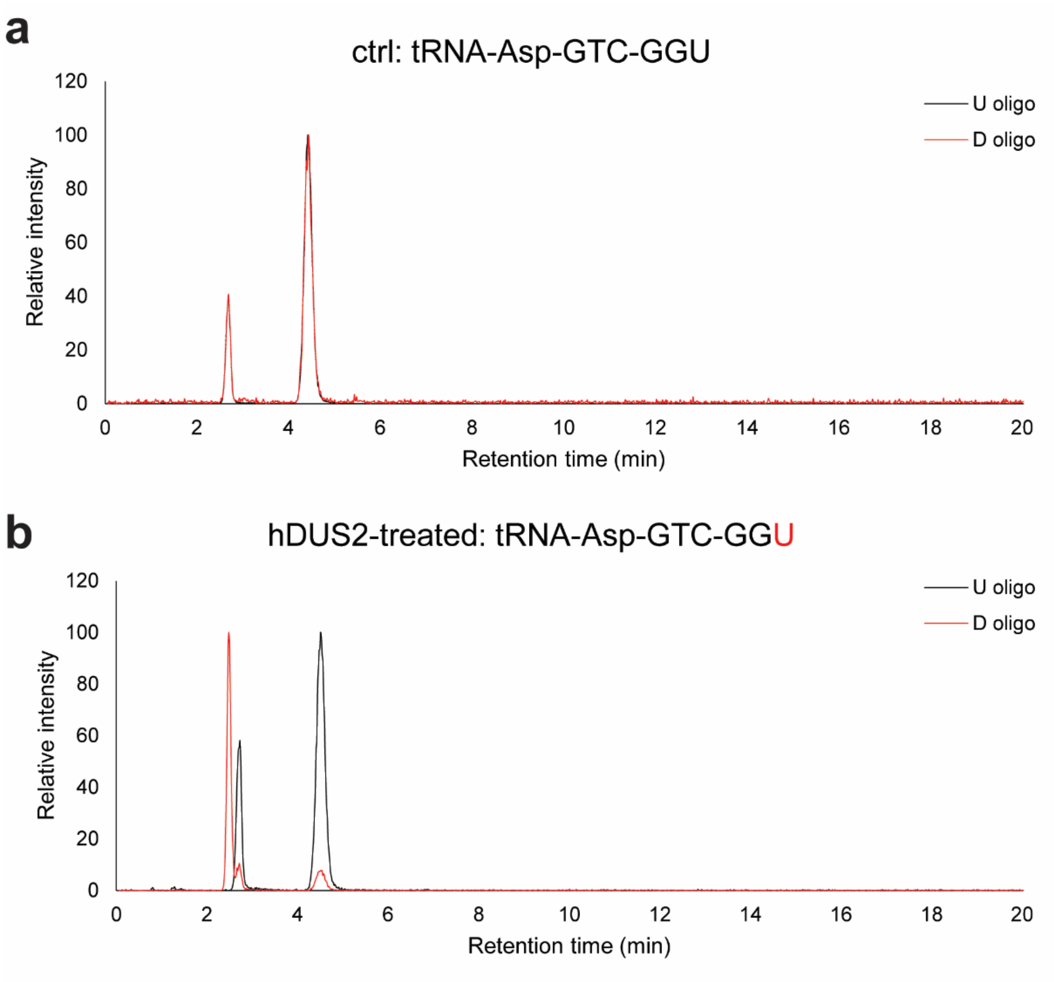


**Supplementary Figure 24.** Oligonucleotide LC-MS analysis of hDUS2-catalyzed dihydrouridine formation on tRNA-Asp-GTC. Extracted ion chromatography of characteristic oligonucleotide fragment from RNase A digestion and QuickCIP dephosphorylation in control tRNA sample **(a)** and hDUS2-treated tRNA **(b)**. U-oligo: $[M-H]^-$ = 933.1662; D-oligo: $[M-H]^-$ = 935.182.

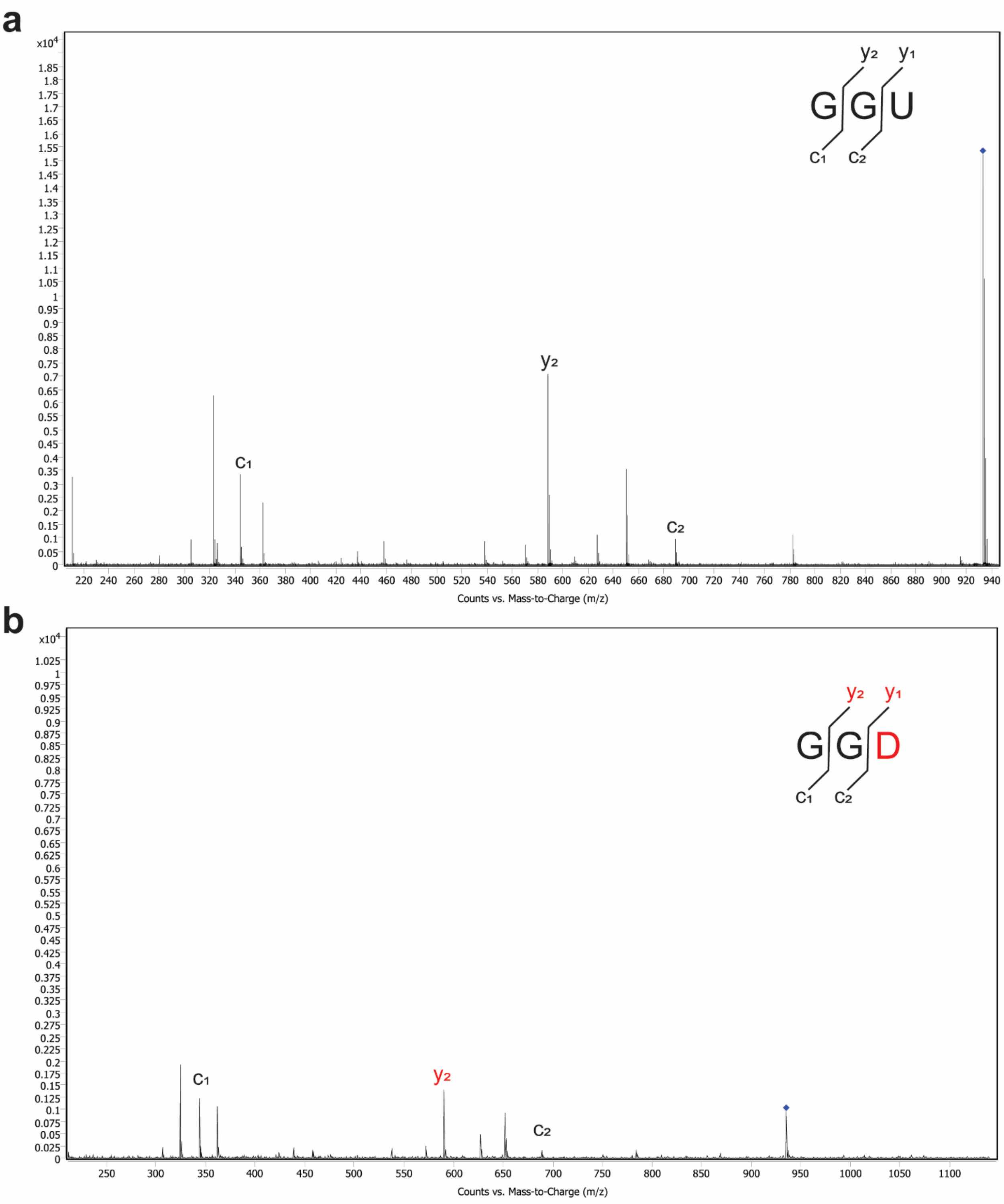


**Supplementary Figure 25.** MS/MS spectrum of GGU and GGD oligo fragments produced from RNase A digestion and QuickCIP dephosphorylation of control tRNA-Asp-GTC sample **(a)** and hDUS2-treated tRNA-Asp-GTC **(b)**. MS/MS fragmentation was performed by collision-induced dissociation. The U-containing precursor ion is $[M\text{-}H]^-$ = 933.1662. The D-containing precursor ion is $[M\text{-}H]^-$ = 935.182.

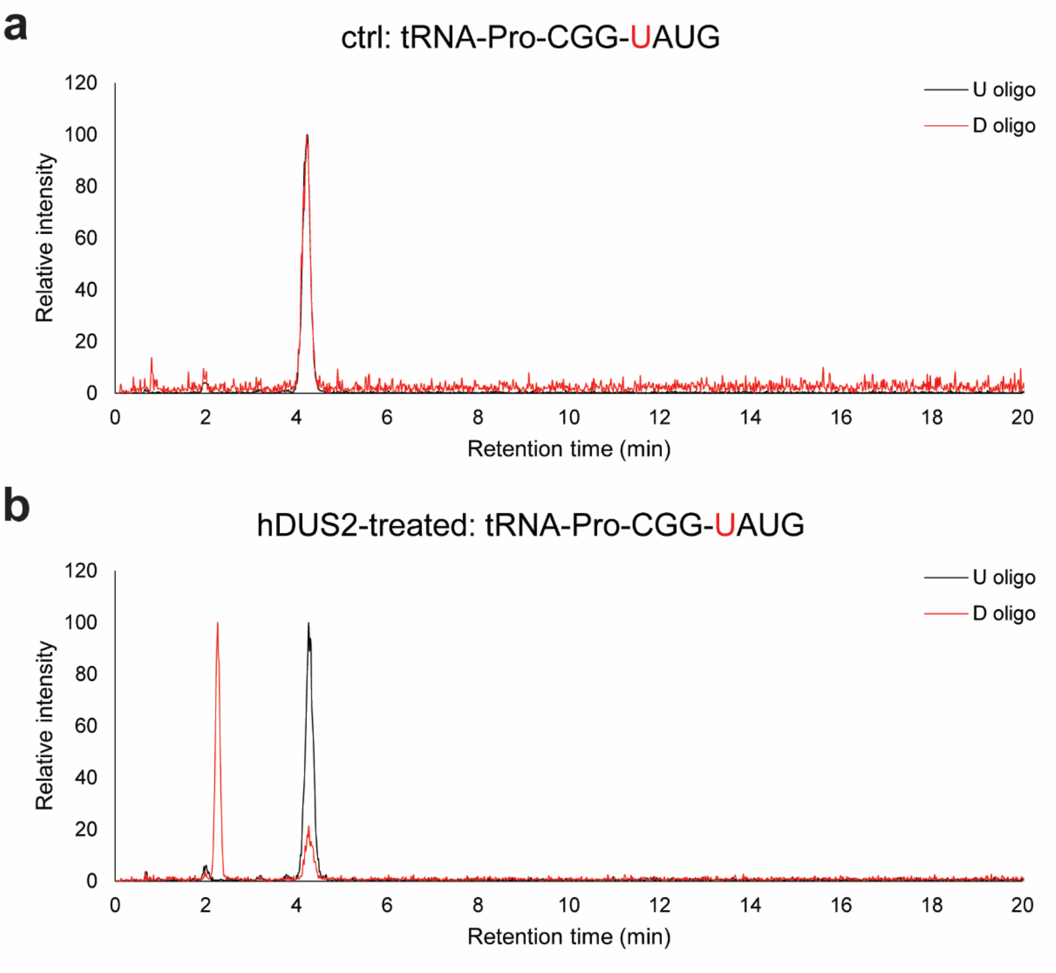


**Supplementary Figure 26.** Oligonucleotide LC-MS analysis of hDUS2-catalyzed dihydrouridine formation on tRNA-Pro-CGG. Extracted ion chromatography of characteristic oligonucleotide fragment from RNase T1 digestion in control tRNA sample **(a)** and hDUS2-treated tRNA **(b)**. U-oligo: $[M-H]^-$ = 1303.165; D-oligo: $[M-H]^-$ = 1305.181.

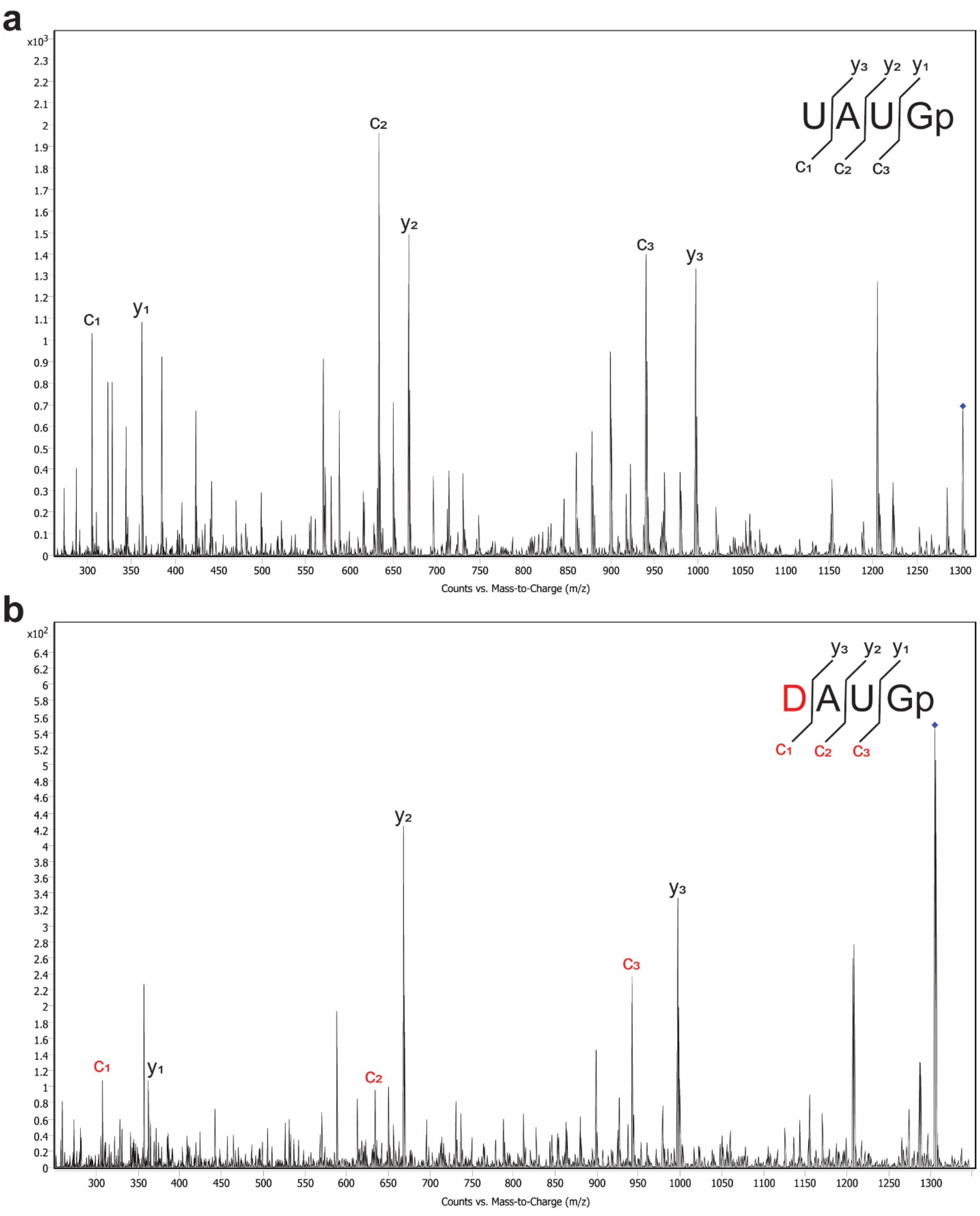


**Supplementary Figure 27.** MS/MS spectrum of UAUGp and DAUGp oligo fragments produced from RNase T1 digestion of control tRNA-Pro-CGG sample **(a)** and hDUS2-treated tRNA-Pro-CGG **(b)**. MS/MS fragmentation was performed by collision-induced dissociation. The U-containing precursor ion is $[M-H]^-$ = 1303.165. The D-containing precursor ion is $[M-H]^-$ = 1305.181.

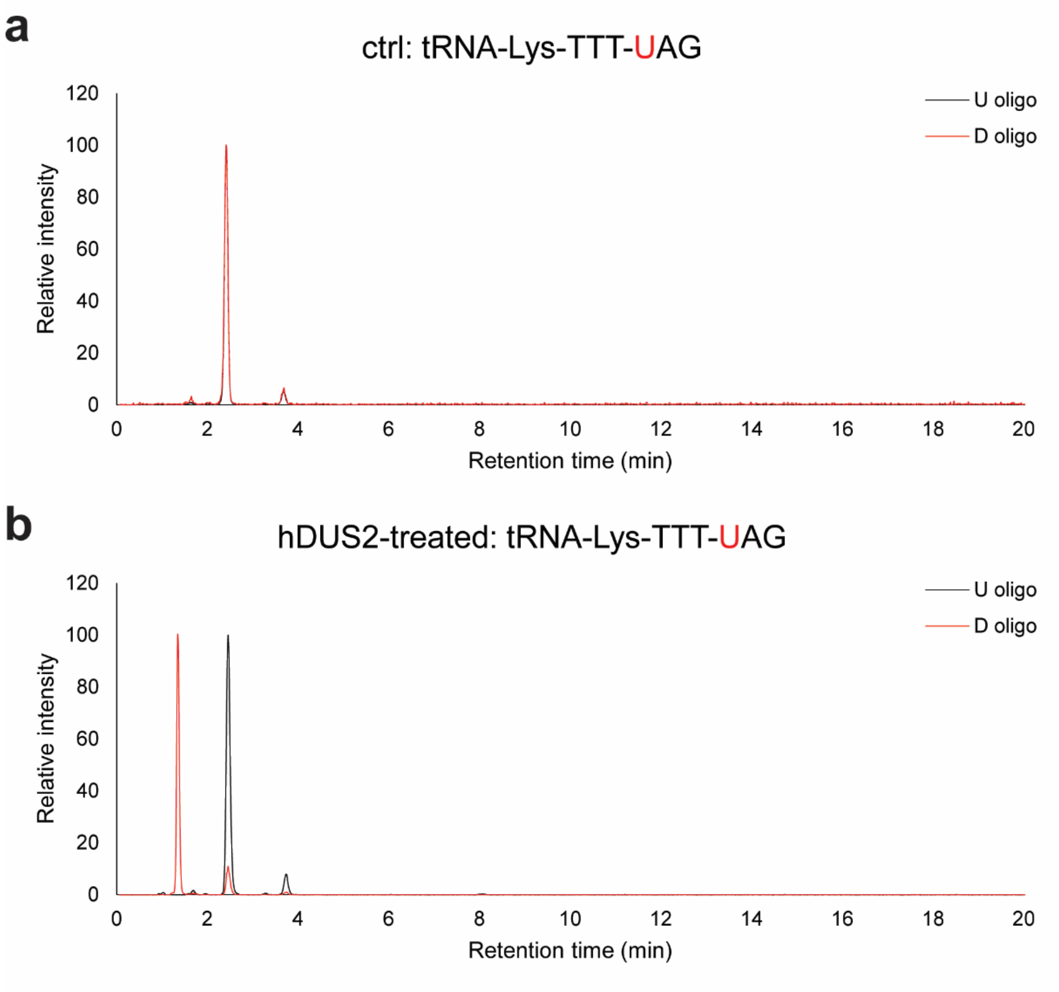


**Supplementary Figure 28.** Oligonucleotide LC-MS analysis of hDUS2-catalyzed dihydrouridine formation on tRNA-Lys-TTT. Extracted ion chromatography of characteristic oligonucleotide fragment from RNase T1 digestion in control tRNA sample **(a)** and hDUS2-treated tRNA **(b)**. U-oligo: $[M-H]^-$ = 997.1404; D-oligo: $[M-H]^-$ = 999.1545.

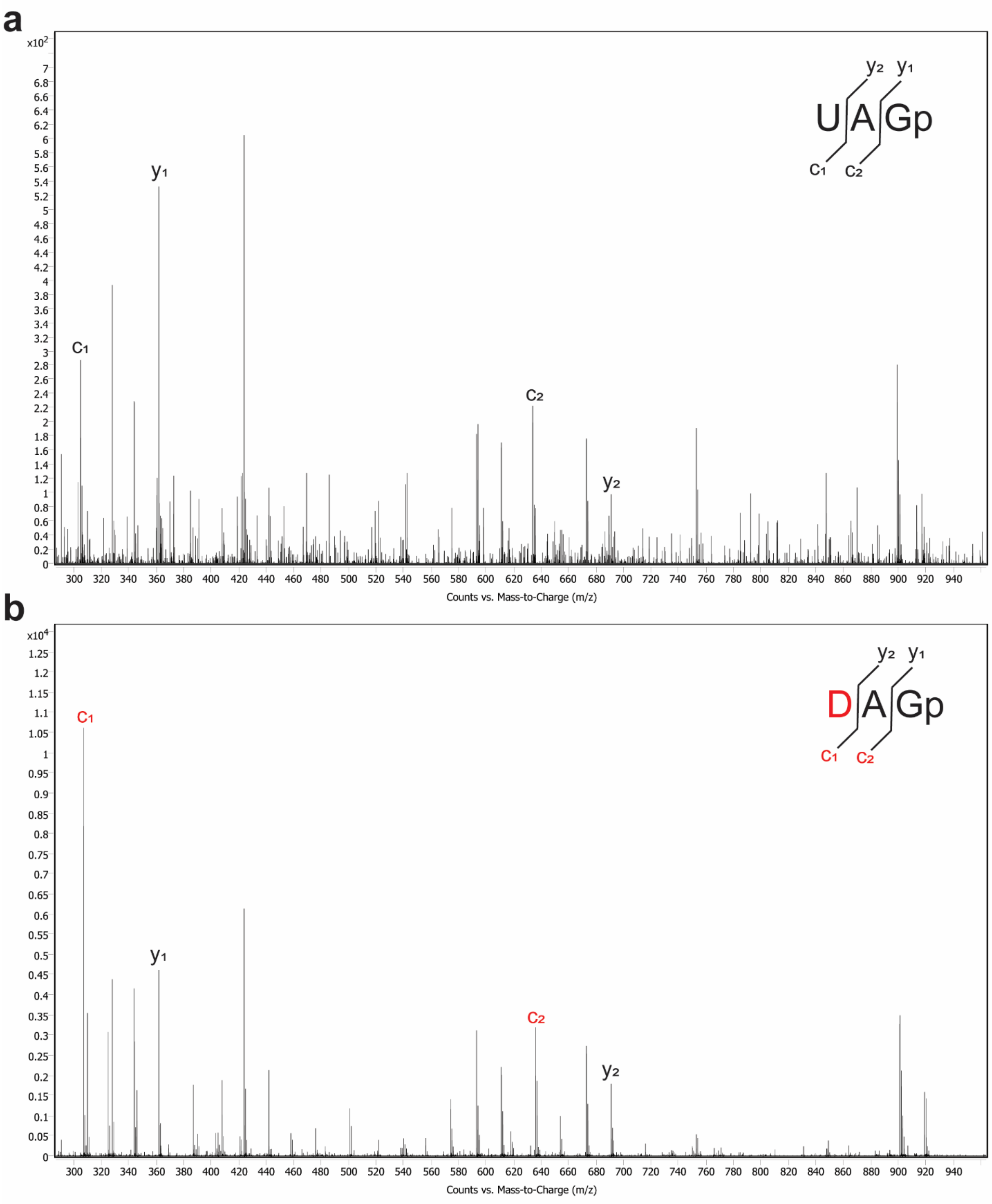


**Supplementary Figure 29.** MS/MS spectrum of UAGp and DAGp oligo fragments produced from RNase T1 digestion of control tRNA-Lys-TTT sample **(a)** and hDUS2-treated tRNA-Lys-TTT **(b)**. MS/MS fragmentation was performed by collision-induced dissociation. The U-containing precursor ion is $[M-H]^-$ = 997.1404. The D-containing precursor ion is $[M-H]^-$ = 999.1545.

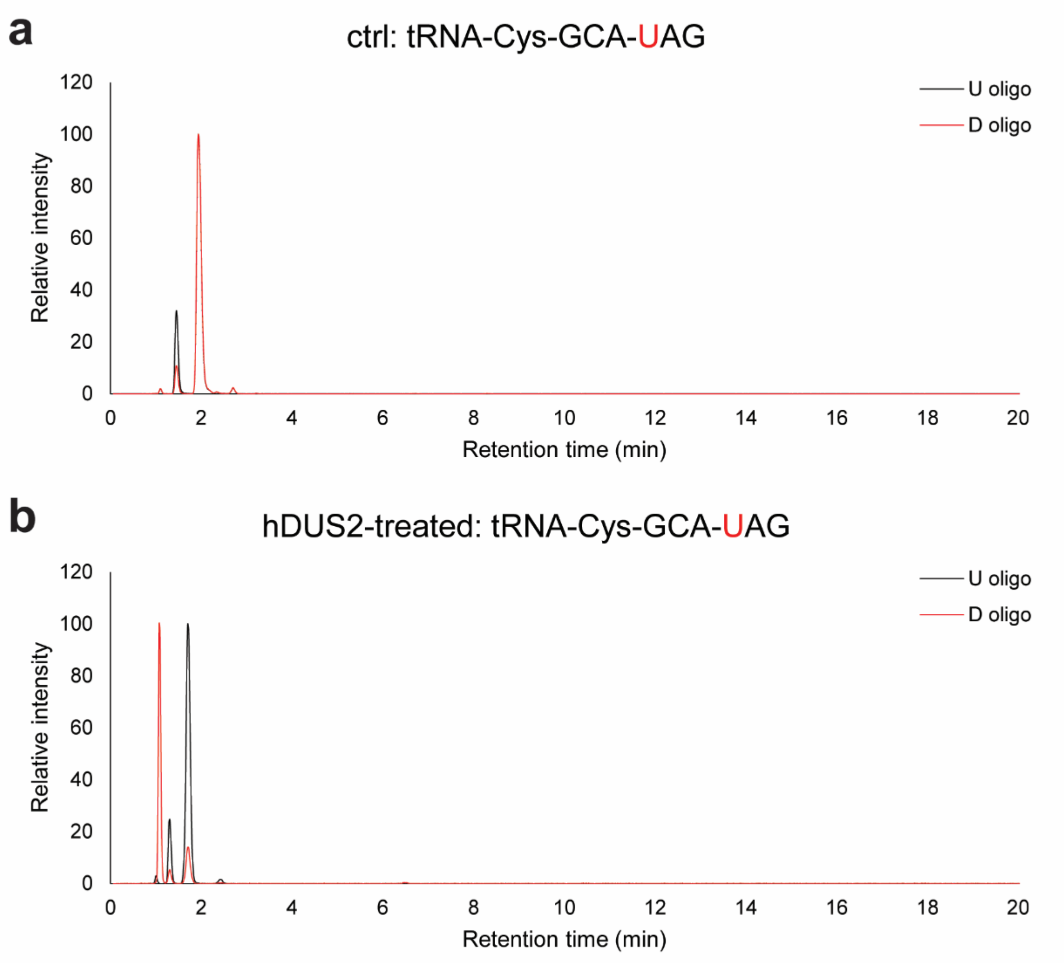


**Supplementary Figure 30.** Oligonucleotide LC-MS analysis of hDUS2-catalyzed dihydrouridine formation on tRNA-Cys-GCA. Extracted ion chromatography of characteristic oligonucleotide fragment from RNase T1 digestion in control tRNA sample **(a)** and hDUS2-treated tRNA **(b)**. U-oligo: $[M\text{-}H]^-$ = 997.1517; D-oligo: $[M\text{-}H]^-$ = 999.1668.

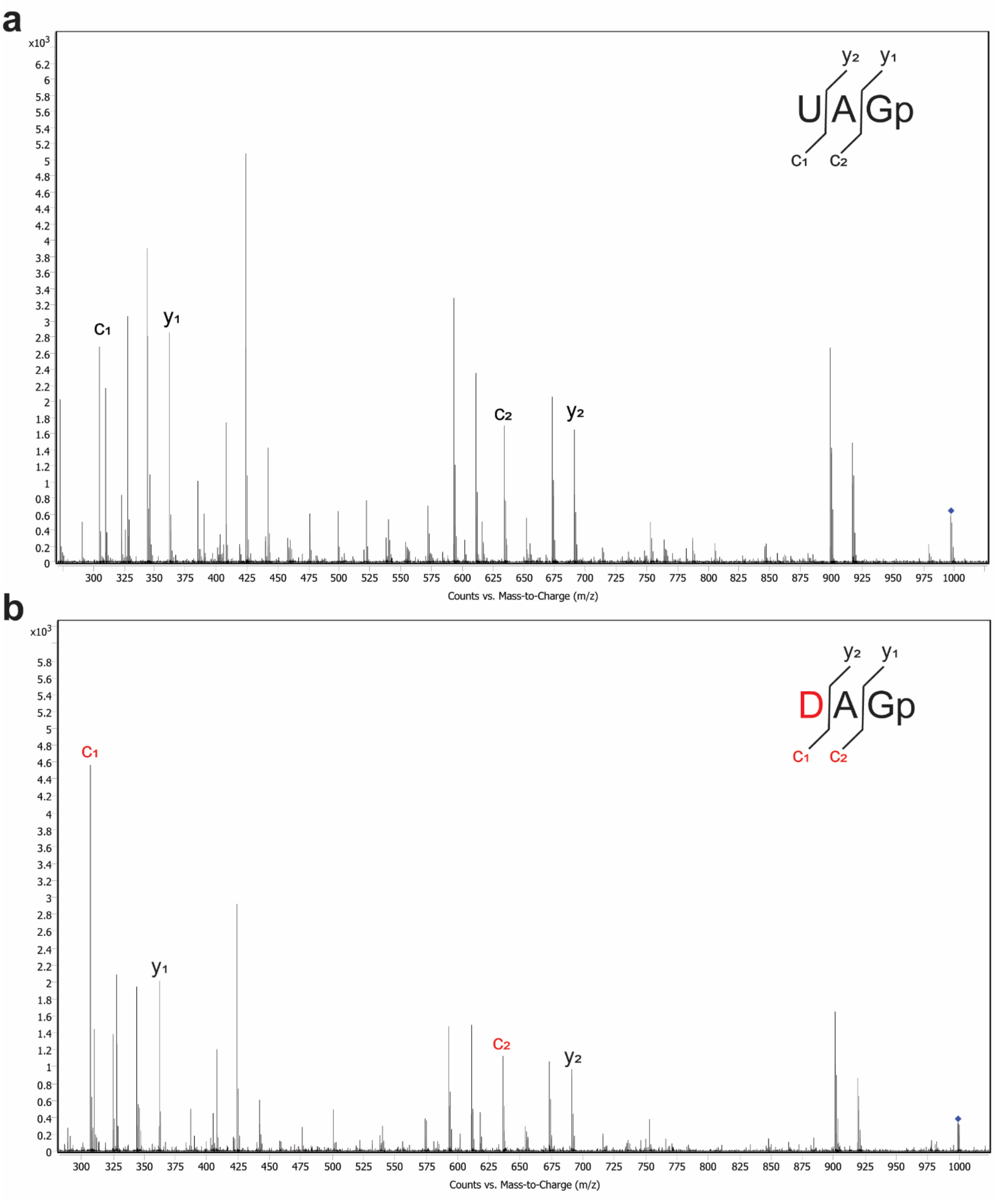


**Supplementary Figure 31.** MS/MS spectrum of UAGp and DAGp oligo fragments produced from RNase T1 digestion of control tRNA-Cys-GCA sample **(a)** and hDUS2-treated tRNA-Cys-GCA **(b)**. MS/MS fragmentation was performed by collision-induced dissociation. The U-containing precursor ion is $[M-H]^-$ = 997.1517. The D-containing precursor ion is $[M-H]^-$ = 999.1668.

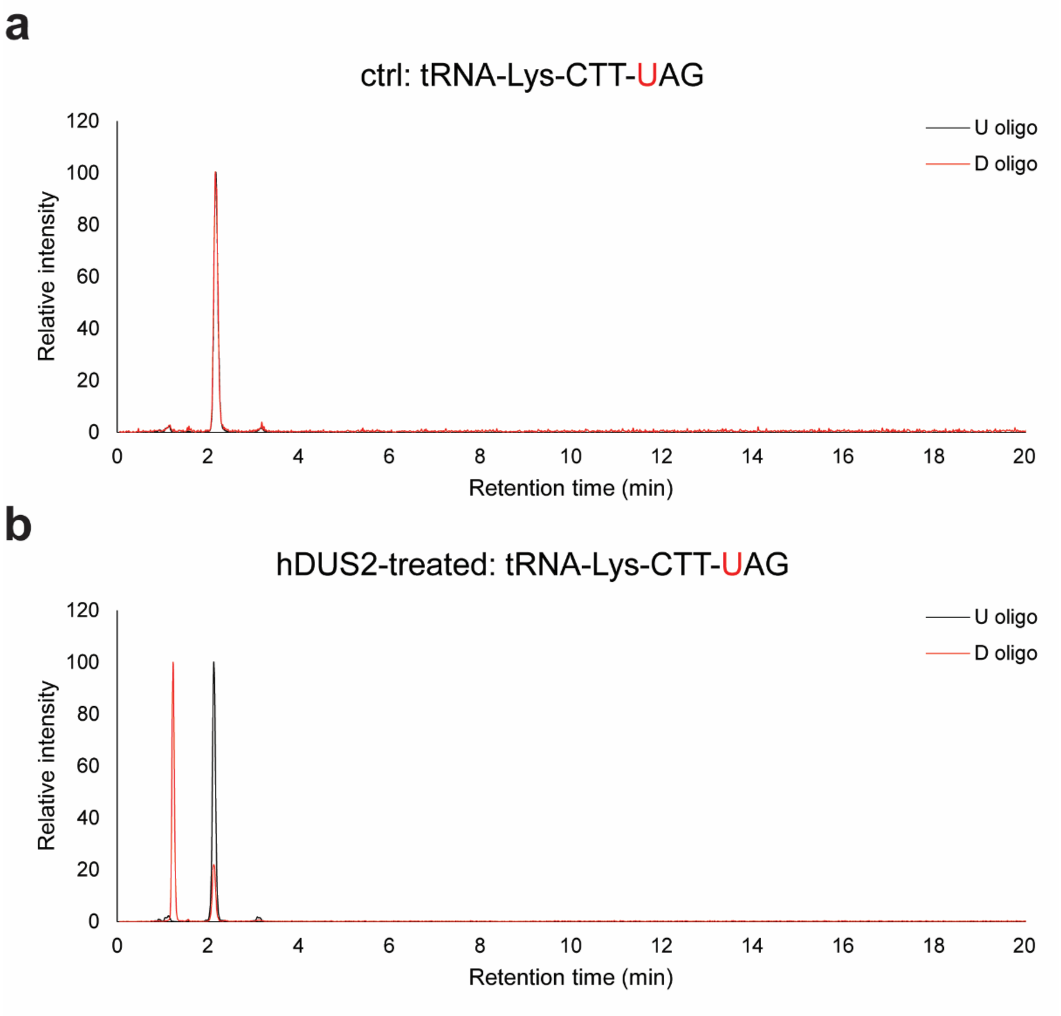


**Supplementary Figure 32.** Oligonucleotide LC-MS analysis of hDUS2-catalyzed dihydrouridine formation on tRNA-Lys-CTT. Extracted ion chromatography of characteristic oligonucleotide fragment from RNase T1 digestion in control tRNA sample **(a)** and hDUS2-treated tRNA **(b)**. U-oligo: $[M\text{-}H]^-$ = 997.144; D-oligo: $[M\text{-}H]^-$ = 999.1591.

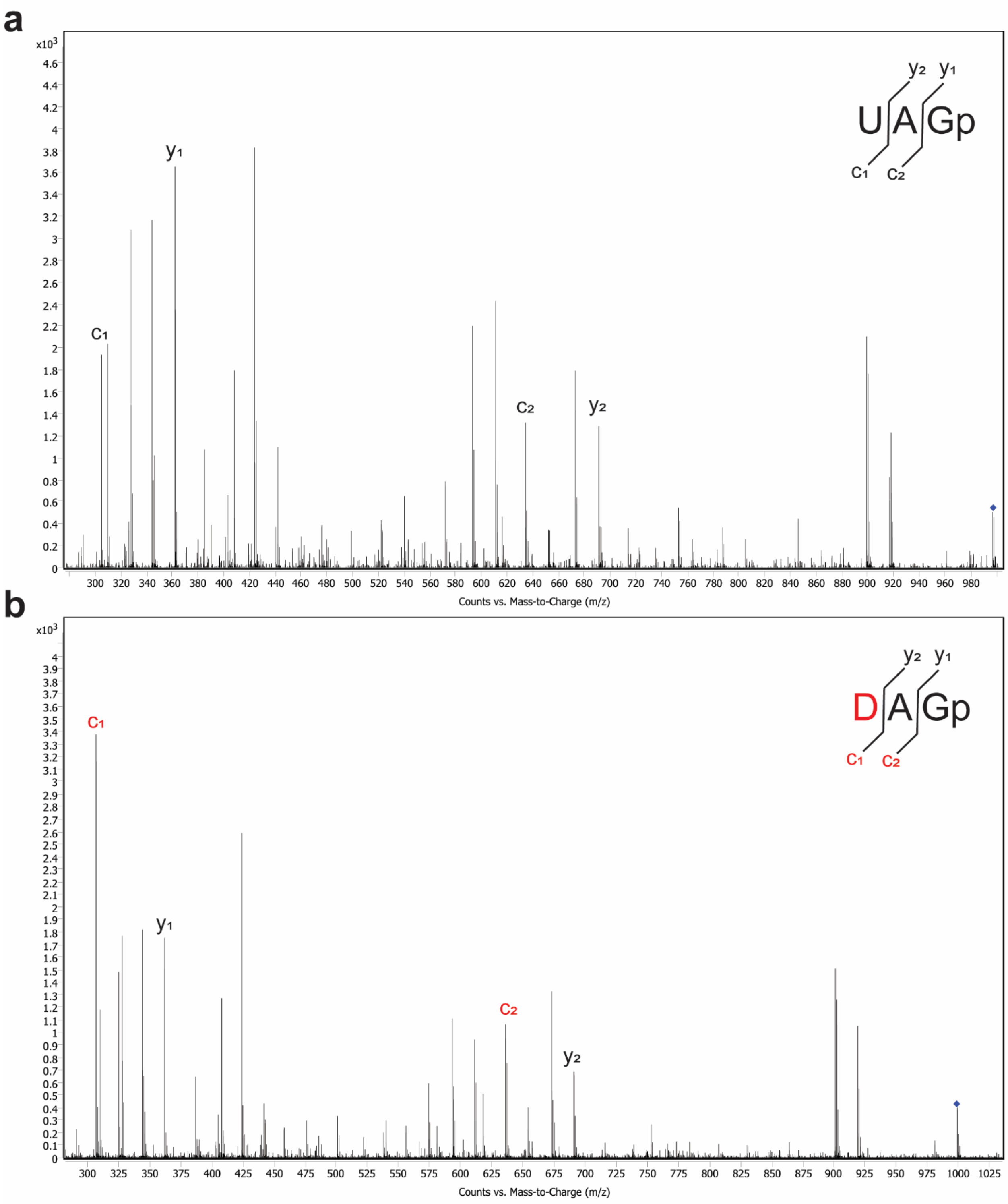


**Supplementary Figure 33.** MS/MS spectrum of UAGp and DAGp oligo fragments produced from RNase T1 digestion of control tRNA-Lys-CTT sample **(a)** and hDUS2-treated tRNA-Lys-CTT **(b)**. MS/MS fragmentation was performed by collision-induced dissociation. The U-containing precursor ion is $[M-H]^-$ = 997.144. The D-containing precursor ion is $[M-H]^-$ = 999.1591.

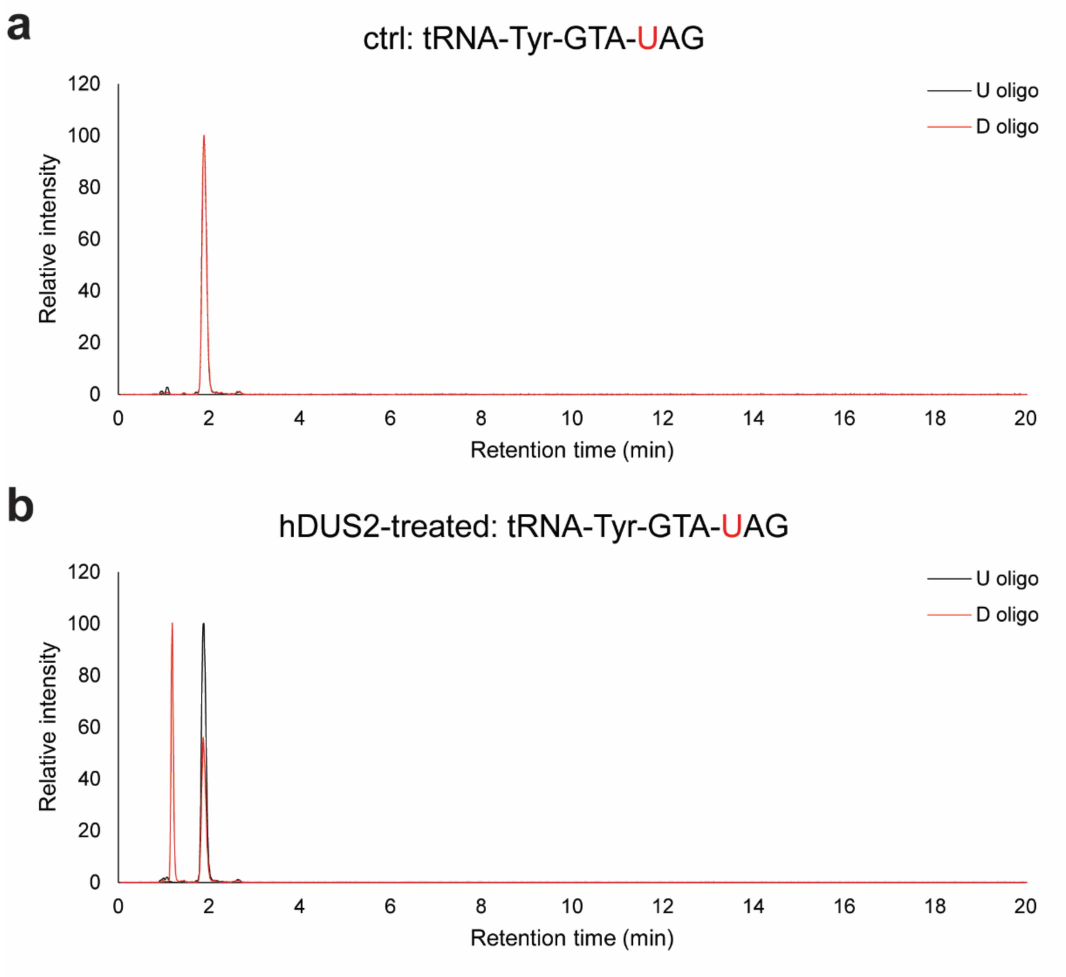


**Supplementary Figure 34.** Oligonucleotide LC-MS analysis of hDUS2-catalyzed dihydrouridine formation on tRNA-Tyr-GTA. Extracted ion chromatography of characteristic oligonucleotide fragment from RNase T1 digestion in control tRNA sample **(a)** and hDUS2-treated tRNA **(b)**. U-oligo: $[M-H]^-$ = 997.1404; D-oligo: $[M-H]^-$ = 999.1545.

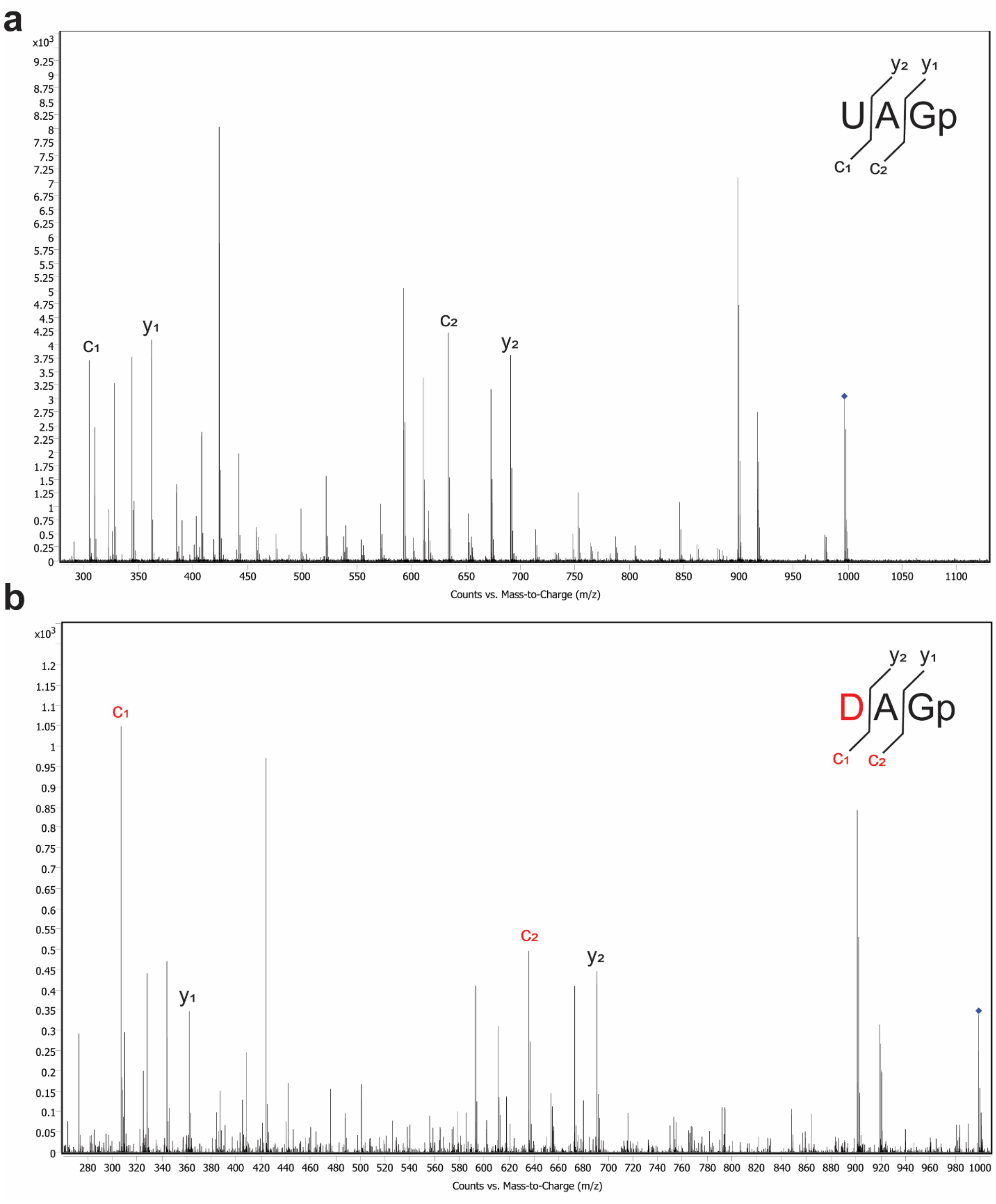


**Supplementary Figure 35.** MS/MS spectrum of UAGp and DAGp oligo fragments produced from RNase T1 digestion of control tRNA-Tyr-GTA sample **(a)** and hDUS2-treated tRNA-Tyr-GTA **(b)**. MS/MS fragmentation was performed by collision-induced dissociation. The U-containing precursor ion is $[M\text{-}H]^-$ = 997.1404. The D-containing precursor ion is $[M\text{-}H]^-$ = 999.1545.

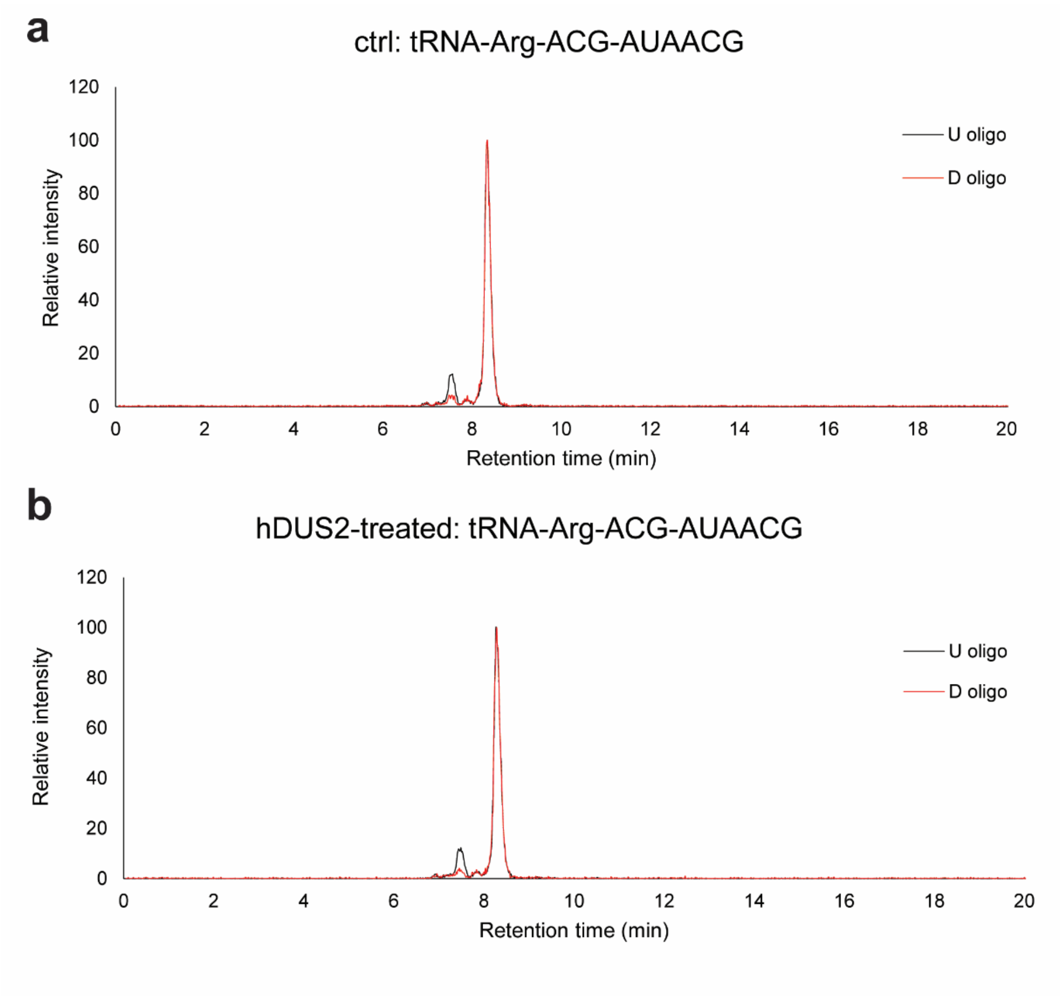


**Supplementary Figure 36.** Oligonucleotide LC-MS analysis of hDUS2-catalyzed dihydrouridine formation on tRNA-Arg-ACG. Extracted ion chromatography of characteristic oligonucleotide fragment from RNase T1 digestion in control tRNA sample **(a)** and hDUS2-treated tRNA **(b)**. U-oligo: $[M\text{-}2H]^{2-}$ = 979.632; D-oligo: $[M\text{-}2H]^{2-}$ = 980.632.

**a**

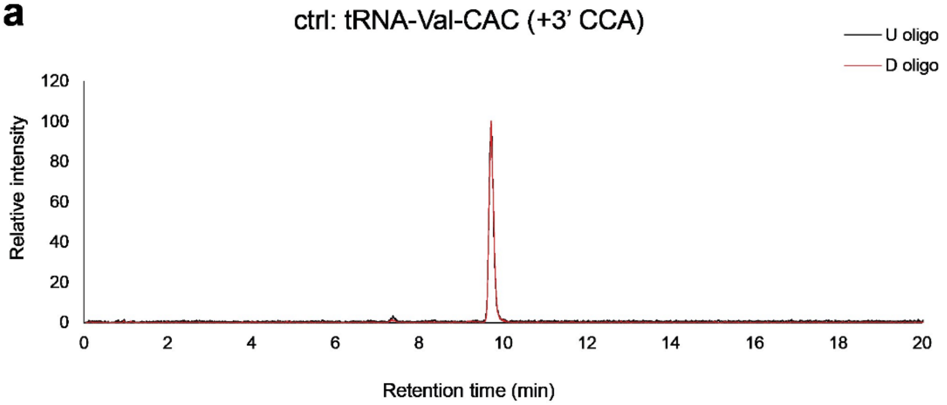


**b**

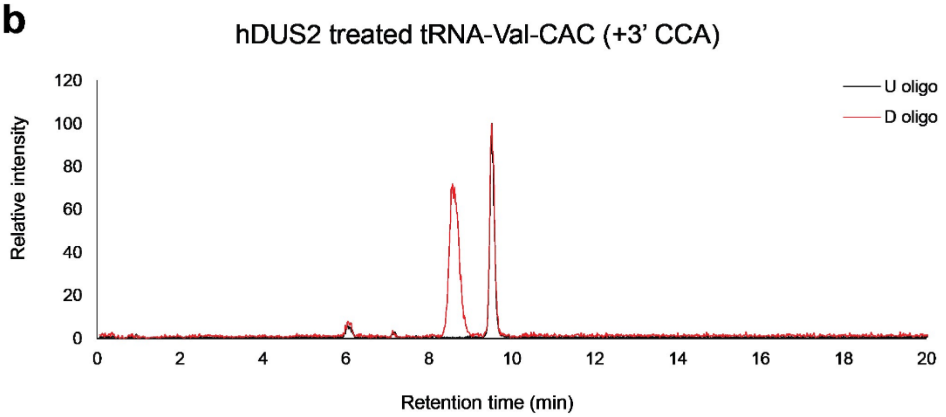


**Supplementary Figure 37.** Oligonucleotide LC-MS analysis of hDUS2-catalyzed dihydrouridine formation on tRNA-Val-CAC with 3' CCA. Extracted ion chromatography of characteristic oligonucleotide fragment from RNase T1 digestion in control tRNA sample **(a)** and hDUS2-treated tRNA **(b)**. U-oligo: $[M-2H]^{2-}$ = 1273.669; D-oligo: $[M-2H]^{2-}$ = 1274.677.

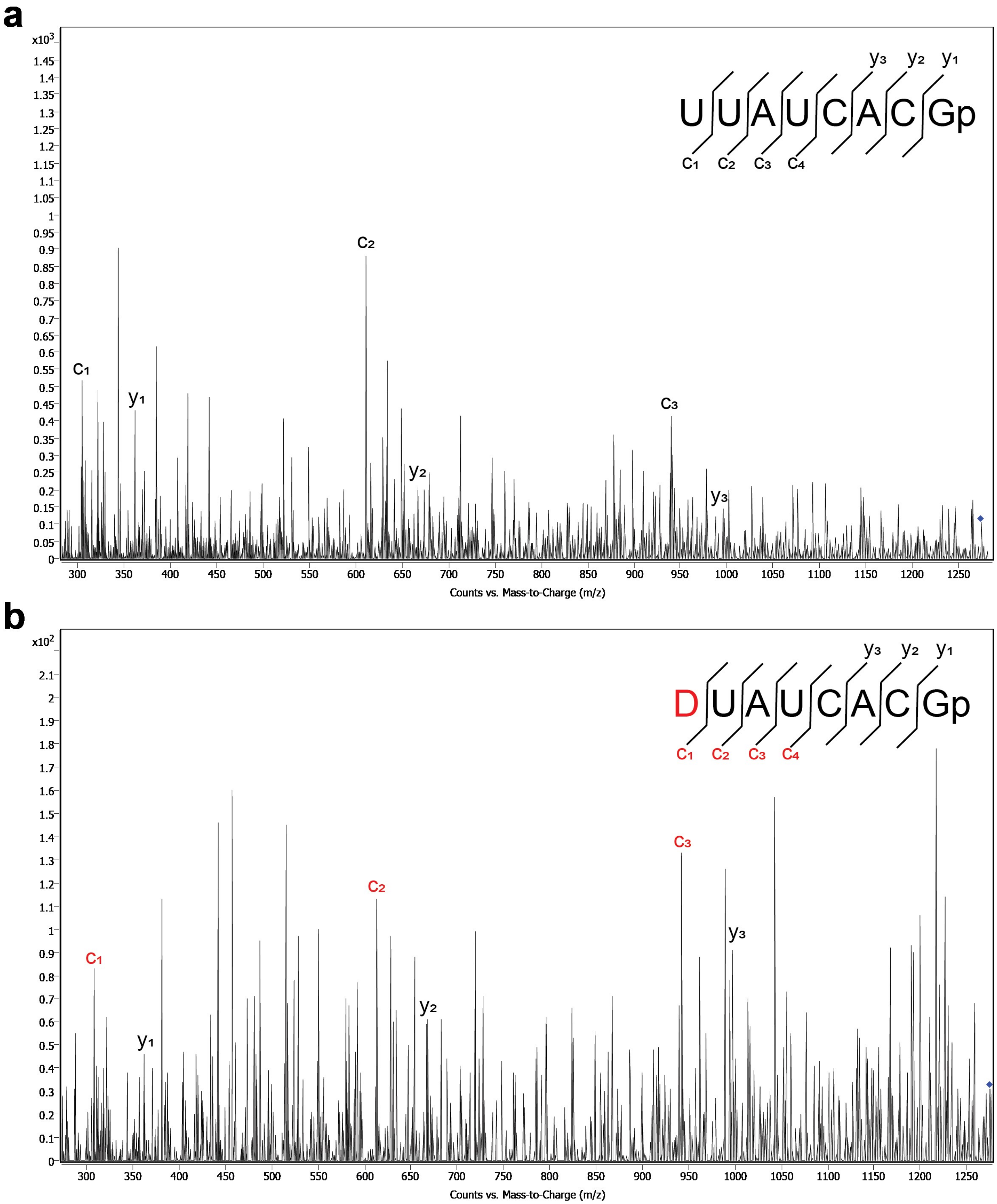


**Supplementary Figure 38.** MS/MS spectrum of UUAUCACGp and DUAUCACGp oligo fragments produced from RNase T1 digestion of control tRNA-Val-CAC (+3'CCA) sample **(a)** and hDUS2-treated tRNA-Val-CAC (+3'CCA) **(b)**. MS/MS fragmentation was performed by collision-induced dissociation. The U-containing precursor ion is $[M\text{-}2H]^{2-}$ = 1273.669. The D-containing precursor ion is $[M\text{-}2H]^{2-}$ = 1274.677.

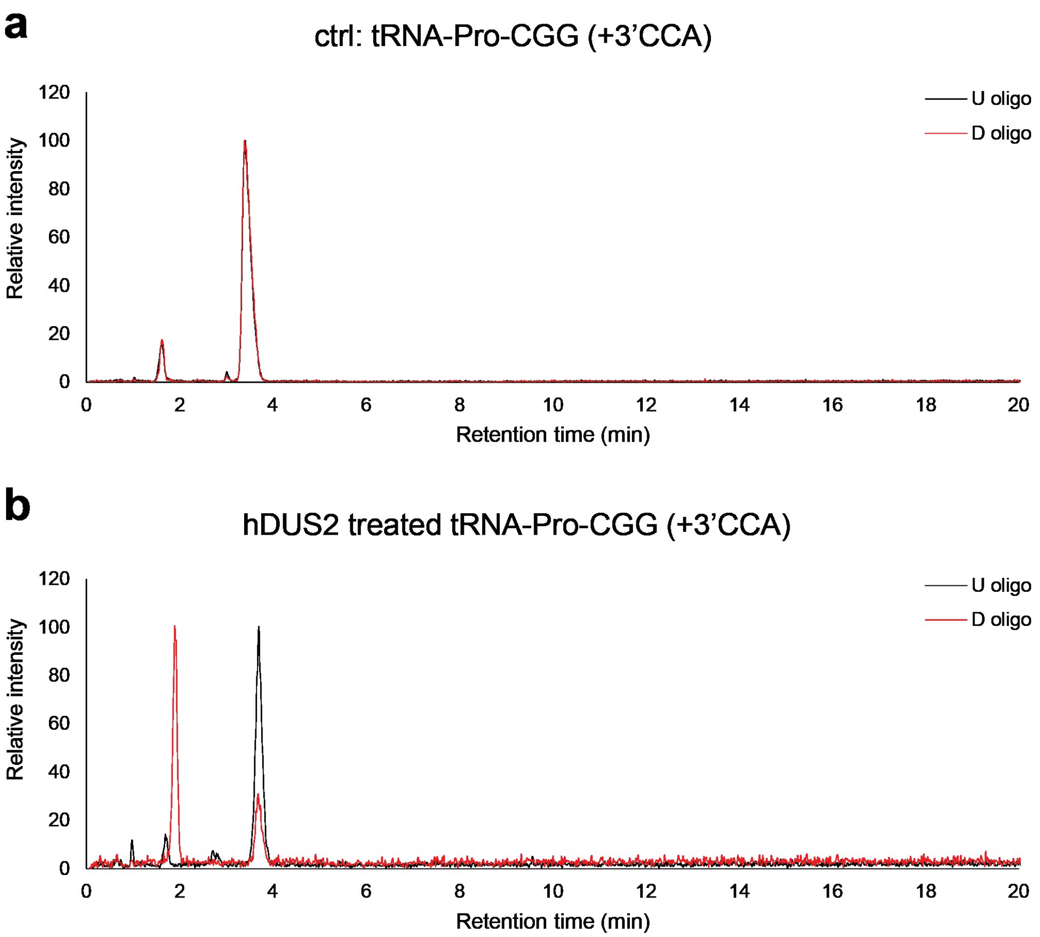


**Supplementary Figure 39.** Oligonucleotide LC-MS analysis of hDUS2-catalyzed dihydrouridine formation on tRNA-Pro-CGG (+3'CCA). Extracted ion chromatography of characteristic oligonucleotide fragment from RNase T1 digestion in control tRNA sample **(a)** and hDUS2-treated tRNA **(b)**. U-oligo: $[M-H]^-$ = 1303.165; D-oligo: $[M-H]^-$ = 1305.181.

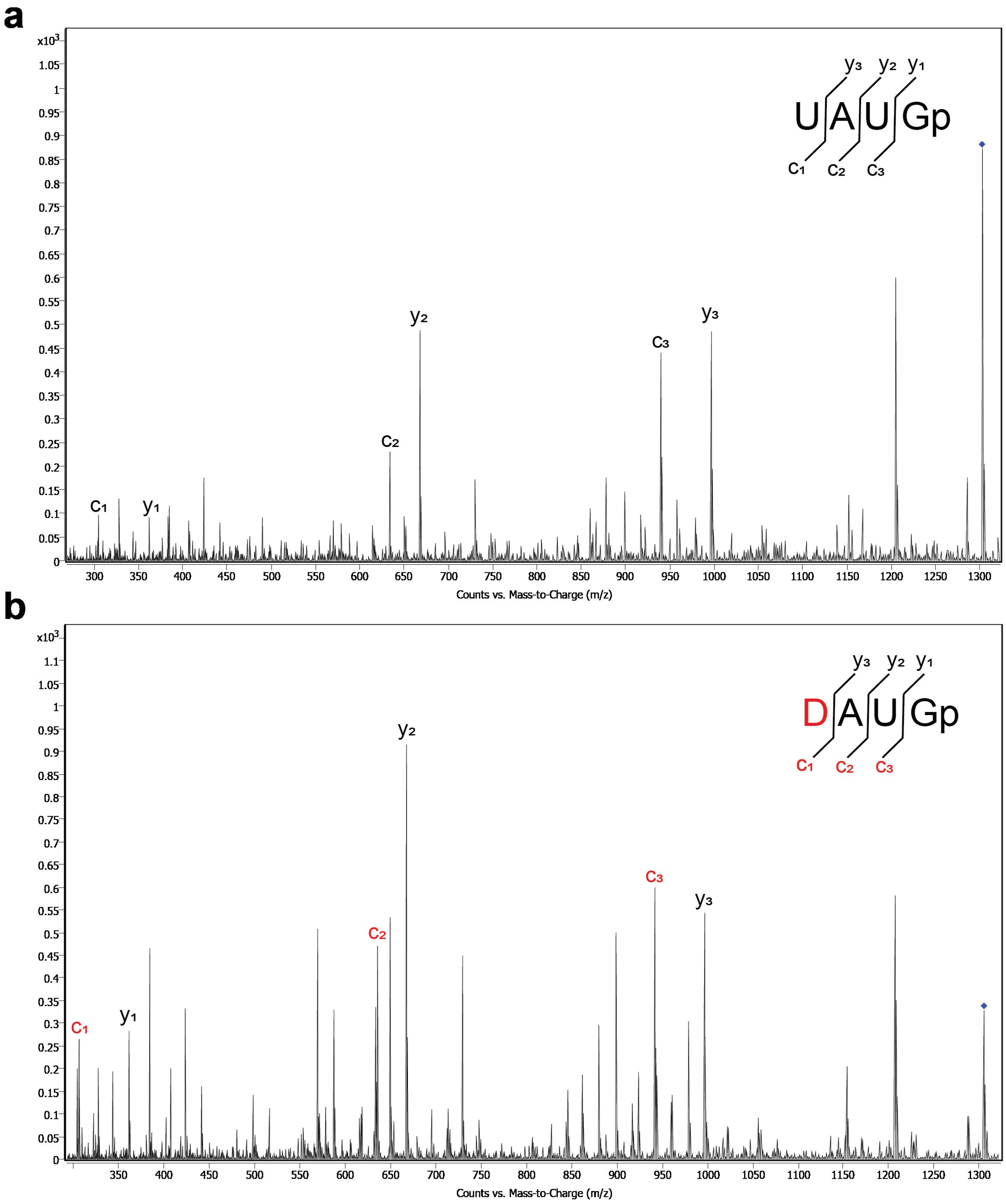


**Supplementary Figure 40.** MS/MS spectrum of UAUGp and DAUGp oligo fragments produced from RNase T1 digestion of control tRNA-Pro-CGG (+3'CCA) sample **(a)** and hDUS2-treated tRNA-Pro-CGG (+3'CCA) **(b)**. MS/MS fragmentation was performed by collision-induced dissociation. The U-containing precursor ion is $[M-H]^-$ = 1303.165. The D-containing precursor ion is $[M-H]^-$ = 1305.181.

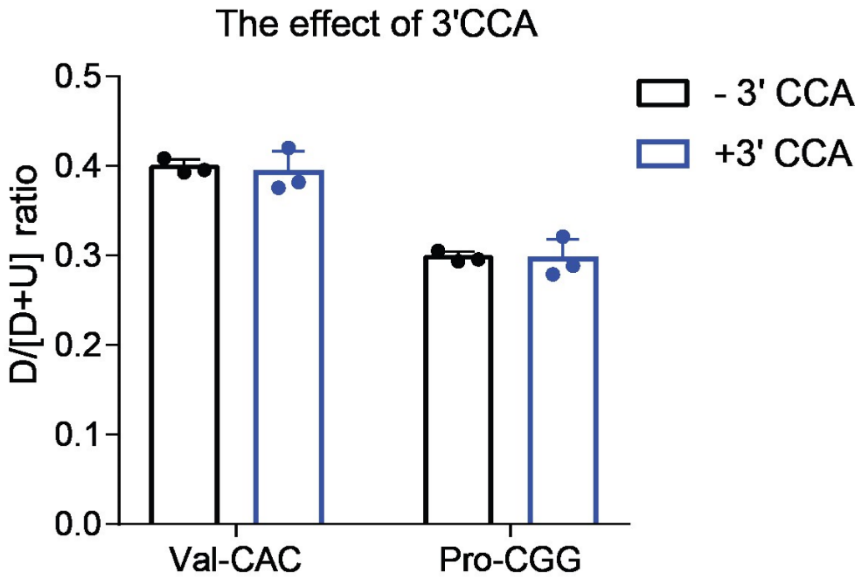


**Supplementary Figure 41.** Oligonucleotide LC-MS quantification of D formation on IVT tRNA with and without 3'CCA. Data was collected from Supplementary Fig. 22, 26, 37, and 39. The D/[D+U] ratio was calculated based upon D and U fragment intensities according to the following formula: (D oligo intensity)/(D oligo intensity + U oligo intensity). Values represent mean ± s.d (n=3).

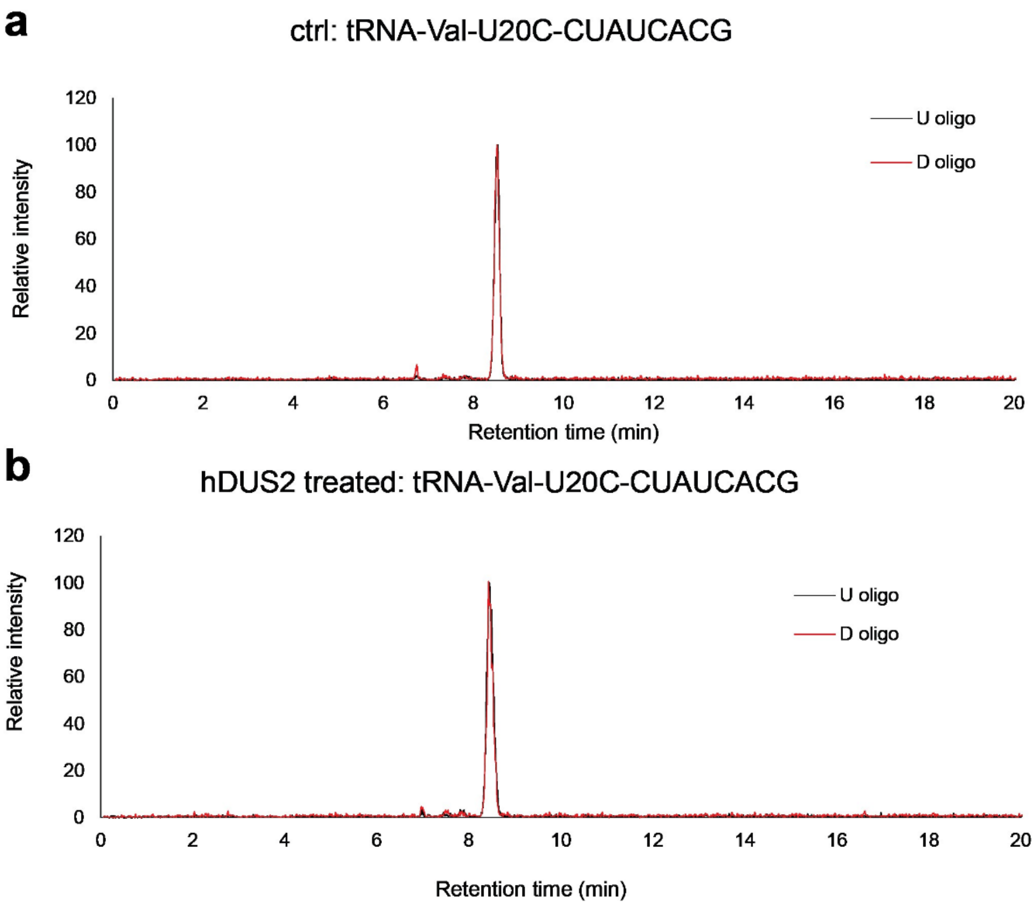


**Supplementary Figure 42.** Oligonucleotide LC-MS analysis of hDUS2-catalyzed dihydrouridine formation on tRNA-Val-CAC-U20C. Extracted ion chromatography of characteristic oligonucleotide fragment from RNase T1 digestion in control tRNA sample **(a)** and hDUS2-treated tRNA **(b)**. U-oligo: $[M-2H]^{2-}$ = 1273.159; D-oligo: $[M-2H]^{2-}$ = 1274.159.

**a**

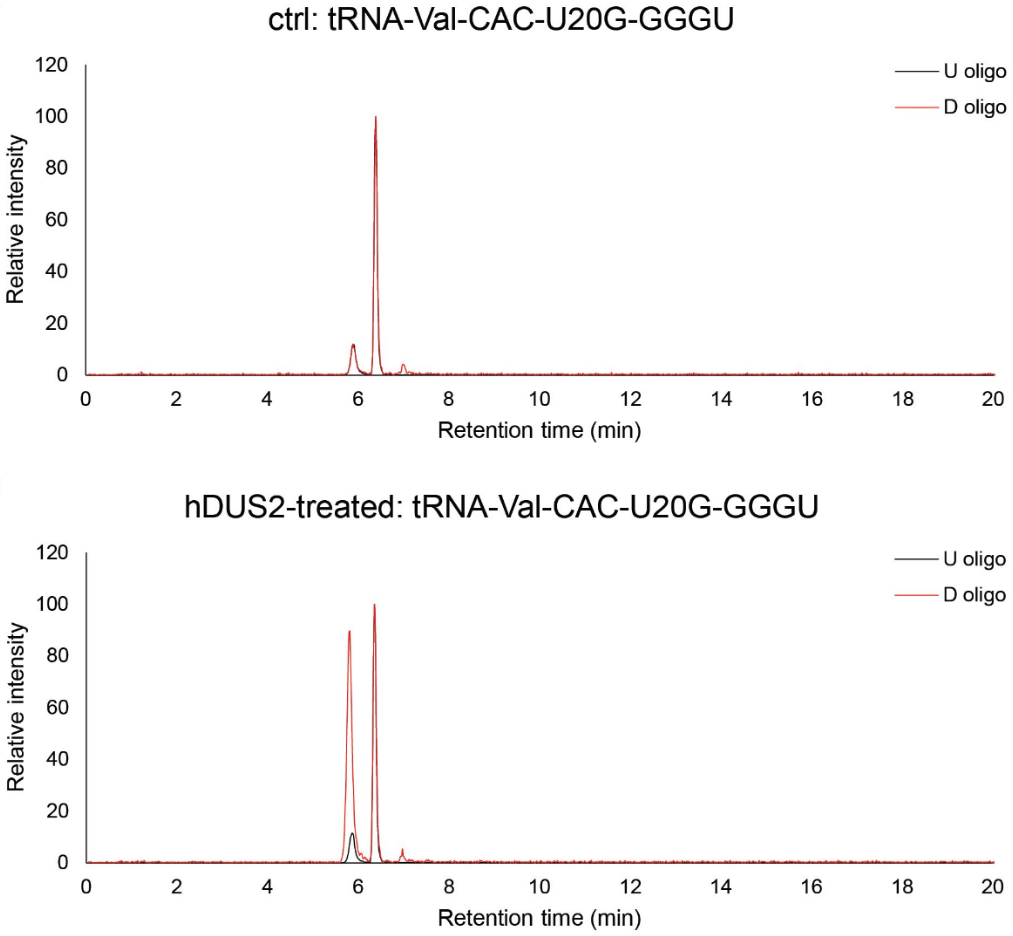


**Supplementary Figure 43.** Oligonucleotide LC-MS analysis of hDUS2-catalyzed dihydrouridine formation on tRNA-Val-CAC-U20G. Extracted ion chromatography of characteristic oligonucleotide fragment from RNase A digestion and QuickCIP dephosphorylation in control tRNA sample **(a)** and hDUS2-treated tRNA **(b)**. U-oligo: $[M-H]^-$ = 1278.233; D-oligo: $[M-H]^-$ = 1280.238.

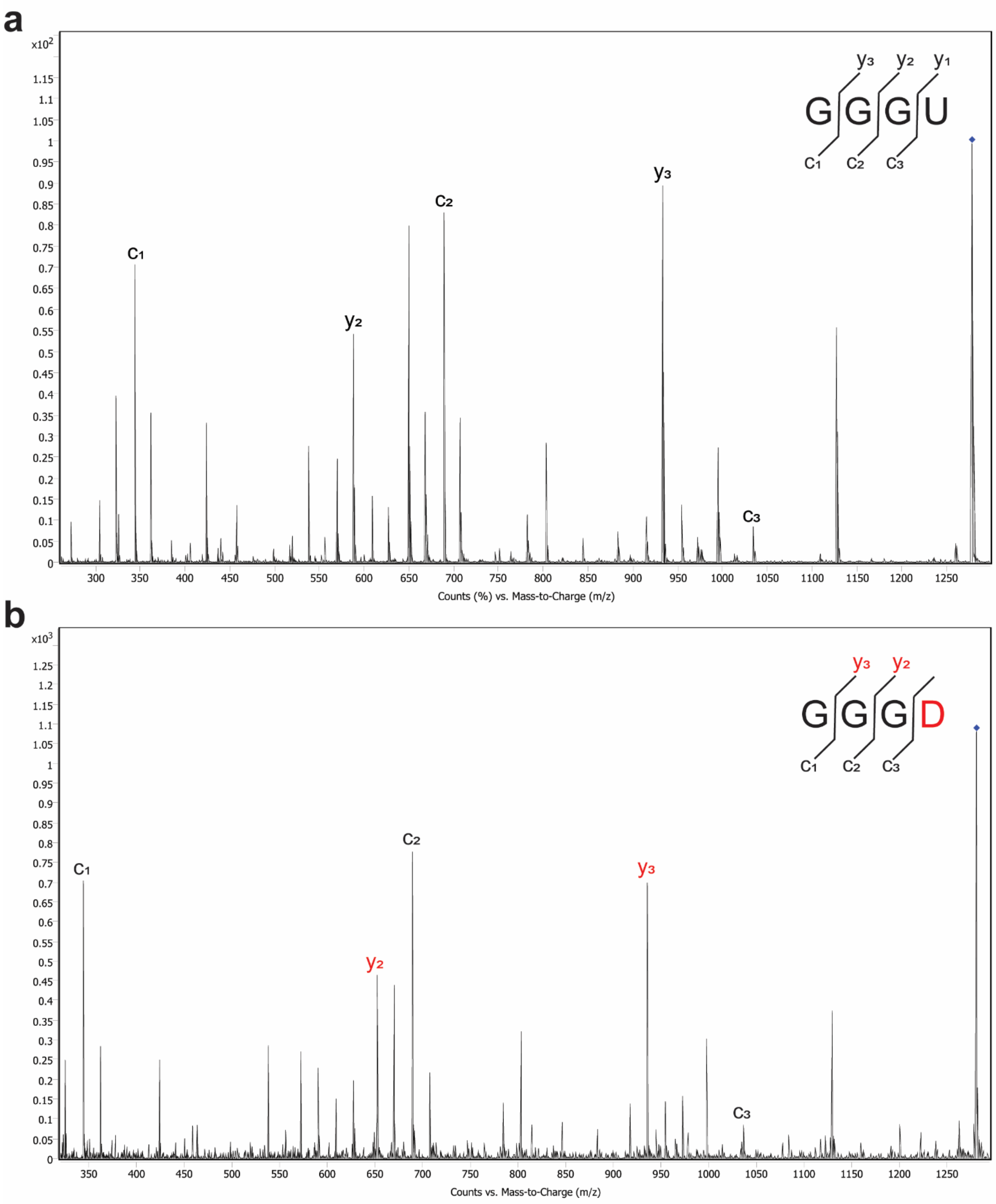


**Supplementary Figure 44.** MS/MS spectrum of GGGU and GGGD oligo fragments produced from RNase A digestion and QuickCIP dephosphorylation of control tRNA-Val-CAC-U20G sample **(a)** and hDUS2-treated tRNA-Val-CAC-U20G **(b)**. MS/MS fragmentation was performed by collision-induced dissociation. The U-containing precursor ion is $[M-H]^-$ = 1278.233. The D-containing precursor ion is $[M-H]^-$ = 1280.238.

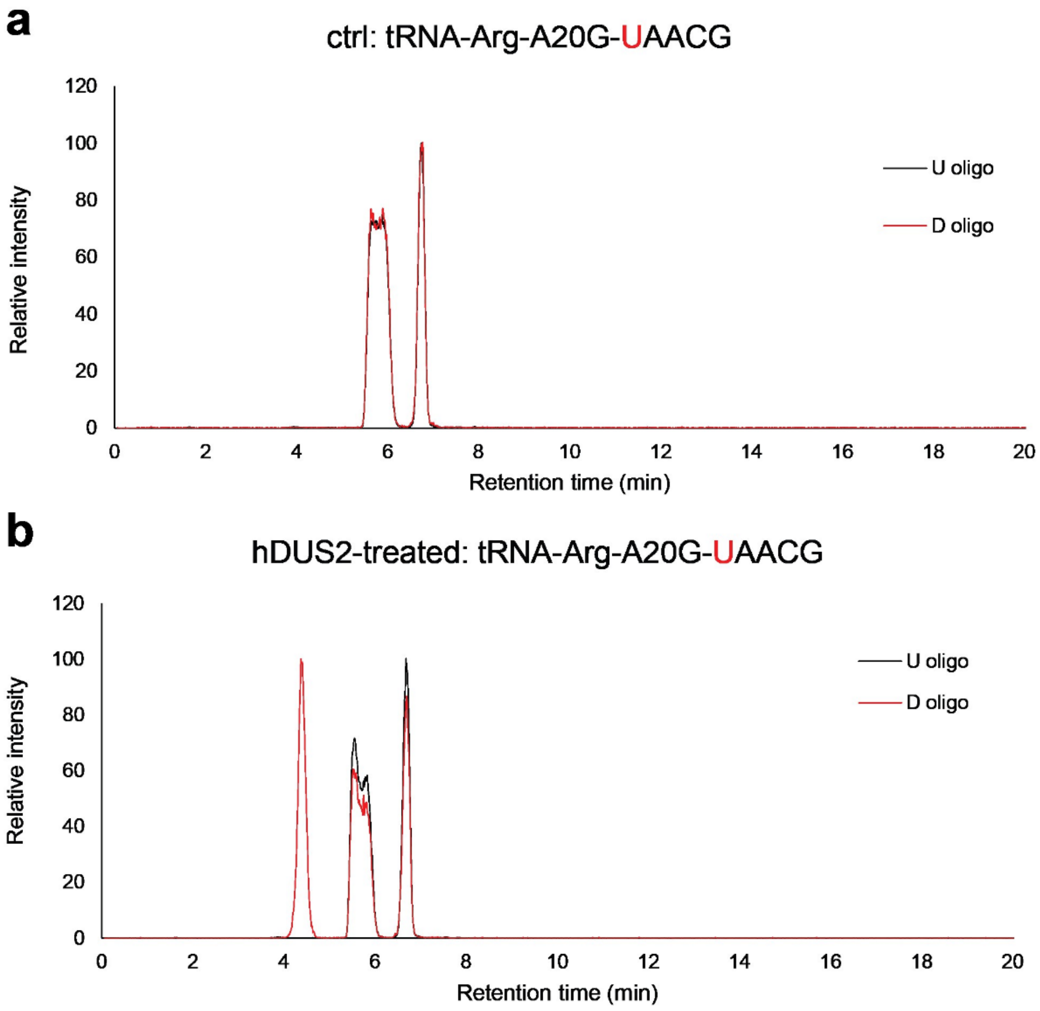


**Supplementary Figure 45.** Oligonucleotide LC-MS analysis of hDUS2-catalyzed dihydrouridine formation on tRNA-Arg-ACG-A20G. Extracted ion chromatography of characteristic oligonucleotide fragment from RNase T1 digestion in control tRNA sample **(a)** and hDUS2-treated tRNA **(b)**. U-oligo: $[M-2H]^{2-}$ = 815.1094; D-oligo: $[M-2H]^{2-}$ = 816.1172.

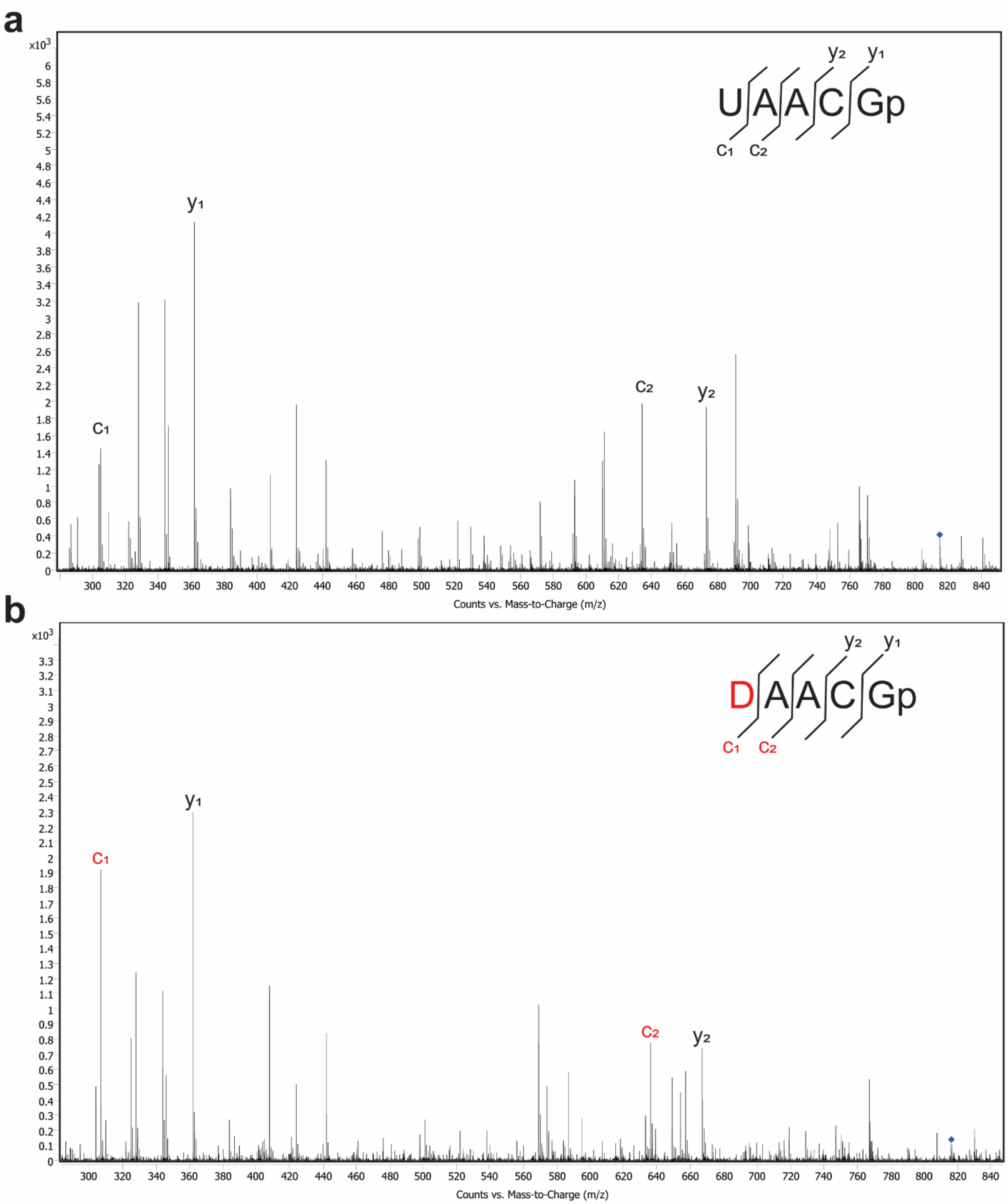


**Supplementary Figure 46.** MS/MS spectrum of UAACGp and DAACGp oligo fragments produced from RNase T1 digestion of control tRNA-Arg-ACG-A20G sample **(a)** and hDUS2-treated tRNA-Arg-ACG-A20G **(b)**. MS/MS fragmentation was performed by collision-induced dissociation. The U-containing precursor ion is $[M-2H]^{2-}$ = 815.1094. The D-containing precursor ion is $[M-2H]^{2-}$ = 816.1172.

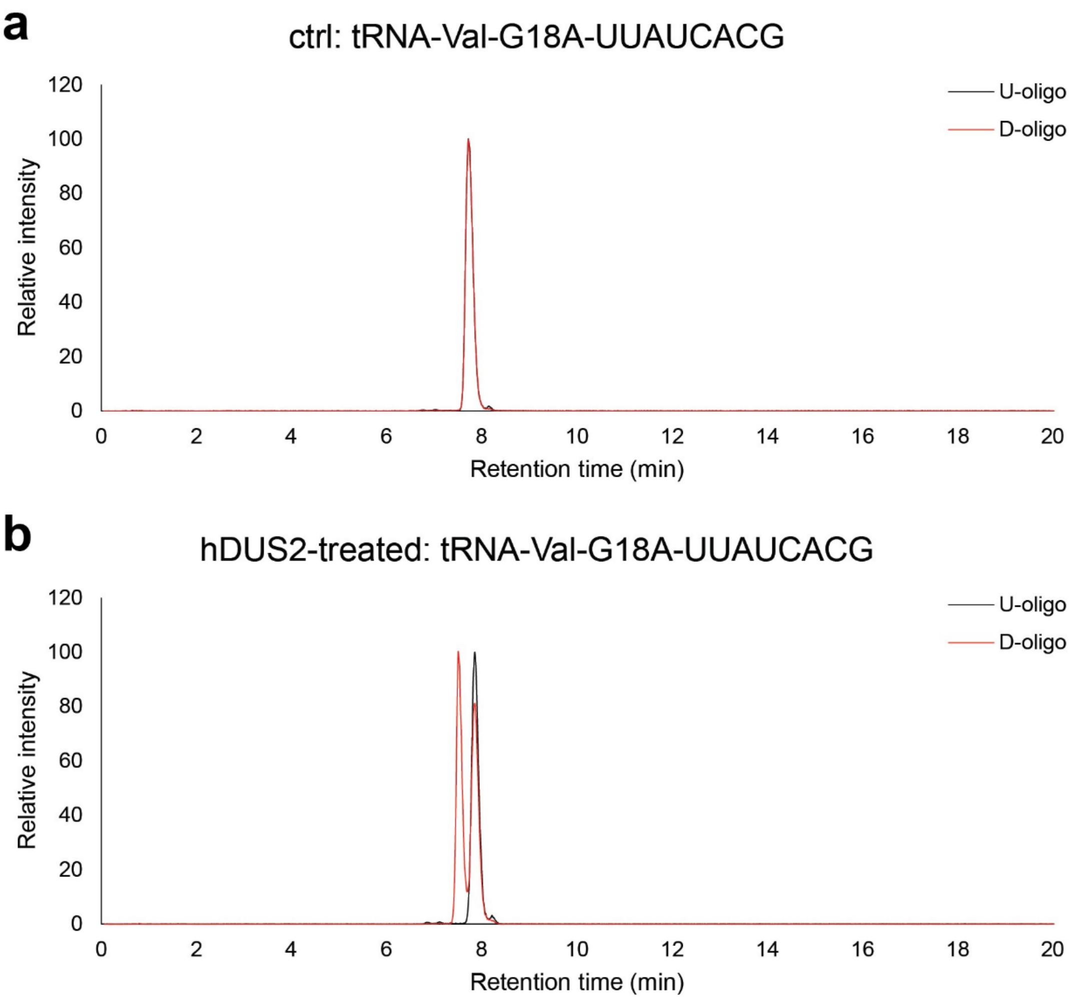


**Supplementary Figure 47.** Oligonucleotide LC-MS analysis of hDUS2-catalyzed dihydrouridine formation on tRNA-Val-CAC-G18A. Extracted ion chromatography of characteristic oligonucleotide fragment from RNase T1 digestion in control tRNA sample **(a)** and hDUS2-treated tRNA **(b)**. U-oligo: $[M-2H]^{2-}$ = 1273.668; D-oligo: $[M-2H]^{2-}$ = 1274.675.

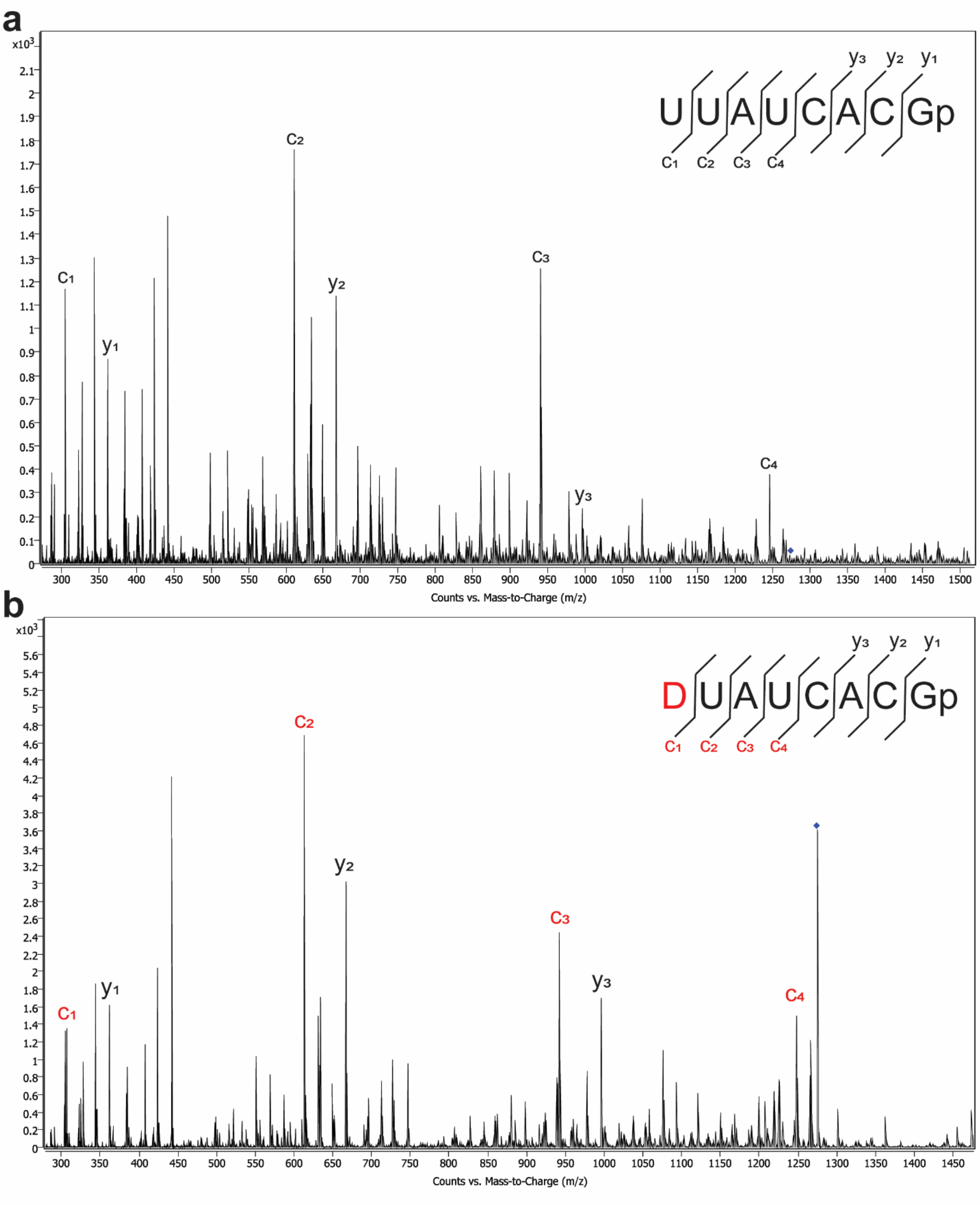


**Supplementary Figure 48.** MS/MS spectrum of UUAUCACGp and DUAUCACGp oligo fragments produced from RNase T1 digestion of control tRNA-Val-CAC-G18A sample **(a)** and hDUS2-treated tRNA-Val-CAC-G18A **(b)**. MS/MS fragmentation was performed by collision-induced dissociation. The U-containing precursor ion is $[M-2H]^{2-}$ = 1273.668. The D-containing precursor ion is $[M-2H]^{2-}$ = 1274.675.

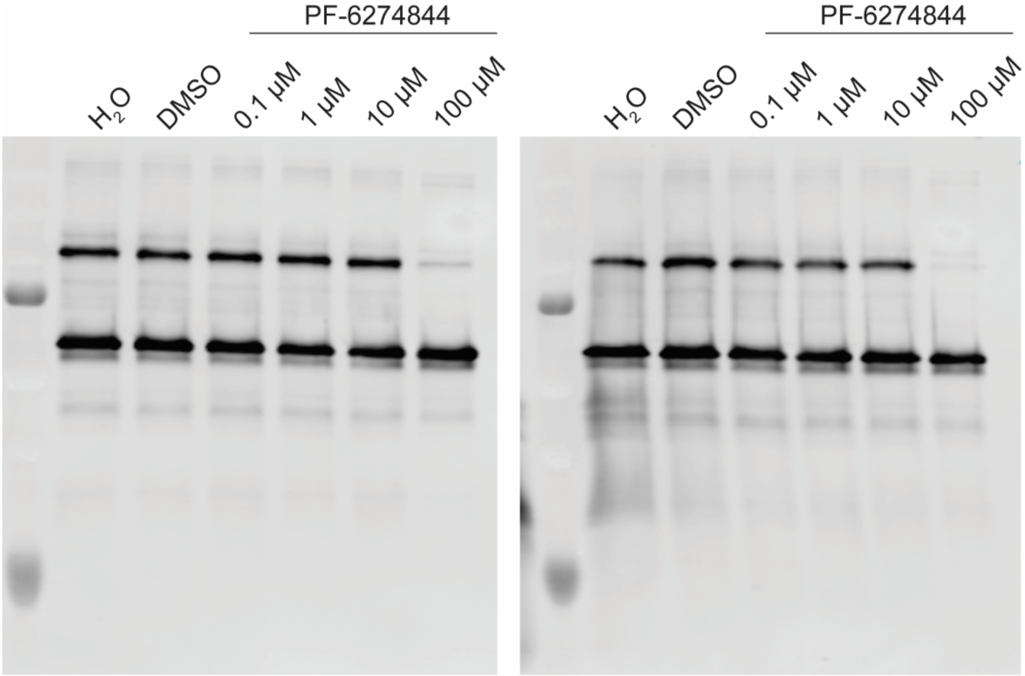


**Supplementary Figure 49.** Inhibition of crosslinking between hDUS2 and BrUrd-modified tRNA-Val-CAC by PF-6274844. 222 nM recombinant hDUS2 was pre-treated with PF-6274844 (0.111-111 µM) in a total volume of 18 µl of reaction buffer (20 mM µl Tris-HCl pH 8.0, 100 mM NaCl, 5 mM $MgCl_2$, 2 mM DTT, 10 mM NADPH) on ice for 30 min. Then 2 µl of refolded BrUrd-modified tRNA-Val-CAC was added to 1 µM final concentration. The mixture was incubated at 37 °C for 1 hour. Anti-hDUS2 Western blot was performed to characterize the inhibition of RNA-protein adduct formation.

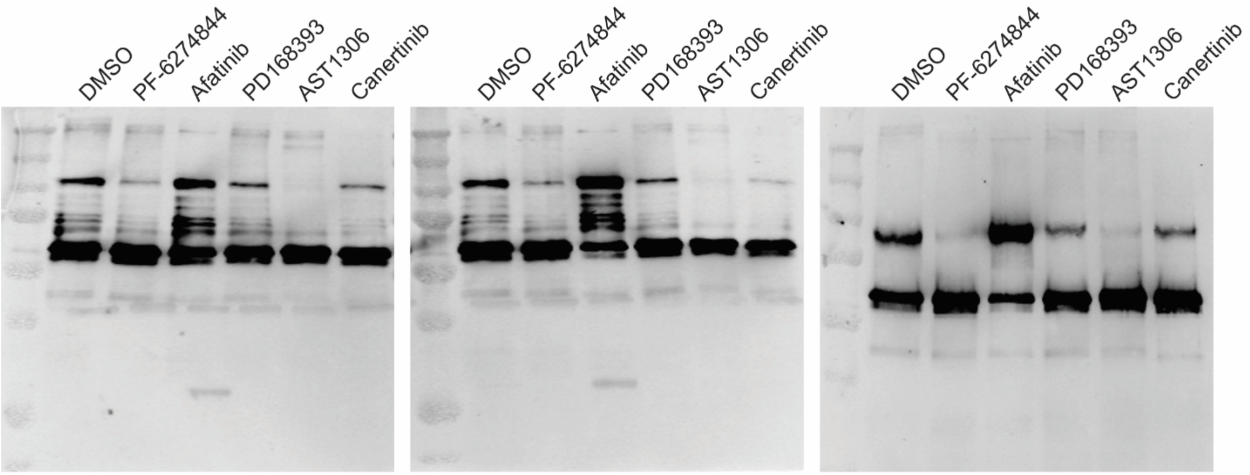


**Supplementary Figure 50.** Small molecule inhibition assay for 4-amino-quinazoline based EGFR inhibitors. Method is similar to that described in Supplementary Fig. 50. 222 nM recombinant hDUS2 was pre-treated with 111 µM PF-6274844 in a total volume of 18 µl of reaction buffer (20 mM µl Tris-HCl pH 8.0, 100 mM NaCl, 5 mM $MgCl_2$, 2 mM DTT, 10 mM NADPH) on ice for 30 min, followed by crosslink with BrU-tRNA-Val-CAC at 37 °C for 1 hour. Anti-hDUS2 western blot was performed to characterize the inhibition of RNA-protein adduct formation.

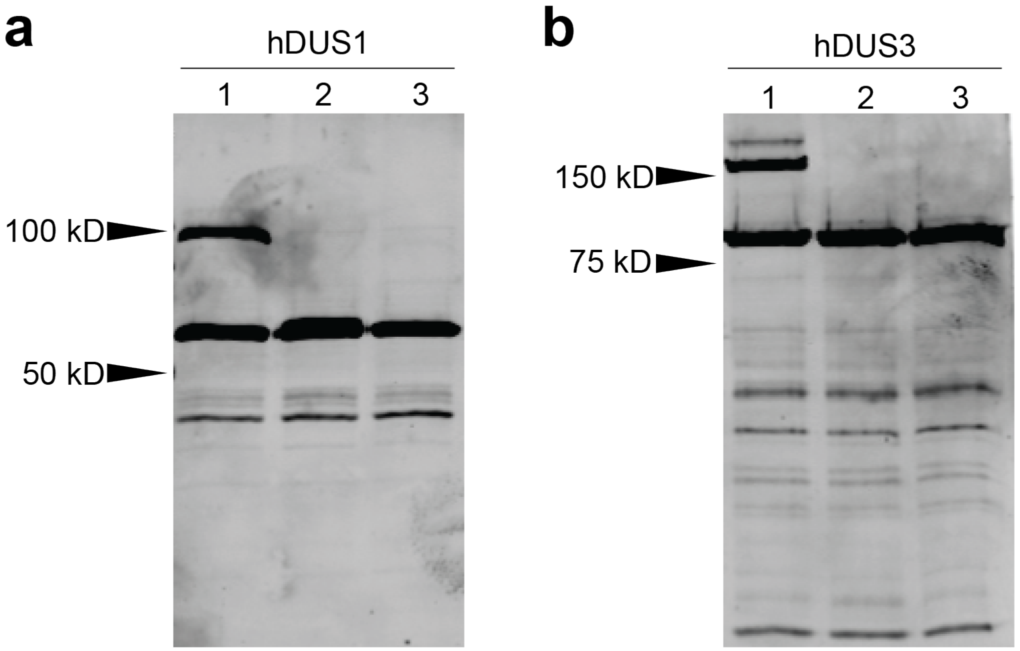


**Supplementary Figure 51.** *In vivo* crosslinking assay with BrUrd-modified tRNA-Val-CAC in cell lysate containing overexpressed DUS protein. Epitope-tagged hDUS1 **(a)** and hDUS3 **(b)** proteins were overexpressed in HEK293T cells by transient transfection with corresponding pcDNA5-3XFLAG plasmids. 10 µM tRNA was incubated with 45 µg of cell lysate (1mg/ml) and 1 ul of RNase Inhibitor, Murine (NEB, M0314S) in 50 µl at 37 °C for 1 hour. Anti-FLAG Western blot was used to detect product formation. 1, cell lysate with BrUrd-modified tRNA; 2, RNase digestion of 1; 3, cell lysate with unmodified tRNA.
